# Temporal mixture modelling of single-cell RNA-seq data resolves a CD4^+^ T cell fate bifurcation

**DOI:** 10.1101/074971

**Authors:** Tapio Lönnberg, Valentine Svensson, Kylie R James, Daniel Fernandez-Ruiz, Ismail Sebina, Ruddy Montandon, Megan S. F. Soon, Lily G Fogg, Michael J. T. Stubbington, Frederik Otzen Bagger, Max Zwiessele, Neil Lawrence, Fernando Souza-Fonseca-Guimaraes, William R. Heath, Oliver Billker, Oliver Stegle, Ashraful Haque, Sarah A. Teichmann

**Author notes:** denotes equal contribution (Tapio Lönnberg, Valentine Svensson, Kylie R James). denotes equal contribution (Oliver Stegle, Ashraful Haque, Sarah A. Teichmann).

## Abstract

Differentiation of naïve CD4^+^ T cells into functionally distinct T helper subsets is crucial for the orchestration of immune responses. Due to multiple levels of heterogeneity and multiple overlapping transcriptional programs in differentiating T cell populations, this process has remained a challenge for systematic dissection *in vivo*. By using single-cell RNA transcriptomics and computational modelling of temporal mixtures, we reconstructed the developmental trajectories of Th1 and Tfh cell populations during *Plasmodium* infection in mice at single-cell resolution. These cell fates emerged from a common, highly proliferative and metabolically active precursor. Moreover, by tracking clonality from T cell receptor sequences, we infer that ancestors derived from the same naïve CD4^+^ T cell can concurrently populate both Th1 and Tfh subsets. We further found that precursor T cells were coached towards a Th1 but not a Tfh fate by monocytes/macrophages. The integrated genomic and computational approach we describe is applicable for analysis of any cellular system characterized by differentiation towards multiple fates.

**One Sentence Summary:** Using single-cell RNA sequencing and a novel unsupervised computational approach, we resolve the developmental trajectories of two CD4^+^ T cell fates *in vivo*, and show that uncommitted T cells are externally influenced towards one fate by inflammatory monocytes.

## Introduction

T helper (Th) cells, also known as CD4^+^ T cells, are key instructors of the immune system (1). They display extensive functional and phenotypic diversity in response to a spectrum of immune challenges, including viral, bacterial, fungal and parasitic infection, immunogenic cancers, and autoimmune and allergic stimuli. Th cell subsets are distinguished from each other most frequently by the cytokines they secrete. Th1 cells produce interferon-γ, leading to macrophage activation and enhanced killing of intracellular pathogens. Th2 cells produce IL-4, IL-5, and IL-13, prompting eosinophils to act against extracellular parasites and venom. Th17 cells produce IL-17 and IL-22, promoting neutrophilic responses against extracellular bacteria and fungi. Follicular T helper (Tfh) cells, a more recently defined Th subset, secrete IL-21, and drive somatic hypermutation of immunoglobulin genes in germinal centre B cells. This produces high affinity antibodies, upon which many licensed vaccines depend for efficacy. Since Th subsets can both control infections and drive immune-mediated diseases there remains tremendous interest in the molecular mechanisms that control their *in vivo* development.

In order for Th cells to develop, CD4^+^ T cells must first be raised from an immunologically naïve state by antigenic stimulation of their highly diverse T cell receptors (TCR), which is followed by processes of clonal proliferation and differentiation. Recent *in vivo* data suggested that the unique TCR sequence of a single naïve CD4^+^ T cell imparts a genetically programmed preference towards a particular Th fate (2). However, co-stimulatory and cytokine signals can also profoundly influence both the magnitude of the response, and skewing towards particular Th fates. Several master transcription factors have been described in CD4^+^ T cells that drive and stabilize Th fates, which supports a view of Th development as a choice between clearly distinct states. However, the relationship between Th subsets, particularly between Tfh and other Th fates remains unclear *in vivo*.

In many cases, immune challenges, such as infection or vaccination, induce concurrent differentiation into two or more Th fates within the same individual. Indeed, by performing a limiting dilution single-cell adoptive transfer of naïve CD4^+^ T cells, it was suggested that daughter cells from a particular clone could bifurcate phenotypically to give rise to both Th1 and Tfh cells (2). However, it was not possible to visualize and pinpoint the bifurcation of Th1/Tfh cell fates *in vivo*.

Resolving Th cell fate decision-making *in vivo* using population-level approaches has been challenging, mainly due to extensive heterogeneity amongst differentiating cells. More specifically, CD4^+^ T cells at any given time point display a distribution of intermediate and transitional states, which blurs the dynamics of Th cell developmental progression (3). Tfh differentiation, in particular, has been difficult to elucidate since it involves multiple stages with potential overlap with transcriptional programs of other Th subsets. Of particular note, computational tools for modelling bifurcations in cellular decision-making have not been available.

Th cell fate decisions are driven by both intrinsic factors and external signalling cues from other cells. Conventional dendritic cells (cDCs) are important cellular sources of antigenic stimulation, co-stimulation and cytokines for Th differentiation in secondary lymphoid tissues. Intra-vital imaging in lymph nodes has demonstrated that cDCs make long-lasting stable contacts with naïve CD4^+^ T cells in order to initiate T cell priming (4). Once activated, CD4^+^ T cells continue to require antigenic stimulation via their TCR to optimize their proliferation and Th differentiation (5–7). Continued signalling has been reported to be important for Th1 responses, although the cell types providing this signal remain unknown (4). A recent report suggested that CXCR3 expression by activated CD4^+^ T cells facilitated continued interaction with adoptively-transferred CXCL9 and CXCL10-expressing cDCs (8), however, interactions with endogenous myeloid cell populations, including cDC subsets and monocytes have not been studied *in vivo*. While Tfh cells are sustained, once generated, via multiple molecular interactions with B cells in developing germinal centres (9, 10), possible roles for myeloid cells in providing early instruction towards a Tfh fate remain relatively unexplored. A recent study targeted antigens to two different cDC-subsets *in vivo,* and suggested that CD8α^−^ cDCs displayed the greater propensity for generating Tfh responses (11). Whether Th1/Tfh fate bifurcation can result from differential interactions with cDC subsets or activated monocytes currently remains unknown.

Herein we have used single-cell RNA sequencing (scRNA-seq) to study the various transcriptional states of individual CD4^+^ T cells during blood-stage *Plasmodium chabaudi* infection in mice. This is an experimental model of malaria in which CD4^+^ T cells are essential for controlling parasite numbers, and which is characterized by concurrent development of Th1 and Tfh cells (12). We have used *Plasmodium-*specific TCR transgenic CD4^+^ T (PbTII) cells to minimise the effects of TCR diversity on Th fate decisions.

Crucially, our approach builds on scRNA-seq profiling coupled with new computational strategies to reconstruct the differentiation trajectories of Th1 and Tfh cells at a single-cell resolution. Our data reveals, for the first time, the molecular detail of how a single antigen-specific CD4^+^ T cell clone can undergo parallel development into Th1 and Tfh states *in vivo*, and reveals the hierarchical regulation of genes involved in this cell fate decision. Finally, we investigated intercellular interactions using scRNA-seq, and predicted roles for inflammatory monocytes, after cDC-dependent T cell activation, in coaching uncommitted CD4^+^ T cells, specifically towards a Th1 fate.

## Results

### scRNA-seq resolves Th1 and Tfh cell fates during *Plasmodium* infection in mice

To study concurrent progression towards Th1 and Tfh fates, and to characterize the heterogeneity associated with this process during an *in vivo* CD4^+^ T cell response, we performed scRNA-seq of PbTII cells during *Pc*AS infection (Figure 1A, Figure S1). We transferred naïve, proliferative dye-labeled PbTII cells into congenically marked wild-type mice, and recovered them at days 2, 3, 4, and 7 post-infection (p.i.) by fluorescence-activated cell sorting (FACS) of those expressing the early activation marker, CD69, or displaying dilution of the proliferative dye (Figure S2). Flow cytometric measurements of the canonical Th1 markers, T-bet (coded by *Tbx21*) and Interferon-γ, and Tfh markers, CXCR5 and Bcl6, indicated that these subsets emerged in parallel by day 7 p.i. (13, 14) (Figure 1B-D). Notably, markers of Th2, Th17 or Treg subsets were not upregulated on the PbTII cells (Figure S3).

**Figure 1.**
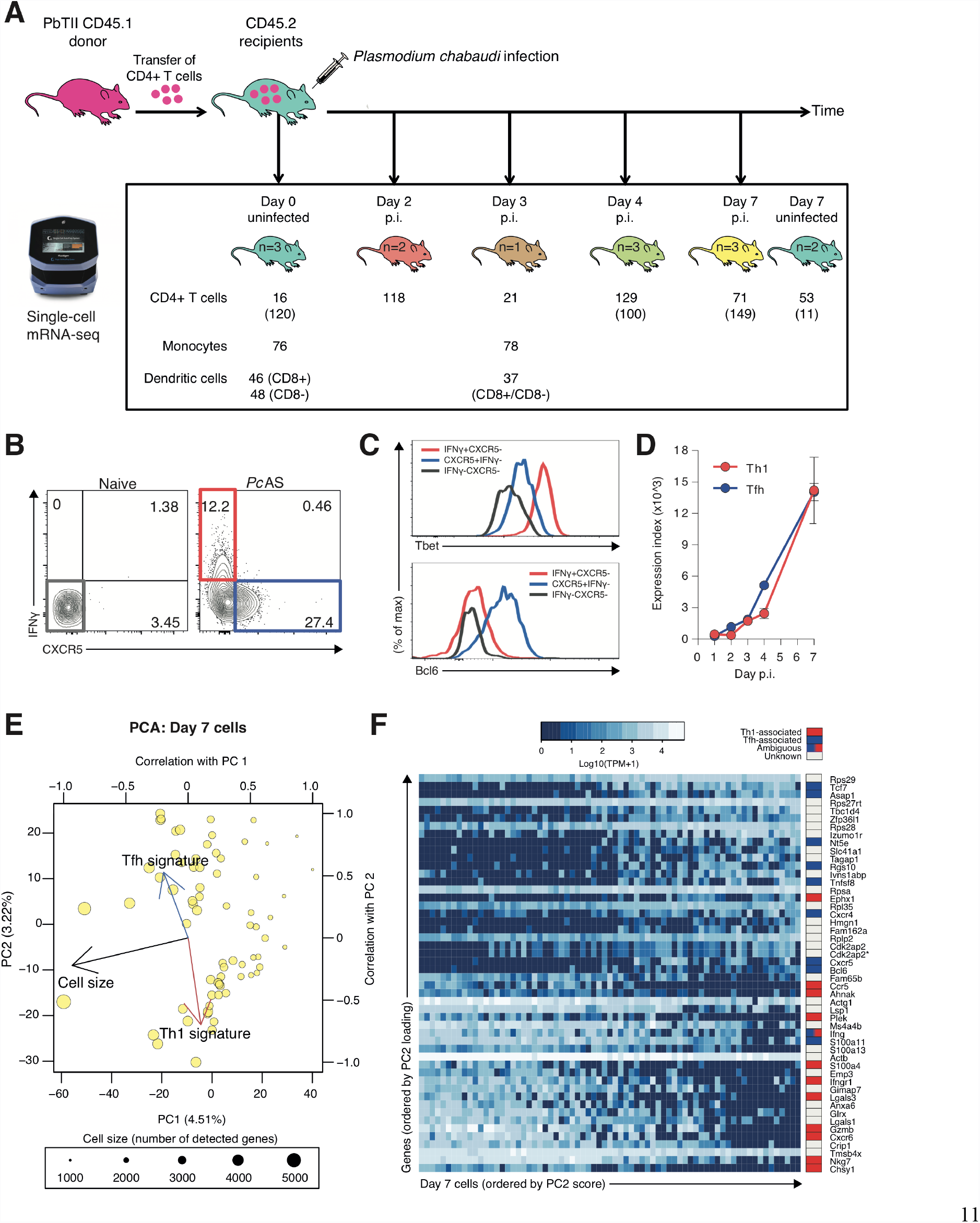
Single-cell mRNA-sequencing of activated antigen-specific PbTII CD4^+^ T cells. (**A**) Experimental setup. PbTII cells were transferred from a single donor to multiple recipients. “n” refers to the number of recipient mice per time point. Also shown are the numbers of single cells from which high-quality mRNA-seq data was successfully recorded. The numbers in parentheses refer to the experiment presented in Fig. S12. p.i., post-infection. (**B-C**) Representative FACS plots showing bifurcation of splenic Th1 (IFNγ^+^T-bet^+^) and Tfh (CXCR5^+^Bcl6^+^) PbTII CD4^+^ T cells at day 7 post-infection with *Pc*AS. (**D**) Flow cytometry data indicate concurrent differentiation of Th1 (IFNγ^+^) and Tfh (CXCR5^+^) PbTII CD4^+^ T cells within the spleen of *Pc*AS-infected mice (n=4). Index expression is the product of MFIand proportion IFNγ^+^ or CXCR5^+^. These data are representative of two independent experiments. MFI, mean fluorescence intensity. (**E**) PCA of single PbTII cells at 7 days post-infection with *Pc*AS. The arrows represent the Pearson correlation with PC1 and PC2. Cell size refers to the number of detected genes. The size of the data points also represents cell size. “Th1 signature” and “Tfh signature” refer to cumulative expression of genes associated with Th1 or Tfh phenotypes (15). PC, Principal Component. **(F)** Expression of top 50 genes with largest PC2 loadings of day 7 cells (D). The genes were annotated as Th1-or Tfh-associated based on public datasets (15, 38, 46, 47). \**Cdk2ap2* appears twice because two alternative genomic annotations exist. PC, Principal Component

We initially used Principal component analysis (PCA) to assess the overall heterogeneity of the PbTII cells (Figure 1E, Figure S4A). In all time points, the first principal component was strongly associated with the number of detected transcripts, which is reflective of changes in cellular RNA content and, in general, is linked to proliferative status (Figure S4B). As expected, the variability related to previously established Th1 and Tfh gene expression signatures became more prominent with the progression of time (15) (Figure S4C). Notably, at day 7 p.i., a PCA using these signature genes alone recapitulated the results of the genome-wide PCA (Spearman correlation −0.87) (Figure S5). Amongst the cells from day 7 p.i., two distinct subpopulations were apparent, separated along PC2 (Figure 1E). Notably, many of the genes associated with these subpopulations have been identified as associated with either Th1 or Tfh fates (Figure 1F, Table S1). Results from a global PCA of the entire dataset were largely in accordance with the time point information, with the Th1/Tfh signature genes showing separation along multiple PCs (Figure S6). Taken together, these results suggested a progressive commitment to Th1 and Tfh fates, and indicated that single-cell transcriptomes could be used for estimating both proliferative states and degrees of differentiation of individual cells.

### Unbiased delineation of Th1 and Tfh trajectories using a Mixture of Gaussian Processes model

The results from the PCA suggest that gene expression variation in PbTII single-cell transcriptomes permit reconstruction of the transcriptional programs underlying Th1 and Tfh differentiation. To more explicitly model the temporal dynamics of the differentiation process, we developed and applied GPfates, a temporal mixture model that builds on the Gaussian Process Latent Variable Model (GPLVM) and Overlapping Mixtures of Gaussian Processes (OMGP) (16). This approach first reconstructs the differentiation trajectory from the observed data (“pseudotime”, Figure 2A-B), thereby establishing an order for the cells. While our model uses the sample time as prior information, the inferred temporal orderings did not strictly adhere to these experimental time points (Figure S7). For example, cells from day 4 p.i. were mixed with some of the cells from day 3 and day 7 at either end of the day 4 pseudotime distribution. This was consistent with the idea that bulk assessments of cells at specific time points fail to take into account the heterogeneity and differential kinetics of responses made by single cells. We also repeated this analysis without supplying the experimental sampling times to the model, finding overall consistent results (Comp. Supp. Figure 8).

**Figure 2.**
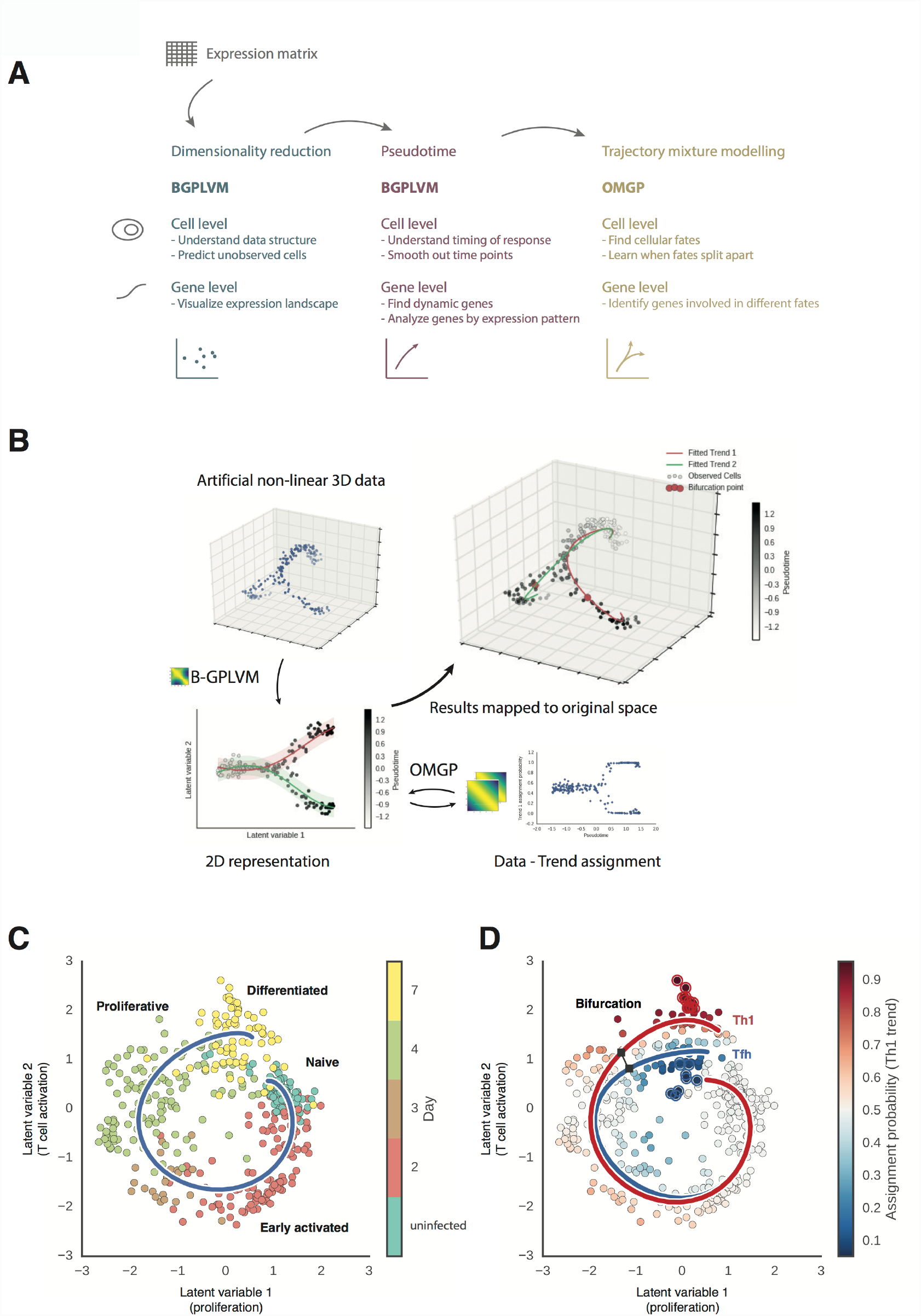
GPfates modelling of bifurcation processes using scRNA-seq data. **(A)** Overview of analysis abilities from the framework of Gaussian Processes. Data is modelled and interpreted on the cellular level using the global genomic level data. Through downstream analysis from these models, it is possible to investigate individual genes to explain the drivers of the different models. **(B)** Sketch of the analysis workflow. A low-dimensional model of the non-linear high-dimensional data is inferred by Bayesian GPLVM. The low-dimensional representation is then modelled as an Overlapping Mixture of Gaussian Processes. This gives us a data-trend assignment per cell which can be used for interpretation. Since the models are all predictive, the low-dimensional model can be interpreted in the original high-dimensional space. **(C)** The low-dimensional representation of our data. The blue line depicts the progression of pseudotime. The text labels illustrate features of typical cells on that region of the pseudotime, and are provided purely as a visual aid.

In a second step, GPfates uses a time series mixture model, which we adapted from a model that was initially developed to deconvolve temporal data into independent separate trends, and which is related to previous time series models for bulk gene expression time series (16). Using this approach, we identified two simultaneous trends (Figure 2C-D). These two alternative trajectories were in agreement with the Th1/Tfh signature genes identified by Hale et al. (15) (Figure 3A-D), indicating that the fitted mixture components correspond to cells with Th1 and Tfh phenotypes. Notably, these trends could not be identified by other published methods for reconstructing single-cell trajectories (17, 18) (Figure S8). Furthermore, the mixture modelling in GPfates could also successfully resolve bifurcation events in two other recently published scRNAseq datasets, which examined lung epithelial development in mice (Comp. Supp. Figure 11) (19) and primordial germ cell development in human embryos (Comp. Supp. Figure 12) (20). This suggests that pseudotime inference coupled with time series mixture modelling is applicable more generally for studying cellular differentiation in scRNAseq data.

**Figure 3.**
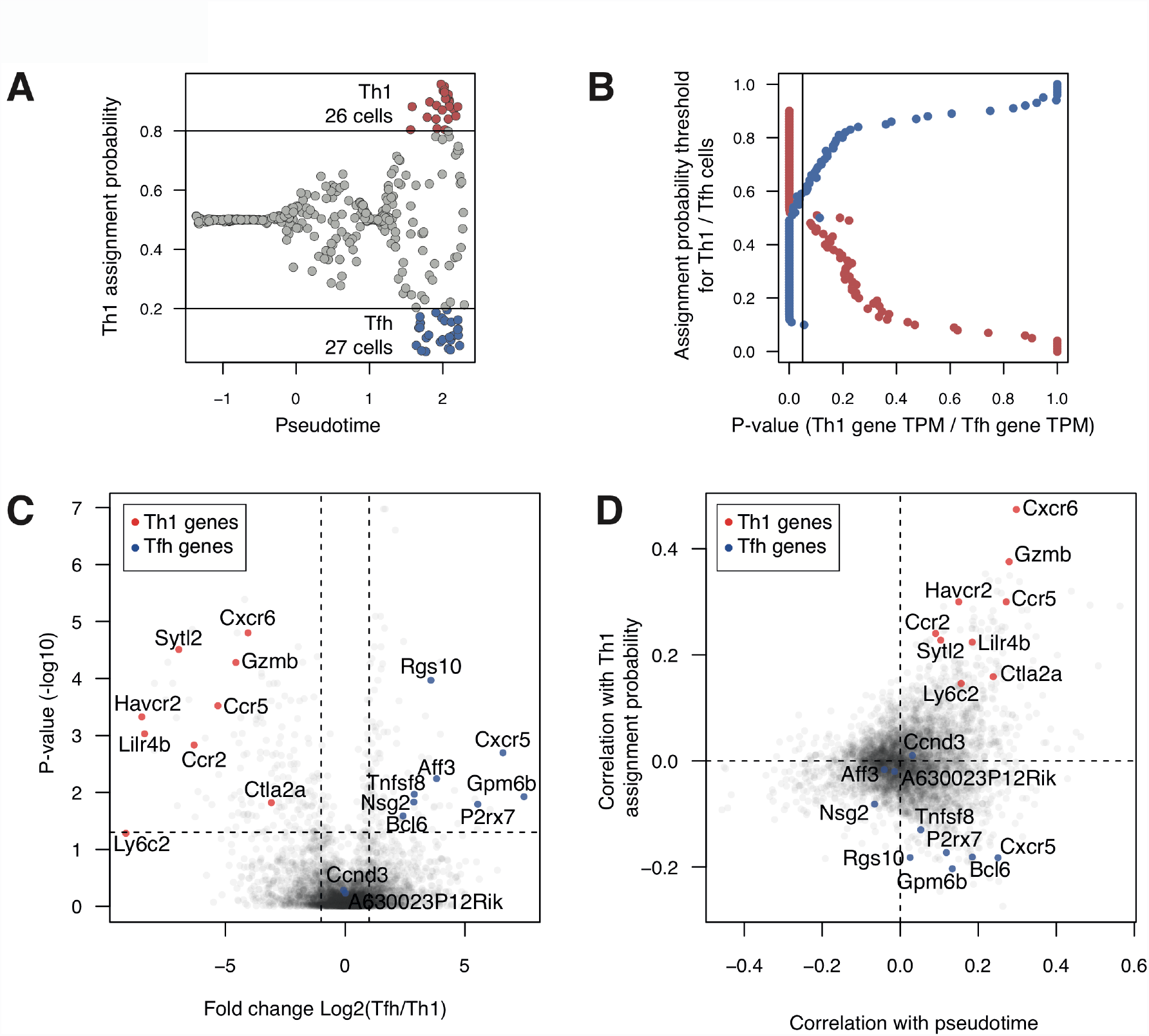
The relationship of known Th1-and Tfh-transcriptomics signatures and the trajectories determined using the GPfates analysis. (**A**) Th1 and Tfh states were defined as cells with assignment probability of ≥0.8 for the respective trend. For each single cell, cumulative expression of Th1 and Tfh signature genes (15) was calculated as in Figure 1E. (**B**) The effect of the probability threshold on the cumulative expression of Th1 and Tfh signature genes. The p-values were calculated using Wilcoxon rank sum test. (**C**) The expression of Th1 and Tfh signature genes in Th1 and Tfh cells defined using the GPfates model (A). For all genes expressed by at least 20% of the single cells, fold changes were calculated. The p-values were calculated using Wilcoxon rank sum test and adjusted for multiple testing using Benjamini & Hochberg correction. (**D**) Correlation of expression of Th1 and Tfh signature genes with pseudotime and with Th1 assignment probability.

Next, we sought to more clearly characterize the bifurcation time point. Using a change point model to annotate the inferred trajectories (see section 4.2 of the Computational Supplement), we could divide pseudotime into *before* and *after* bifurcation. We sought to characterize single cells that existed at the Th1/Tfh bifurcation point. Firstly, bifurcation initiated amongst cells from day 4 p.i. (see section 6.2 of computational supplement for a robustness analysis), specifically at a relatively early point in pseudotime compared with all day 4 p.i. cells (Figure 4A). Bifurcating PbTII cells also expressed the largest number of genes compared to those at all other points in pseudotime.

**Figure 4.**
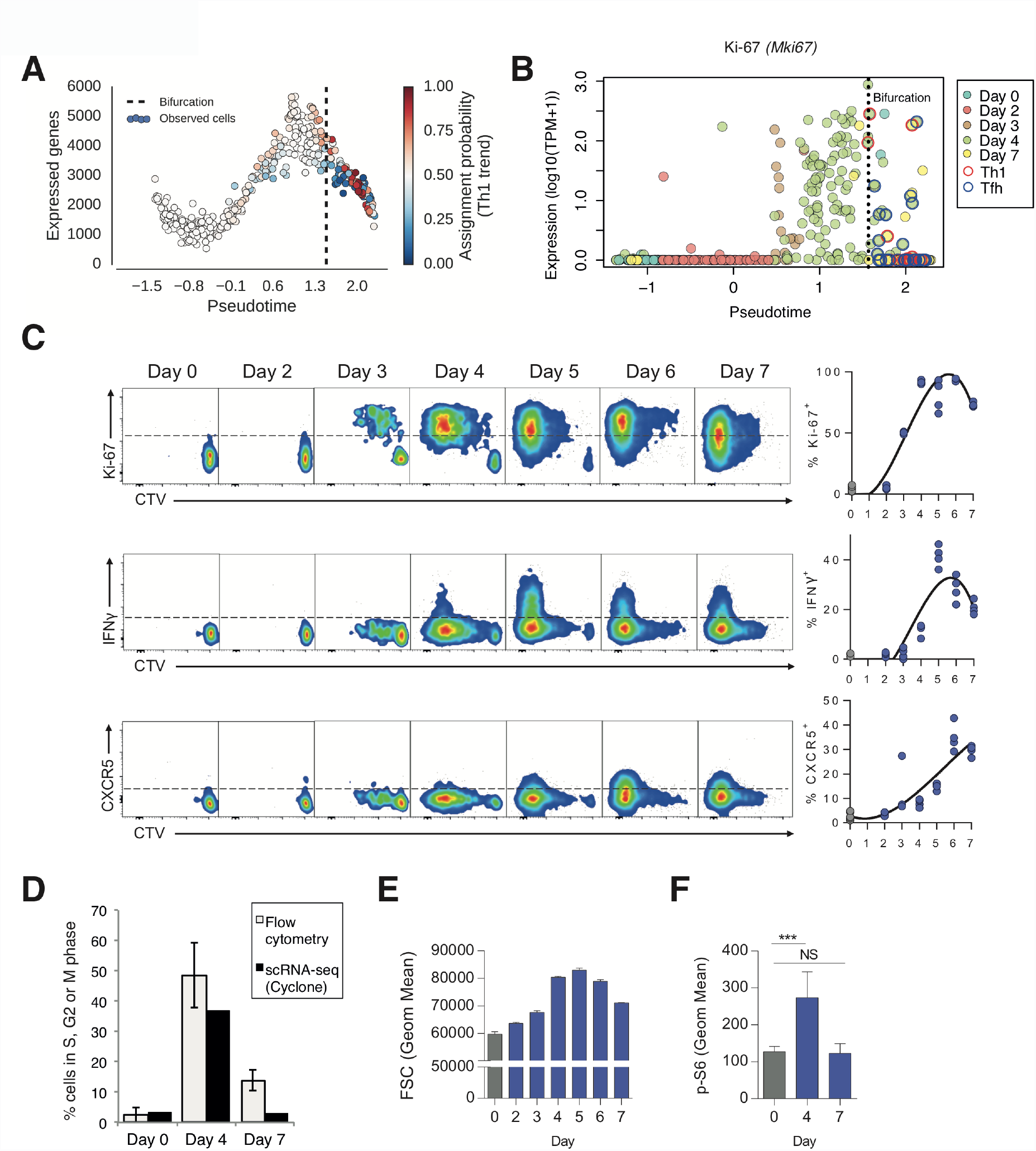
The bifurcation of T cell fates is accompanied by changes in transcription, proliferation and metabolism. (**A**) The relationship of Th1-Tfh bifurcation point and the number of detected genes per cell. (**B**) The expression level of the proliferative marker Ki-67 (encoded by the *Mki67* gene) across pseudotime. (**C**) Representative FACS plots showing kinetics of CellTrace^™^ Violet (CTV) dilution and Ki67, IFNγ or CXCR5 expression, with summary graphs showing % of PbTII cells expressing these (after 10^6^ PbTII cells transferred) in uninfected (Day 0) and *PcAS*-infected mice at indicated days post-infection (n=4 mice/timepoint, with individual mouse data shown in summary graphs; solid line in summary graphs indicates results from third order polynominal regression analysis.) Data are representative of two independent experiments. (**D**) Experimental and computational analysis of cell cycle speed of PbTII CD4^+^ T cells activated in response to *Pc*AS. The allocation of cells to cell cycle phases was performed by flow cytometry using Hoechst staining (Figure S9B) and computationally using the Cyclone algorithm (22). The relative cell-cycle speed was determined by measuring the fraction of cells in S, G2, or M phases. (**E**) Cell size estimation using FSC (Forward Scatter) measurements of PbTII cells. (**F**) Cellular metabolic activity of PbTII cells in naive mice (n=3) and at days 4 and 7 post-infection (n=6) as determined by flow cytometric assessment of ribosomal protein S6 phosphorylation (p-S6). Histogram and proportions are representative of two independent experiments. Statistics are one-way ANOVA and Tukey's multiple comparisons tests ***p<0.001.

High transcriptional activity correlated with upregulation of *Mki67* and other known proliferation marker genes (21) (confirmed at the level of Ki-67, Figure 4B-C and S9A). It also correlated with cell cycle activity, based on computational allocation of cells into cell cycle stages (22), and flow cytometric confirmation of DNA content and cell size (Figure 4D-E). Bifurcating PbTII cells also had increased expression of genes associated with aerobic glycolysis (data not shown), an indication of increased metabolic requirements being met by glucose metabolism and increased mTORC1 activity. Consistent with this was the observed elevated levels of ribosomal protein S6 phosphorylation by day 4 p.i. (Figure 4F).

Taken together, our data indicate that bifurcating PbTII cells exhibit a highly proliferative and metabolically active state, coupled with the upregulation of thousands of genes. Importantly, progression from the Th1/Tfh bifurcation point to either fate was marked by widespread silencing of gene expression across the genome. Although this decrease in gene expression can be partially explained by a deceleration in cell cycle speed, it is also consistent with other cellular differentiation processes characterized at a single-cell resolution (19).

### Detectable expression of endogenous T cell receptor loci reveals breadth of clonotype fates

Since previous reports have suggested a role for TCR sequences in determining Th cell fate (2), our TCR transgenic approach was designed to minimize this potential source of variability. Importantly however, PbTII cells were generated in mice with functional *Rag1* and *Rag2* genes, and therefore, retained natural expression of highly diverse endogenous TCR chains in addition to the transgenic TCR. Sequence analysis of TCR transcripts in single PbTII cells confirmed universal expression of the PbTII Vα2 and Vβ12 chains in all cells (Supplementary Tables 2 & 3). Moreover, it confirmed highly diverse, though lower levels of expression of endogenous TCRα chains in many cells (Figure S10).

Given the vast combinatorial diversity of endogenous TCR sequences, we employed these as unique molecular barcodes to scan for PbTII cells that could be inferred with high confidence to have derived from a single common PbTII progenitor clone. Notably, we identified six clones comprising two or more sibling cells, while all other PbTII cells were individually unique. Of these six clones, two consisted of sibling cells that mapped close to the bifurcation point. For the remaining four clones, siblings exhibited highly diverging patterns of gene expression, with three sibling groups falling at the extremities of the Th1-Tfh phenotype spectrum (Figure 5A). These results demonstrate that during an *in vivo* infection, the progeny of a single CD4^+^ T cell clone can differentiate into both Th1 and Tfh cells.

**Figure 5.**
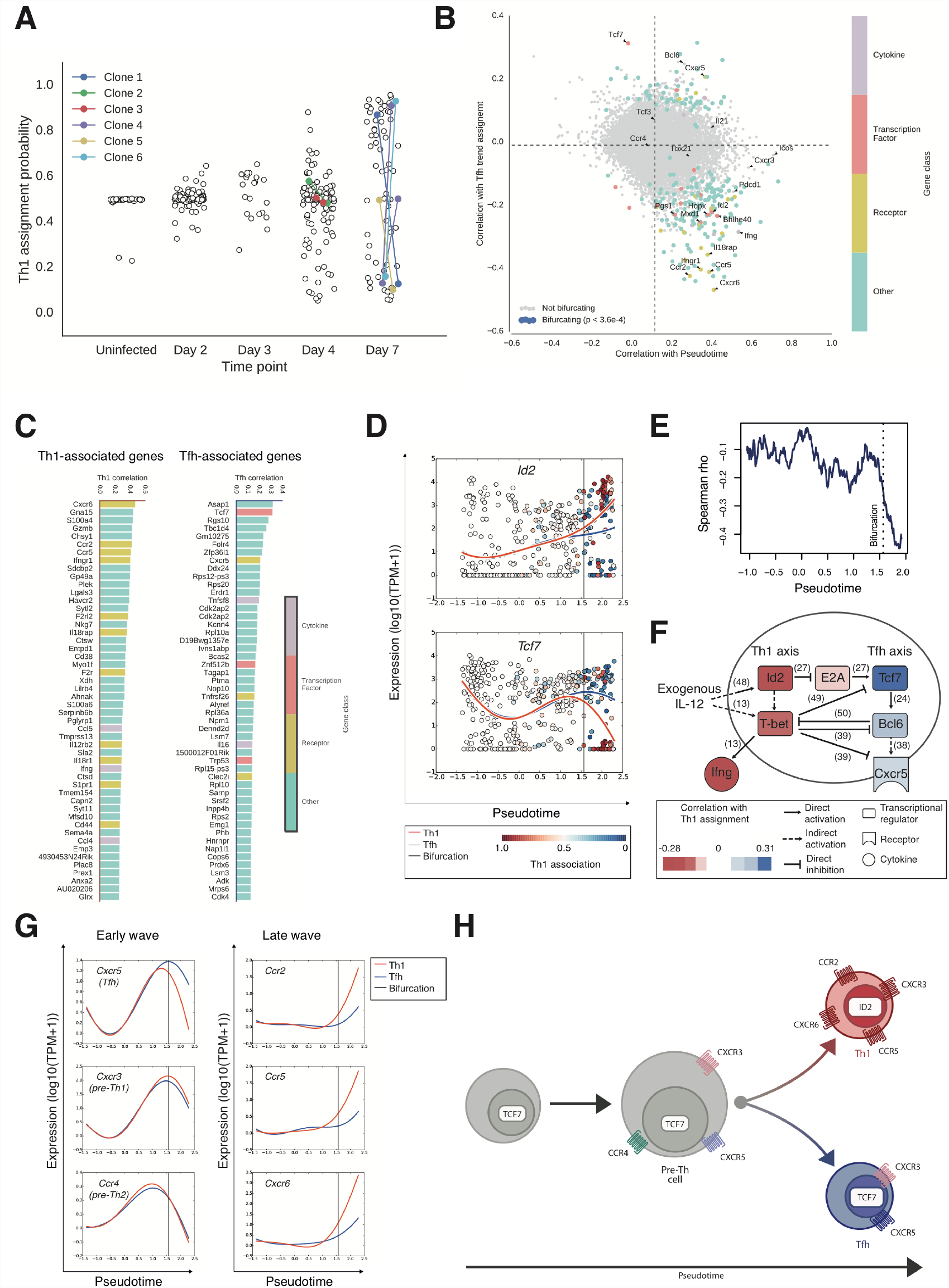
Mechanisms underlying the differentiation of Th1 and Tfh cells. (**A**) Parallel Th1 and Tfh differentiation within cells of a single CD4^+^ T cell clone. The colours represent clones determined by sequence analysis of secondary T cell receptor genes (Supplementary Tables 2 and 3). (**B**) Identification of genes associated with the differentiation of Th1 or Tfh cells. For every gene, the correlation of its expression with pseudotime (x-axis) and Tfh trend assignment (y-axis) are shown. Statistical significance was determined using the bifurcating score (methods). Genes satisfying the significance threshold of FDR<0.002 are represented in colours according to the functional classification of the genes (methods and Supplementary Table 4). FDR, False Discovery Rate, estimated by performing the same analysis with permuted data. (**C**) The genes with strongest association with Th1 (left) or Tfh differentiation (right). The genes were filtered using the bifurcation score as in (B). The genes were then ranked in descending order of association with either Th1 or Tfh trend. *Cdk2ap2* appears twice because two alternative genomic annotations exist. (**D**) The expression of *Id2* (upper panel) and *Tcf7* (lower panel) across the pseudotime. The curves represent the Th1 (red) and Tfh (blue) trends when weighing the information from data points according to trend assignment. The colour of the data points represents the strength of the relationship with the two alternative trends. (**E**) The correlation of *Id2* and *Tcf7* expression at single-cell level. Using a rolling window method, Spearman rho was calculated in windows of 100 cells. The pseudotime values are mean values within each window. (**F**) A model depicting known interactions of *Id2* and *Tcf7*. The colours represent the Pearson correlation of gene expression and the Th1 trend assignment in single cells. The numbers in parentheses refer to the original publications (13, 24, 27, 38, 39, 48–50). (**G**) The expression kinetics of the chemokine receptor genes *Cxcr5, Cxcr3, Ccr4, Ccr2, Ccr5,* and *Cxcr6* across pseudotime. The curves represent the expression patterns associated with the Th1 (red) and the Tfh (blue) trends. (**H**) A model summarizing the expression patterns of *Id2*, *Tcf7*, and the chemokine receptors during Th1-Tfh cell fate determination. The size of the cell represents proliferative capacity (Fig. 4 (B-E). The colour of receptors and transcription factors represent differences in expression level.

### Transcriptional signatures associated with bifurcation of Th1 and Tfh fates

Next, we sought to identify genes whose expression followed the pattern of branching. We derived *bifurcation statistic* to estimate the concordance with the bifurcation for individual genes (see section 4.2 of the Computational Supplement text for details, Figure 5B). Among the highest-ranking bifurcating genes, the most common pattern was an increase in expression during progression to the Th1 fate. These genes were positively correlated with both pseudotime and the Th1 trend assignment (Figure 5B). This suggests that Tfh cells are in fact developmentally closer to the highly proliferative progenitor state than Th1 cells as the Th1 fate involves up-regulation of numerous genes not expressed in either the progenitor or Tfh states.

The highest-ranking transcription factors were *Tcf7* for the Tfh fate, and *Mxd1, Bhlhe40, Hopx, Pgs1* and *Id2* for the Th1 fate (Figure 5C). In addition, the hallmark Tfh transcription factor *Bcl6* was strongly associated with the Tfh fate. *Tcf7* is required for T cell development, and has been recently shown to be instrumental for Tfh differentiation (23, 24). Notably, it represented one of the rare genes defined by a decrease in expression when moving towards the Th1 fate. Of the Th1-associated transcription factors, *Mxd1* is a negative regulator of the proliferation-associated, proto-oncogene, *Myc* (25) and *Bhlhe40* has been recently identified as a cofactor of *T-bet* (coded by *Tbx21*) (26). *Id2* is known as an antagonist of *Tcf7* (27) and as a regulator of effector CD8^+^ T cell responses. Notably, while this manuscript was under revision, the role of Id2 as a key driver of Th1 responses was independently shown by another study (28).

Our results strongly support reciprocal regulation of *Id2* and *Tcf7* as a key feature of the Th1/Tfh bifurcation process. Expression of *Id2* and *Ifng* were highly correlated in the later stages of Th1 differentiation, and negatively correlated with *Tcf7,* both at a transcriptional and protein level (Figure 5D-F, Figure S11). Notably, the hallmark Th1 transcription factor *Tbx21* was induced before the bifurcation point, and showed only modest separation after bifurcation (Figure S12).

To validate the robustness of these gene signatures and the timing of the bifurcation, we repeated the infection, and at days 0, 4 and 7 sequenced additional single PbTII-cells using the Smart-seq2 protocol (29) (Figure 1A & S13A). Consistent with the original data, the cells from day 7 (but not day 4) segregated into two subpopulations correlating with Th1 and Tfh gene signatures (Figure S13A). Subset-characteristic co-expression patterns of the bifurcating genes identified by GPfates emerged by day 7 (Figure S13B). Notably, at this time, the cells from the different mice could be equally separated into distinct Th1-and Tfh-subpopulations using the top bifurcating genes (Figure S13C). Taken together, this indicated that the gene expression patterns associated with the cell fate bifurcation were reproducible across experiments and sequencing platforms.

In Th1 cells, a large fraction of the bifurcating genes were cytokine and chemokine receptors, including the top-ranked gene, *Cxcr6,* confirmed at protein level (Fig S14A and S14B), other established Th1 markers, *Ifngr1* and *Il18rap* (30, 31), and the chemokine receptors *Ccr2* and *Ccr5* (Figure 5C). These data were consistent with the idea that Th1 cells can migrate to peripheral tissues and remain receptive to external signals. In contrast, the only bifurcating chemokine receptor associated with a Tfh fate was *Cxcr5*, a gene established to mediate migration of Tfh cells into B cell follicles (32, 33).

*Cxcr5* was among an early wave of chemokine receptor genes, including *Cxcr3* and *Ccr4* (Figure 5G) whose expression and translation into protein (Figure S14C) was initiated before the Th1/Tfh bifurcation point had been reached. We hypothesized that differences in the timing of expression of receptors reflected their roles in controlling differentiation or effector function. We reasoned, for instance, that while *Cxcr6*, *Ccr2* and *Ccr5* served to mediate trafficking and effector function of Th1 cells, others such as *Cxcr3* and *Cxcr5* controlled Th cell fate via interactions with other immune cells (Figure 5H). Indeed, *Cxcr5* allows T cell trafficking towards B cells (34, 35), while *Cxcr3* has been associated with cDC-driven Th1 fates (8).

### Myeloid cells support a Th1 but not Tfh fate

After activation and proliferation, PbTII cells reached an uncommitted state around the bifurcation and expressed chemokine receptors that indicated receptiveness to other chemokine-expressing cells. Given that B cells were essential, as expected, for supporting a Tfh fate in PbTII cells (Fig S15), we hypothesized that myeloid cells provided alternative, competing signals to promote a Th1 fate.

To study this, we performed scRNA-seq on splenic cDCs and inflammatory monocytes when activated PbTII cells were yet to bifurcate. We sorted CD8α^+^ and CD11b^+^ cDCs and Ly6C^hi^ monocytes from naïve and infected mice (Figure S16) and subjected these to single-cell analysis. PCA of cDCs firstly distinguished between the two naïve cell types, separating them along PC2 (Figure 6A & S17) with an efficiency consistent with recent data (36), and further highlighting a number of expected and previously unknown cDC subset-specific genes (Figure S18A-C). We next compared naïve cDCs with those from infection (Figure 6A & S16), and separated these along PC6 (Figure 6A). Analysis of differential gene expression between cDCs from naive and infected mice identified 30 genes, 29 upregulated (Figure 6B & S19), including interferon-associated transcription factors, *Stat1* and *Irf1*, and CXCR3-attractant chemokine genes, *Cxcl9* and *Cxcl10*. Notably, gene expression patterns amongst individual cDCs varied according to the gene. For example, *Stat1* and *Irf1* were heterogeneously expressed amongst individual naïve cDCs, and further upregulated during infection (Figure 6C). This was similar for *Cxcl9,* which was expressed by CD8α^+^ cDCs in naive mice, while *Cxcl10* was induced only upon infection (Figure 6C). These data revealed interferon-associated gene expression amongst individual cDCs, and also suggested interactions between cDCs and uncommitted CXCR3^+^ PbTII cells, consistent with a recent study (8). Next, PCA of Ly6C^hi^ monocytes from naïve and infected mice distinguished them from each other along PC2 (Figure 6D & S20). Differential gene expression analysis between naïve and infected groups uncovered ~100 genes, both up-and down-regulated during infection (Figure 6E & S21). This illustrated a fundamental difference in the directionality of transcriptional changes in individual monocytes compared to cDCs during *Plasmodium* infection, with only monocytes exhibiting down-regulation of gene expression (Figure 6B-C & E-F). Interestingly, a high proportion (~40%) of genes upregulated in cDCs were also induced in Ly6C^hi^ monocytes, including transcription factors *Stat1* and *Irf1*, and the chemokine *Cxcl10* (Figure 6E & F), suggesting possible overlapping biological functions between these cell types. In addition, monocyte-specific chemokines were also observed, including *Cxcl2, Ccl2* and *Ccl3* (Figure 6E & F). Furthermore, specific examination of all immune cellular interaction genes (Figure S22) revealed emerging variable expression of *Tnf, Cd40, Pdl1*, *Ccl4, Ccl5, Cxcl16, Cxcl9, and Cxcl11* in monocytes, thus suggesting complex interactions and multiple roles for Ly6C^hi^ monocytes during infection.

**Figure 6.**
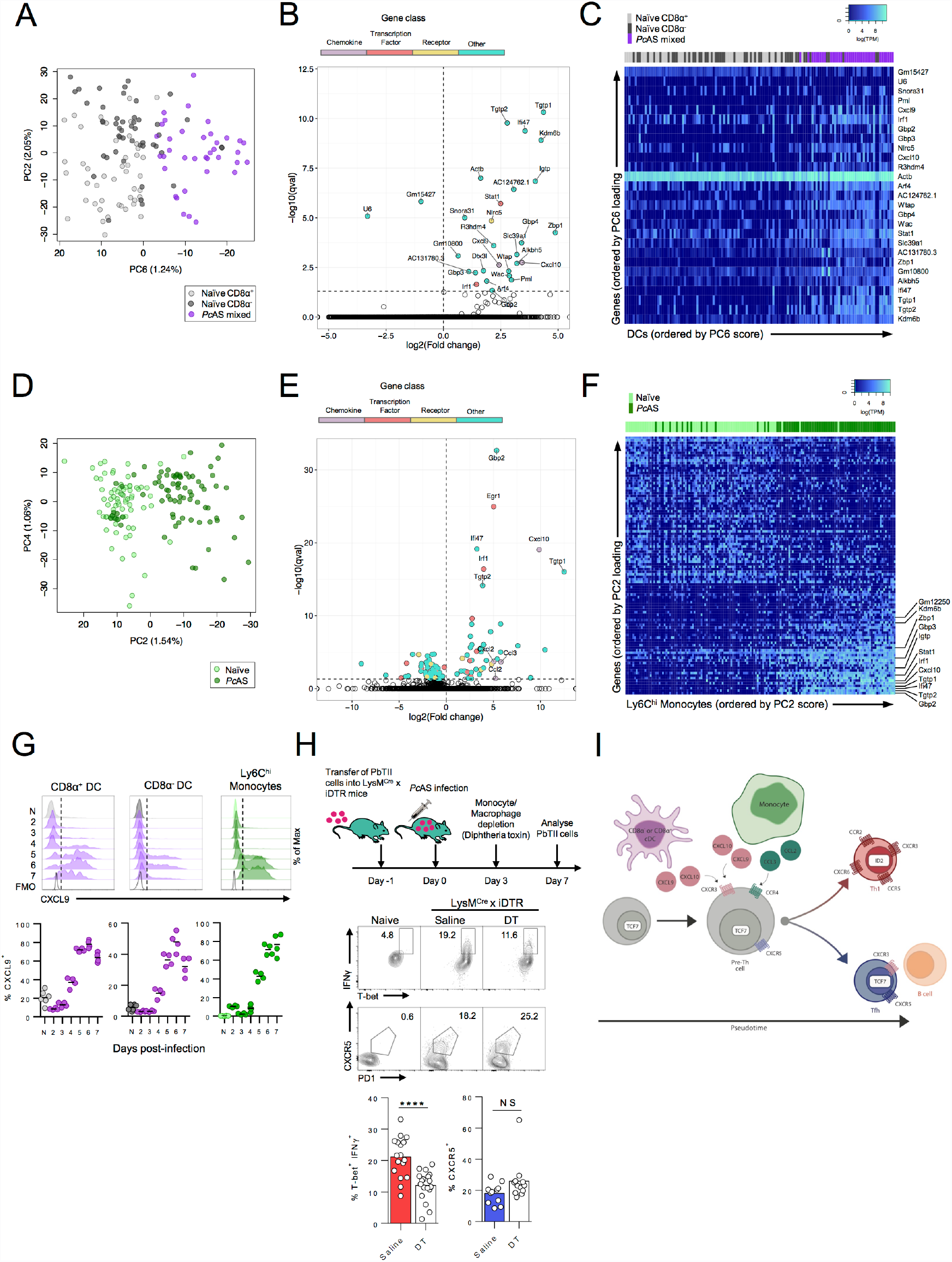
Myeloid cells influence Th bifurcation in uncommitted PbTII cells. (**A-C**) 131 single splenic CD8α^+^ and CD11b^+^ CD8α^−^ cDCs from a naïve mouse, mixed cDCs from a day 3-infected mouse, and **(D-F)** 154 single Ly6C^hi^ monocytes from naïve and infected mice were analysed by scRNAseq, with mRNA reads filtered by minimum expression of 100 TPM in at least 2 cells. (**A** & **D**) Principal Component Analyses, showing PC combinations best separating populations of (A) cDCs, and (D) Ly6C^hi^ monocytes from naïve and infected mice. (**B** & **E**) Volcano plots showing fold-change and confidence for differentially expressed genes (17) between (B) cDCs or (E) monocytes in infected versus naïve mice - genes filtered on expression in >10 cells; genes satisfying qval < 0.05 are represented in colours according to functional classification displayed. Full gene lists are provided in Figure S19 and S21, respectively. (**C** & **F**) Expression heatmaps for significantly (qval<0.05) differentially expressed genes in (C) cDCs and (F) Ly6C^hi^ monocytes, between naive and infected mice: cells and genes are ordered according to PC score and loading respectively, using PC6 for cDCs, and PC2 for Ly6C^hi^ monocytes. The 12 common genes between cDCs and monocyte heatmaps are annotated in (F). (**G**) Representative FACS histograms and proportions of splenic CD8α^+^ cDCs, CD8α^−^ cDCs and Ly6C^hi^ monocytes expressing CXCL9 in naive and infected mice between 2-7 days post-infection - individual mouse data plotted with line at mean; data representative of two independent experiments (n=4 mice/time point/experiment). (**H**) Scheme depicting experimental design: PbTII cells were transferred into *LysM*^*Cre*^ x *iDTR* mice 1 day prior to infection. At 3 days p.i., mice were treated with diphtheria toxin (DT) or control saline, with PbTII Th1/Tfh responses assessed at 7 days p.i.. Representative FACS plots (gated on splenic PbTII cells) showing Th1 proportions (T-bet^hi^ IFNγ^+^) and Tfh proportions (CXCR5^+^) in DT or saline-treated *LysM*^*Cre*^ x *iDTR* mice; data pooled from two independent experiments. Numbers depict proportions within respective gates. Statistics: Mann-Whitney U Test. ****p<0.001; NS, not significant. (**I**) Summary model proposing chemokine interactions between non-bifurcated PbTII cells and myeloid cells support a Th1 fate, while Tfh fates are sustained by B cells.

Given that *Cxcl9-11, Ccl2, Ccl3* and *Ccl5* signal through either *Cxcr3* or *Ccr4*, which were expressed by activated but uncommitted PbTII cells, we next hypothesized that Ly6C^hi^ monocytes, in addition to cDCs, might interact with PbTII cells, thereby influencing Th1/Tfh fate (8). To begin testing this, we first confirmed chemokine expression at protein level by Ly6C^hi^ monocytes, focussing on CXCL9 (Figure 6G). Kinetics of CXCL9 production was similar in cDCs and Ly6C^hi^ monocytes, consistent with a possible role in interacting with CXCR3^+^ PbTII cells. To test whether monocytes could influence Th1/Tfh bifurcation *in vivo*, we employed *LysMCre* x *iDTR* mice, in which Ly6C^hi^ monocytes could be depleted after PbTII cell activation, but before bifurcation (Figure 6H, Figure S22A). We also noted a modest reduction in CD68^+^ macrophages using this approach, with no evidence for depletion of cDCs or marginal zone macrophages (Figure S23). In this transgenic approach, Th1 fates, but not Tfh fates, were supported by monocytes/macrophages (Figure 6H). Together, these data supported a model in which progression of activated, uncommitted PbTII cells towards a Tfh fate was dependent upon B cells (Figure S14), and a Th1 fate was promoted by chemokine-expressing myeloid cells, including Ly6C^hi^ inflammatory monocytes.

## Discussion

By capturing single CD4^+^ T cell transcriptomes over time, and using a novel analysis approach to reconstruct the continuous course of events, we have resolved the bifurcation of naive CD4^+^ T cells into Th1 and Tfh cells at an unprecedented level of molecular detail, and illustrated that external cellular signals influence Th fate around the point of bifurcation. Importantly, the GPfates modelling of scRNA-seq data is not limited to immune cells or single bifurcation events. The mixture of time series model we used can also be combined with existing computational workflows (17, 37) (see section 5.2 of the Computational Supplement). Therefore, it provides the means for high-resolution analysis of differentiation in any cellular system, mainly towards two fates, as shown by our examination of existing embryonic development and lung tissue regeneration data (Comp. Supp. Figure 11), and, in principle, also for differentiation into multiple cell types (Comp. Supp. Figure 12), for example, during haematopoiesis. The filtered expression data and gaussian process models presented in this study can be found on our interactive web application at data.teichlab.org, where users can visualise their own genes of interest.

Our data reveals the developmental relationship between Th1 and Tfh cells on a genomic scale, and shows that the same naïve precursor can give rise to both fates simultaneously. It provides insights for the early stages of differentiation, and describes the order of transcriptional events before and after the bifurcation of Th1 and Tfh fates. To date, this process has remained incompletely characterised. Here, we use pseudo-temporal ordering of cells to reveal the hierarchy of transcriptional regulation of these events at an unprecedented resolution. Our data highlight the importance of stochastic expression of transcription factors as well as chemokine receptors, suggesting a role for noisy gene expression in Th development.

Transcriptomic profiling previously suggested developmental similarities between Tfh and Th1 cells (38), with *in vitro* studies suggesting relatively late bifurcation of Tfh and Th1 cells (39). However, highly immunogenic viral or bacterial infections induced CD4^+^ T cells to segregate into Bcl6^+^ (Tfh) or Blimp-1^+^ (Th1) subpopulations within two days, and by three days, fate-committed Tfh cells had developed (40–42). In our parasitic model, single CD4^+^ T cell transcriptomes remained remarkably similar until four days of infection. Although it is difficult to directly compare viral or bacterial systems with our parasitic model, we speculate that due to infection-related differences in antigen-presenting cell function, antigen load and availability, *Plasmodium* infection in mice does not drive Th bifurcation as early as observed with highly immunogenic viruses or bacteria. Evidence of sub-optimal MHCII antigen-presenting cell function early during *Plasmodium* infection (43, 44) raises the hypothesis that Th bifurcation is sensitive to immune-suppression. Our data indicate that uncommitted, activated CD4^+^ T cells are heterogeneous, but nevertheless closely related at a transcriptional level, suggesting considerable flexibility throughout the proliferative phase of their response. Such plasticity during Th differentiation has been proposed to be beneficial as a means of countering evolution of immune-evasion strategies by pathogens (3).

As CD4^+^ T cells progress from immunological naivety towards a Th fate, they may experience different cellular microenvironments, even within the confines of secondary lymphoid tissue. The observation that bifurcation towards Th1 and Tfh cells was preceded by upregulation of chemokine receptors prompted us to investigate possible interactions with chemokine-expressing myeloid cells. Previous studies have highlighted the potential for cDCs in lymph nodes to produce Th1-associated chemokines (8). Our study, which focused on the spleen, was consistent with this concept, and, furthermore, implicated inflammatory monocytes in Th1 support. However, since our transgenic approach for depleting monocytes also removed a portion of splenic red pulp macrophages, we cannot discount the possibility that red pulp macrophages may partly contribute to a Th1 fate. Nevertheless, our data support a model in which myeloid cells in the spleen influence bifurcation, and support a Th1 fate during *Plasmodium* infection. Moreover, our studies emphasise that although cDCs are the predominant professional antigen-presenting cell for initiating CD4^+^ T cell activation in the spleen, other myeloid cells also exhibit a capacity to influence towards a Th1 fate. In contrast, Th bifurcation towards a Tfh fate was not supported by monocytes/macrophages. Instead, given that CXCR5 was the only chemokine receptor significantly associated with bifurcation towards a Tfh fate, cellular interaction with B cell follicles may be the primary mechanism for supporting activated CD4^+^ T cells towards a Tfh fate. Our model suggests that activated, uncommitted CD4^+^ T cells become receptive to competing chemoattractant signals from multiple cell types in different zones of the spleen. This model focuses on intercellular communication as the main driver of bifurcation. However, upstream of these processes, internal stochasticity in uncommitted CD4^+^ T cells may control the balance of chemokine receptor expression (45), thus mediating differential trafficking and variation in intercellular interactions. Future experiments combining our integrated single-cell genomics and computational approach with *in vivo* positional and trafficking data may reveal molecular relationships between internal stochasticity, migratory behaviour, Th fate and perhaps immunological memory.

## Acknowledgments

We thank the Wellcome Trust Sanger Institute Sequencing Facility for performing Illumina sequencing, Wellcome Trust Sanger Institute Single-cell Genomics Core Facility for single-cell sample processing and the Wellcome Trust Sanger Institute Research Support Facility for care of the mice used in these studies. We thank QIMR Berghofer Flow Cytometry and Animal Facilities for expert advice and care of wild-type and transgenic mice. We wish to acknowledge Stephan Lorenz, Joanna Cartwright and Tom Metcalf for expert technical assistance. We thank Guy Emerton for constructing the database and the interface for accessing the data. We thank Michel Raymond for his work in defining cytokines and cell-surface receptors. Susanna Ng is acknowledged for assistance with graphic design.

This work was supported by European Research Council grant ThSWITCH (number 260507), Australian National Health and Medical Research Council Project grant (number 1028641) and Career Development Fellowship (no. 1028643), University of Queensland, Australian Infectious Disease Research Centre grants and the Lister Institute for Preventative Medicine. KRJ was supported by grants from EMBL Australia and OzEMalaR. FOB was supported by the Lundbeck Foundation. MZ was supported by the Marie Curie ITN grant “Machine Learning for Personalized Medicine” (EU FP7-PEOPLE Project Ref 316861, MLPM2012).

The data presented in this paper is publically available in the ArrayExpress database.

## Supplementary Computational Methods - The GPfates model

### 1 Introduction

GPfates is based on a three-stage approach that first i) infers a low-dimensional representation of single-cell RNA-seq data, then ii) infers pseudotime to iii) model the temporal dynamics of gene expression profiles with a mixture model. These steps build on existing modeling components: The Gaussian Process Latent Variable Model [Lawrence, 2006], and the Overlapping Mixture of Gaussian Processes [Lázaro-Gredilla et al., 2012]. For a graphical illustration of the major steps involved in this analysis, see Figure 2D of the main text (as well as Supp. Comp. Fig 1). In Sections 2 and 3 we describe the statistical models that underlie the components of GPfates. In Section 4 we describe downstream analysis methods for interpreting the fitted model. Finally, in Section 5, we present additional validation experiments using simulations, robustness analyses and by analyzing multiple existing data sets.

**Sup. Comp. Fig. 1:**
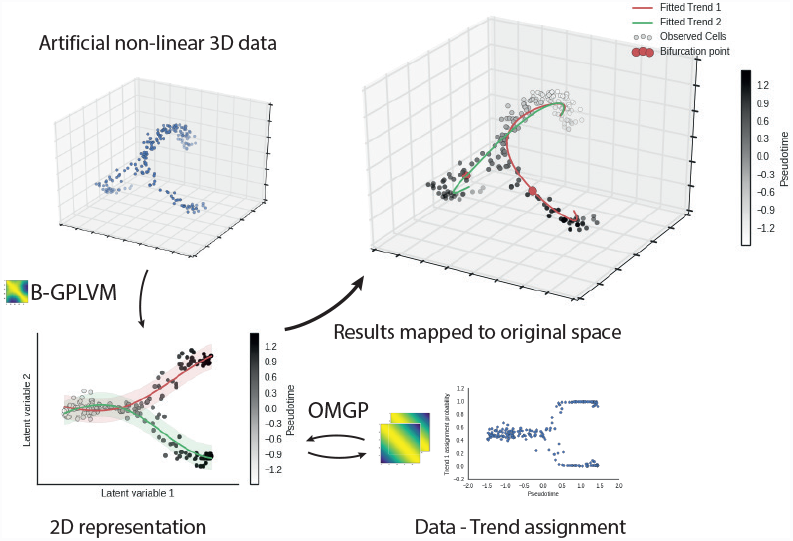
Illustration of the analysis workflow. A low dimensional parametrization of the data is found using Bayesian GPLVM. The low-dimensional representation is viewed as a mixture problem, and solved by an Overlapping Mixture of Gaussian Processes. This allows us to represent our cells as members of different smooth processes. But also interpret in terms of the high-dimensional space parametrized by the GPLVM.

### 2 Pseudotime inference

#### 2.1 Gaussian Process Regression

A main component of GPfates is to model temporal transitions. We use the Gaussian process (GP) framework, thereby casting this problem as non-parametric regression. Let us begin by assuming that the developmental time *t* for each cell we observe is known. Then, the output *y*_*g*_ (i.e. expression of gene *g*) is modelled as a continuous function of the input *t* (i.e. developmental progression)

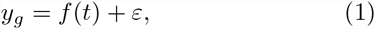

where

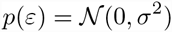

is Gaussian distributed residual noise and *f*(*t*) denotes the unknown regression function. In this work *y*_*g*_ is considered to be an *N*-dimensional vector of *N* cells with observed expression of the gene *g*. We denote the expression of *g* in an individual cell *n* as [*y*_*g*_]_*n*_.

A GP can be interpreted as a function-valued prior on the elements of *f*, which is defined by a covariance function that in turn is parametrized by the input (developmental time) *t*:

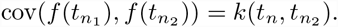

The *covariance function k*(*t*_*n*_1__, *t*_*n*_2__) encodes prior assumptions on the smoothness and lengthscales of the function *f*(*t*). The most widely used covariance function is the Squared Exponential (SE) covariance function,

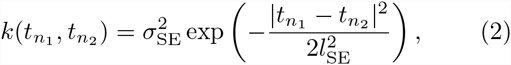

and this is the covariance function we will generally be used in this work. This covariance has the hyperparameters 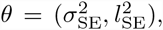, which parametrize the amplitude 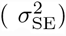 and the lengthscale 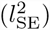 of functions under the prior. Throughout the remainder of the text we will omit the hyperparameters from equations for the sake of brevity. Note that there is a whole compendium of valid covariance functions, which can also be combined using sum or multiplication; see [cite: Rasmussen, GP 2006] for an overview.

We write that a function *f* is *Gaussian Process distributed* by

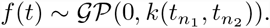

This prior on the function *f* can be linked to the finite observed data using a Gaussian likelihood:

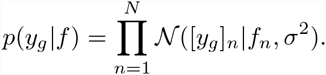

Together with the prior on the corresponding (finite) elements of *f*,

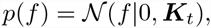

this results in the marginal likelihood

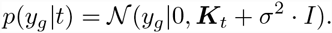

Here *K*_*t*_ is an *N × N* matrix of pairwise evaluations of the covariance functions at the observed times *t*. I.e.

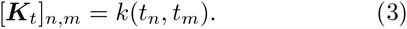

By considering the joint distribution of the observed data *y*_*g*_ and an unseen function value *f*(*t*_⋆_), it is possible to derive the predictive distribution for *f*(*t*_*⋆*_):

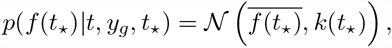

where

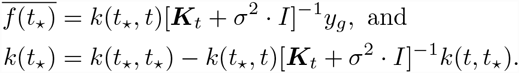

For a full review on Gaussian Processes, see Williams and Rasmussen [2006].

So far, we have only described Gaussian Process Regression for expression *y*_*g*_ of a single gene *g*. If we consider a collection of *G* genes {1,…, *G*}, their expression can be modelled together by

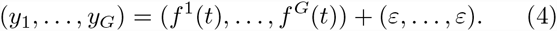

We use *Y* to compactly denote the *N × G* expression matrix of cells *×* genes, where

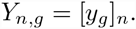

The assumption that all genes are governed by similar functional relationships with *t* means we place the same GP prior (with shared covariance function):

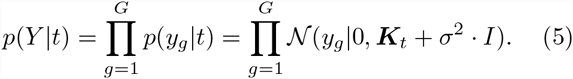

In the next section we will see the usefulness of considering multiple genes at once.

#### 2.2 Pseudotime inference by Bayesian GPLVM with *per*-cell prior

The Gaussian Process regression framework described above assumes we know the time *t* of each cell. While in many single-cell RNA-seq experiments record a collection times over some time-course, these are rather sparse, and it has been pointed out [Trapnell et al., 2014] that cells are sampled from a population where responses are *unsynchronized*. Each cell has reached a certain stage in the differentiation process under investigation, which we do not observe directly. The progress in to this process is referred to as *pseudotime*. We can however infer this from the data. In the Gaussian Process Latent Variable Model (GPLVM) [Lawrence, 2006], we use the multiple output case of Gaussian Process regression (equation 4), but consider the values of *t* to be parameters which we wish to infer.

The joint probability of the GPLVM is

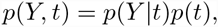

where *p*(*Y*|*t*) is defined in equation 5, and the prior *p*(*t*) is such that for cell *n*,

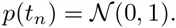

Following Reid and Wernisch [2016], we can also consider the prior *p*(*t*) to be informed about the experimental ordering of collection times of the cells, putting the mean of *t*_*n*_ to correspond to the time point of cell *n*. If we use our Malaria time course as and example, we can put the prior on *t* so that

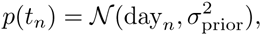

where day_*n*_ ∈ {1, 2, 3, 4, 5} correspond to the collection order of those cells. The parameter 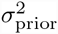 alters the strength of the prior.

The objective of Bayesian GPLVM [Titsias and Lawrence, 2010], is to find the posterior probability distribution 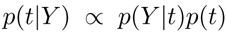. This is intractable though, due to the *t* values appearing non-linearly in the matrix inverse 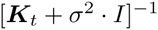.

In Titsias and Lawrence [2010], a lower bound to the marginal likelihood is calculated by estimating the posterior *p*(*t*|*Y*) by a variational distribution *q*(*t*). The distribution

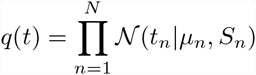

is described in that paper, and Bayesian training of the model to maximize this lower bound. This is the method we use.

Because the scale of *t* is ambiguous, in particular if no priors are specified, we prefer to at times scale the inferred *t* to the range [0, 1] when reporting the pseudotime, to avoid confusion about negative “time”. In these cases we refer to pseudotime as *scaled pseudotime* in the legends.

#### 2.3 Dimensionality reduction

In many cases it is useful to work on a reduced representation of cellular expression profiles. For example, when modelling transcriptomic data, fitting a model to a low-dimensional representation can be preferable to fitting it to expression profiles of thousands of genes. Formally, the objective of *dimensionality reductions* is to find some *M*-dimensional representation of the *G*-dimensional expression measurements, where *M << G*. Typically *M* is 2 or 3, which aides visual interpretation. Analogous to the pseudotime inference, these latent cell states can also be inferred using the GPLVM. Say *X* is an *M × N* matrix so that each cell *n* correspond to an *M*-dimensional vector,

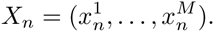

We want to model the expression matrix *Y* so that

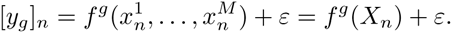

Note that now the covariance function is evaluated as *k*(*X*_*n*_1__, *X*_*n*_2__), where, in the Squared Exponential covariance function in equation 2, the operator |·| is evaluated as the Euclidean norm for vectors, rather than absolute value.

Just as the *t*_*n*_ values are inferred from data above, so can the *X*_*n*_ vectors be inferred from the data.

### 3 Bifurcation inference using overlapping mixtures of Gaussian processes

In a continuous setting, a bifurcating process can be seen as one function, splitting apart into two functions over time. One approach to model this could be to consider two functions throughout time, but before the bifurcation happens, the two functions are identical. With this in mind, we can use a mixture model to tease apart the shared and bifurcated functions.

#### 3.1 Mixture model

Mixture models are hierarchical models where an observation is assumed to be generated from one of *C components*, each of which is described by its own model. The goal of mixture models is to infer which component an observation stems from, and at the same time model that component.

The Overlapping Mixture of Gaussian Processes (OMGP) model [Lázaro-Gredilla et al., 2012] assumes there are *C* different underlying latent functions producing the *N* observed cells. This model was originally developed for the application of missile tracking, and in that setting an observation is e.g. a radar based location at a given real time point. As such, the main focus of the definition of the model is for the case of *C* completely independent components. The approach presented here is based on the realisation that the model would also be able to handle the case of *branching* trajectories. There would simply be a time interval where it does not matter which mixture trajectory data is sampled from. In our setting, an observation is a single cell, and the analog to real time is pseudotime (Supp. Comp. Fig. 2). As an additional extension, we phrase a version of the OMGP model which is non-parametric in the number of trajectories.

**Sup. Comp. Fig. 2:**
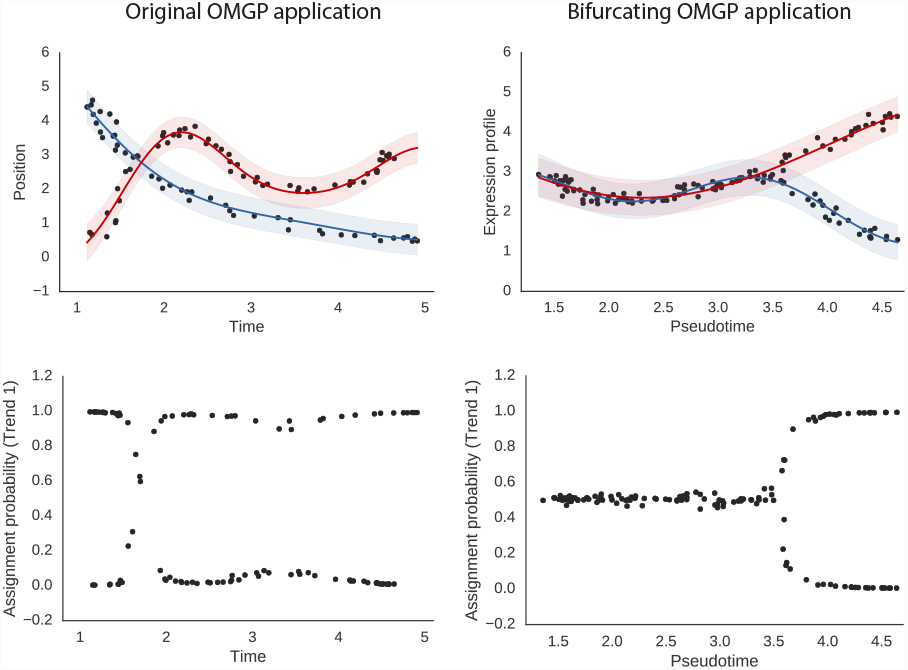
Comparison of the original OMGP use case (left) and our use case (right), in both cases where the number of trends *K* = 2. In the original use case trends are expected to be independent throughout time, albeit with some ambiguity in some locations. In our application, we interpret ambiguous cell assignment to be in a common precursor state.

In the original regression case described in equation 1, data is assumed to be generated by a single smooth unknown function. When modeling our gene expression data with the Overlapping Mixture of Gaussian Processes, data is considered to be generated by

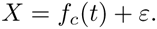

However, we are lacking information about which latent function *f*_*c*_ generated any given observation (*t*_*n*_, *X*_*n*_) of pseudotime and gene expression for the *N* observed cells. Here *X* correspond to some representation of the transcriptional state of the cells. It could be the expression of all genes (*X* = *Y*), a single gene (*X* = *y*_*g*_), or an *M*-dimensional inferred representation as discussed above.

This is viewed as a mixture modelling problem, where each cell has a latent variable *z*_*i*_ specifying to which component *f*_*c*_ the cell should be allocated to. Write *F* for the collection of all latent functions. The covariance functions *k*_*c*_ for each *f*_*c*_ can be different from each other, though for the applications we discuss here, we take them as Squared Exponential covariance functions with different hyperparameter values.

In the OMGP formulation, the likelihood is

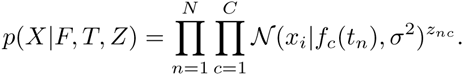

We specify a multinomial prior on the latent variables *Z*, namely

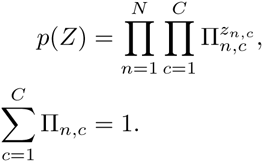

Additionally, each of the latent functions *f*_*c*_ has an independent Gaussian process prior:

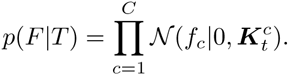

The covariance matrices 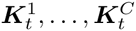 for the latent functions *f*_1_,…, *f*_*C*_ are generated from a covariance functions *k*_1_(*t*_*n*_1__, *t*_*n*_2__),…, *k*_*C*_(*t*_*n*_1__, *t*_*n*_2__) like in equation 3.

Now we rephrase this as a Dirichlet Process Gaussian Process mixture model [Hensman et al., 2015]. Let every latent function *f*_*c*_ have an associated “stick-breaking length” *v*_*c*_, based on the “stick-breaking” formulation of the Dirichlet Process. Here *V* = [*v*_1_, …, *v*_∞_] is the collection of stick-breaking lengths for constructing the Dirichlet process for the assignment. The joint distribution of the OMGP model is

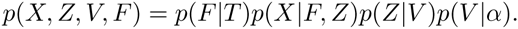

The value *α* is a parameter of the model which controls the expected concentrations of mixtures (which we in practice take as *α* = 1, a common default), and

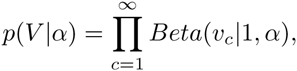

where *Beta*(·, ·) is the beta distribution. The prior distribution over the collection of Gaussian Processes is

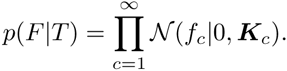

Following the stick-breaking formulation,

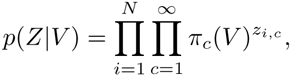

where 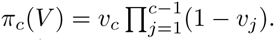

The assignments between observations *X* and the latent functions *F* is given by a binary *N × C* matrix *Z*. The assignments to latent functions are considered as additional variational parameters. Let *ϕ* be an *N × C* matrix where *ϕ*_*nc*_ is the approximate posterior probability of assigning the *n*th observation to the *c*th latent function. The *ϕ* parameters are inferred by collapsed variational inference as described in Hensman et al. [2012]. Overall, the likelihood of the model is

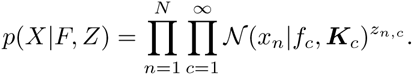

(It should be noted that everything described generalizes to the case where the latent functions *f*_*c*_ are vector valued, as long as all output dimensions of such a function share the same covariance function. In this case, probabilities factorize over output dimensions, but beyond that all calculations are the same.)

#### 3.2 Parameter inference

In Lázaro-Gredilla et al. [2012] the latent variables Z in the parametric version of OMGP were inferred using an expectation-maximization scheme. Here we describe how we perform variational inference for the *ϕ*-parameters in the non-parametric version of the model.

To make the inference problem tractable, the variational distribution *q*(*Z*) is introduced with variational parameters *ϕ*, at a given truncation level *C* such that

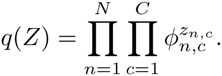

with the objective of approximating *p*(*Z*|*F, X, T*).

The lower bound of the log-likelihood of the OMGP model, which we write as *L*_KL_, when approximating *p*(*Z*) by *q*(*Z*) can be split up in three terms as

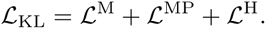

Here 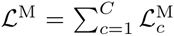 is the log-likelihood of the latent functions as represented by Gaussian processes. For the *c*th latent function, the variational distribution of *f*_*c*_ which maximizes the lower bound was derived in Lázaro-Gredilla et al. [2012] to be

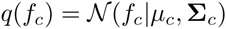

where 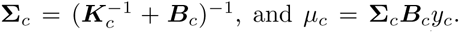 Here ***B***_*c*_ is a diagonal matrix with entries 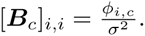. Thus the log-likelihood for a particular latent function *f*_*c*_, assuming we have optimal assignments *ϕ*, is

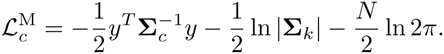

The second and third parts of *L*_*KL*_ were derived in [Hensman2014] as

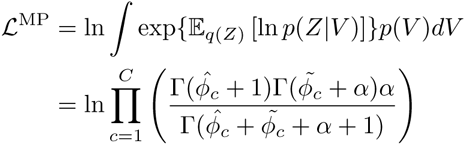

and

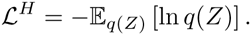

For optimizing variational mixture assignment parameters we follow Hensman et al. [2012], and use *natural gradient descent*. For hyperparameters of the kernels, as well as the variance parameter *σ*^2^ of the model, we perform gradient descent.

If we know 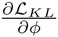 we can calculate the natural gradient by equation (22) in Hensman et al. [2015]. The gradients 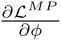 and 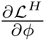 were derived in Hensman et al. [2015], the only unknown part is 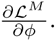

We then use the identity 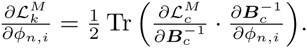 Here 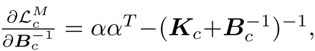 and the matrix 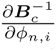 will be zero everywhere, except in the diagonal element (*n*, *n*) where it will be 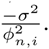

Using the chain rule, we can calculate log-likelihood gradients of the model hyperparameters for any covariance function, since we know 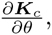 resulting in a very general and modular framework. We only need 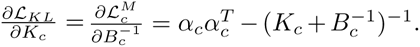 In the case of the model variance *σ*^2^ we have 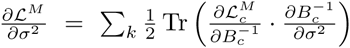 where 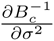 will be a diagonal matrix with 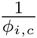 on element (*i*, *i*) for all *i*.

### 4 Downstream analysis

#### 4.1 Ranking genes by bifurcation

Once the OMGP model has been fit, it can be used to investigate individual genes in terms of their bifurcating trajectory.

The log-likelihood of the OMGP model depends on the covariance matrices 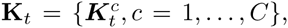 the variational mixture parameter matrix *ϕ*, and the *N* observations (*t*, *X*). Let us assume that we have mixture parameters *ϕ*_*b*_ which have been found to distinguish a bifurcating trend based on some *X* response variables. We can now keep the fitted parameters and evaluate the marginal likelihood of a model where the response variables *X* are replaced by gene expression values *y*_*g*_. We call this new model 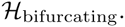 We wish to find genes which fit this bifurcating model better than a model where this is no bifurcation. To this end, we make a third model 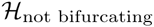 identical to the precious one, except we replace *ϕ*_*b*_ with ambiguous assignments *ϕ*_*a*_. To asses whether a given gene *g* is better described by the *bifurcating* or the *not bifurcating* model, we evaluate the Bayes factor:

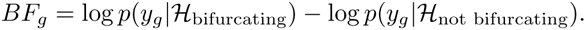

We refer to this ratio as the *bifurcation statistic*.

To estimate p-values, we used a permutation approach where we perform the same analysis for every gene *g*, except with permuted *t* values to estimate a null distribution.

As a proxy for *effect size* of bifurcation, we consider how well the expression values of a gene correlate with the trend assignments to a latent function. Strong positive correlation will mean the gene is particularly upregulated in the cells unambiguously belonging to the trend. Conversely, a strong negative correlation indicates the gene is down-regulated in the strongly assigned cells compared to all cells.

#### 4.2 Inferring the bifurcation time point

It is possible to qualitatively appreciate from the GP assignment probability (*ϕ*_*c*_) for each trajectory (*f*_*c*_(*t*)) of the OMGP model, which cells are *ambiguous* and which cells are *exclusive* to individual GP's. In the case of two trends, ambiguous cells have assignment probability (*ϕ*) close to 0.5. A model where the data can be described by two trends, but not by one, will have a higher likelihood. Similarly, if only a *region* of the *ϕ* parameters over time are replaced by ambiguous cell assignment values, the new model will have a lower likelihood.

For the sake of clarity, we make the assumption that the OMGP will begin as ambiguous, and then become less ambiguous over time, splitting into two trends, in this special case. To investigate these cases, we pick a time-point *t*_*b*_ in an OMGP, then replace all *ϕ* values prior to *t*_*b*_ with 0.5. We define this new *ϕ* as *ϕ>t*_*b*_:

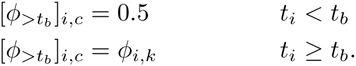

Now we can evaluate the model likelihood for this particular *t*_*b*_ and define

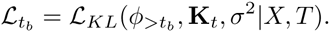

This procedure is repeated for multiple *t*s over the predictor variable of the OMGP model. In our implementation, we consider 30 evenly spaced bins by default, which has given enough resolution for the data investigated (though the number of bins can easily be changed).

The likelihood has to decrease by definition. However, after the bifurcation the decrease is much more pronounced. We use a break-point heuristic to detect this elbow, which is indicative of the bifurcation time.

To identify the region at which the likelihood decreases more rapidly, we fit a piece-wise linear curve to the log-likelihoods, defined by

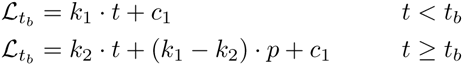

This curve consists of two linear pieces, broken up at the point *p*. When the curve is fitted, we consider the break-point *p* to be the point after which we can be confident that a bifurcation has occurred, see Supp. Comp. Fig 3.

**Sup. Comp. Fig. 3:**
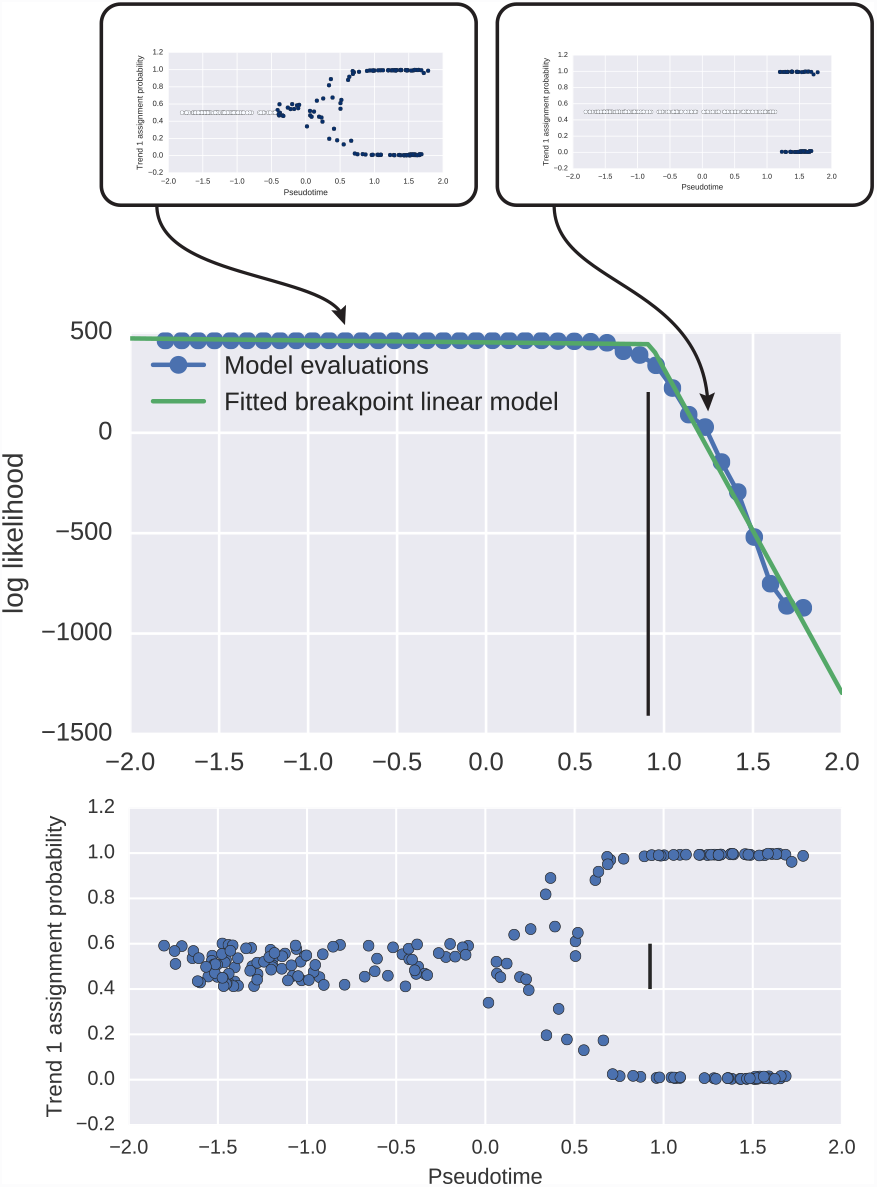
Inferring bifurcation point. The plot illustrates how different points along the pseudo-time are sampled. Ambigous assignment probabilities replaces trained assignment probabilities in the observations earler than the sampled points. The breakpoint model identifies the points where a decrease in likelihood differences becomes more extreme.

### 5 Implementation and combination with existing workflows

#### 5.1 Practical use of GPfates

The basis principle of GPfates is the combination of pseudotime and mixture modelling.

Input to the GPLVM is an expression table consisting of log scaled relative abundance values Transcripts Per Million, TPM, with a value of 1 added to handle cases where expression is 0. As relative abundance follow a log-normal distribution, the Gaussian likelihood used for Gaussian Process regression should be appropriate.

In practice, the pseudotime should represent the biological process of interest. If this process is clear, the expression data should be usable without pre-processing. In single cell time course experiments where the process of interest is less immediate, a strategy highlighted in Trapnell et al. [2014] is to select the gene set used could be to rank the genes by an ANOVA test over the time points, and select a larger number of significant genes.

Similarly, the low-dimensional representation of the transcriptomic cell state should represent the biological response of interest. It can be beneficial to select the parts of the representation which correspond to this. For example, in the analysis of CD4+ T cell time course, we use the second GPLVM latent variable as a representation of T cell response, and model this factor by the OMGP.

While the pseudotime can be inferred directly from the expression matrix *Y*, in many cases it helps interpretation to perform an intermediate step of dimensionality reduction. This process could also be beneficial if the data has a very complex structure.

Another practical consideration we must consider is that single cell expression vales can be quite noisy. This limits the time-scale at which we can expect to measure proper functional differences in expression levels. Due to this, we tend to put lower limits on the lengtscale *l*_SE_ of the squares exponential covariance function.

#### 5.2 Integration of existing methods

We have presented use of the GPfates method when pseudotime or low-dimensional representations have been based on the GPLVM. This is because the OMGP follows from this framework, and the statistical assumptions are consistent between the models.

In practice, other methods for inferring pseudotime or low-dimensional representations could also be considered. Here we briefly outline possible strategies for applications of GPfates downstream of popular single-cell analysis methods.

Recall that to perform the GPfates inference, we need pseudotime *t* and some representation of transcriptomic state *X*. These variables can be set as the output from other methods.

In Monocle [Trapnell et al., 2014], the low-dimensional representation *X* is found by independent component analysis, and the pseudotime *t* for each cell is defined by the path distance to a starting cell through a minimum spanning tree in the coordinates of *X*.

In Wanderlust, a heuristic is used to build a stable k Nearest Neighbor (kNN) graph of the data in the high-dimensional space of protein measurements. The pseudotime *t* for a cell is then defined as the average shortest path from a known starting cell through the kNN graph. Note that for CyTOF data, which Wanderlust is designed for, only up to 40 analytes can be measured at once, so it could be feasible to take *X* to be the original expression matrix (*Y* in our notation).

Another dimensionality reduction technique which have been used for single cell RNA seq data is Diffusion Maps [Haghverdi et al., 2015]. Here *X* is a spectral embedding of the data manifold, based on the Laplace operator. It has been pointed out that these embeddings preserve branching structure in the data. Taking the pseudotime *t* as the Diffusion Pseudotime [Haghverdi et al., 2016], an approximation of geodesic distance over the data manifold (from a known starting cell), based on the diffusion map, GPfates modelling could be used downstream to quantify the branching structure of the data.

We list alternative compatible pseudotime methods in table 1.

**Table 1:** Examples of other pseudotime methods and bifurcation methods.

| Pseudotime Method | Strategy | OMGP Compatibility |
| --- | --- | --- |
| Monocle Pseudotime<br>[Trapnell et al., 2014] | Minimum Spanning Tree path length in 2D ICA space | Yes |
| Diffusion maps<br>[Haghverdi et al., 2015] | Spectral embedding of data manifold | With postprocessing, e.g. DPT<br>[Haghverdi et al., 2016] |
| Wanderlust<br>[Bendall et al., 2014] | Heuristic k-NN graph geodesic distance | Yes |
| GPLVM | Latent data parametrization | Yes |
| SCUBA Pseudotime<br>[Marco et al., 2014] | Principal curve in 3D t-SNE embedding | Yes |
| Bifurcation Method | Strategy |  |
| SCUBA<br>[Marco et al., 2014] | Transitions between clusters in pseudotime bins |  |
| Monocle states<br>[Trapnell et al., 2014] | Create k PQ trees from a Minimum Spanning Tree |  |
| OMGP | Model data as mixture of continuous processes |  |

As a demonstration, we generated a toy data set with three branches, and extracted the pseudotime using both the Monocle method and the Wanderlust method. Then fitted and OMGP with *C* = 3 on the output. The results can be seen in Comp. Supp. Fig 4, which illustrates the correct identification of the branching processes for either input.

**Sup. Comp. Fig. 4:**
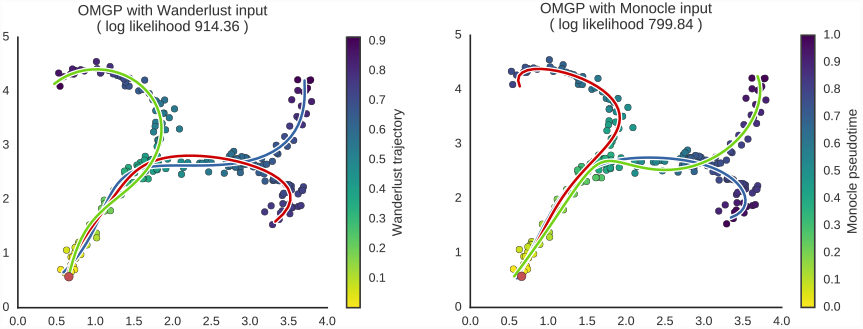
OMGP is compatible with e.g. Wanderlust and Monocle, as demonstrated with a toy data set

#### 5.3 Software availability

We have made a software package for using the GPfates method, which is available at https://github.com/Teichlab/GPfates. It provides guidance and sensible defaults for the kind of analysis we have described here. It makes extensive use of the `GPy`^1^ package, and the `GPclust`^2^ package, where we implemented the nonparametric OMGP model.

### 6 Assessment of GPfates on simulated and real data

#### 6.1 Sample-size robustness analysis

Our full analysis consists of several independent consecutive steps: first the GPfates method where we are i) finding a low-dimensional representation, ii) smoothing the data over a pseudotime, and iii) finding a trend mixture model. After this we perform downstream analysis where we are iv) dentifying the end states and bifurcation.

How much data do we need to accurately reconstruct trends from all four of the above steps, and how much data is needed for individual steps? We investigated both how stable the full procedure is, as well as the individual steps, by re-running it on sub-sampled datasets with fewer cells than the entire dataset.

To measure the stability of the methods, we consider absolute Pearson correlation of the parameters inferred for sub-sampled data relative to the results obtained from performing the analysis on the full data set.

We found that recovering a low-dimensional representation is extremely stable with respect to the number of cells (Supp. Comp. Fig. 5), with almost perfect correlations between analysis of the sub-sampled data and the original GPLVM values. (For example, the lowest absolute Pearson correlation for a run with 50 cells was 0.96). Similarly, the pseudotime inference is also very stable to sub-sampling.

**Sup. Comp. Fig. 5:**
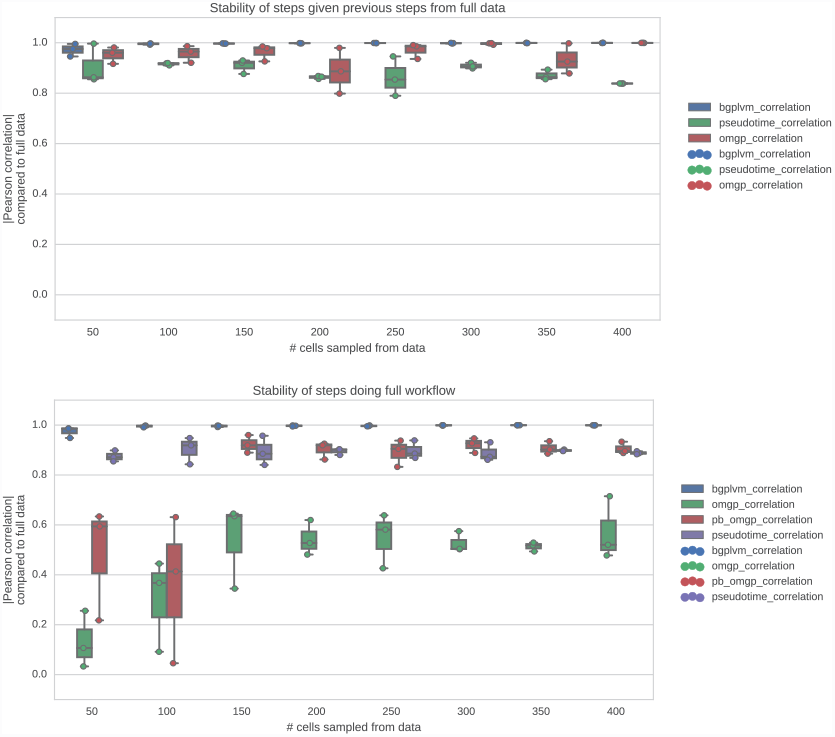
Robustness of analysis steps by subsampling. Parameters inferred from a subsample of the data are compared to parameters inferred using the full data. The top panel indicates this analysis for independent steps assuming the previous step is known. The lower panel shows the result when running the workflow from start to end.

Finding the entire OMGP mixture model over pseudotime requires a larger number of cells. We don't see any higher degrees of consistency until we reach 150 sub-sampled cells, with correlations around 0.5. It is rare to see single cell studies with so few cells, and in the study accompanying this text we had many more cells (408). Identifying only the end states is rather robust (but in many cases might be best analyzed as a cluster problem rather than a continuous value problem), where we start seeing a correlation of 0.9 at 150 cells.

The individual steps were in general very stable to sub-sampling, relative to the “gold standard” of using the full data set. When running the entire procedure, we see that smaller errors early on in the analysis will propagate and affect later steps.

#### 6.2 Predicted bifurcation time is not biased by collection times

We consider the risk that the identified bifurcation point in the CD4+ T cell data potentially just reflects the time points at which we have collected data. We test the robustness of the prediction of the bifurcation as having happened at Day 4 by re-running the analysis after removing cells collected at Day 4. In this analysis we find that the bifurcation happens at some point between Day 3 and Day 7 where we don't have any observed cells. The alternate hypothesis would have been that the bifurcation would be found in either Day 3 or Day 7. This provides confidence both in the bifurcation point identification, and more generally in the meaningfulness of the low-dimensional GPLVM representation of the data (Supp. Comp. Fig. 6).

**Sup. Comp. Fig. 6:**
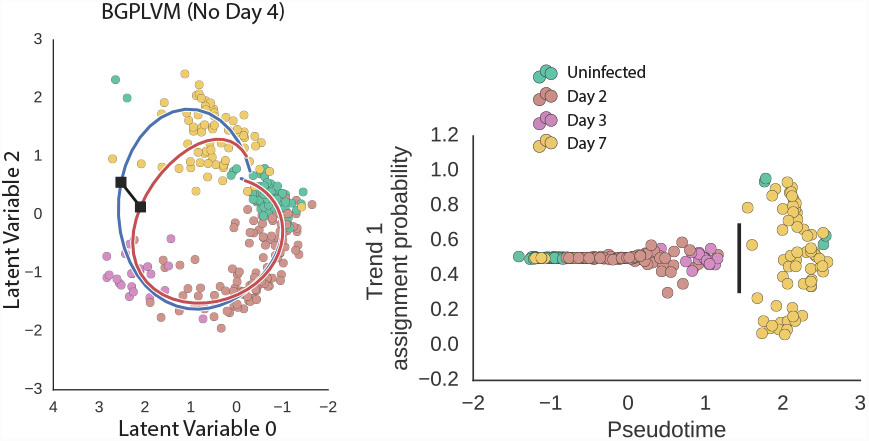
Complete reanalysis of our T-cell data excluding cells cellected at day 4. The bifurcation point is identified as being between Day 3 and Day 7, and is not forced in to either of the days.

#### 6.3 Assessment of the ability to select the number of trends in OMGP

In principle, the marginal log likelihood of the OMGP model should let us select the *C* number of trends which optimally explain the data. We investigated this by generating four synthetic data sets with between 1 and 4 underlying trends. For each of the data sets, we optimized OMGP models with the number of trends *C* varying from 1 to 9 (three times per *C* value). We found that the marginal likelihood of the models corresponded to the correct number of trends in the cases of 3 and 4 ground truth trends, but not for the 1-trend and 2-trend synthetic data. For 1 trend, the likelihood was lowest for a larger number of trends, and for 2 trends, the likelihood was very similar for 2 and 3 trends. This suggests that the OMGP may have a tendency to overestimate the number of trends if there is a single progression. Supp. Comp. Fig. 7.

**Sup. Comp. Fig. 7:**
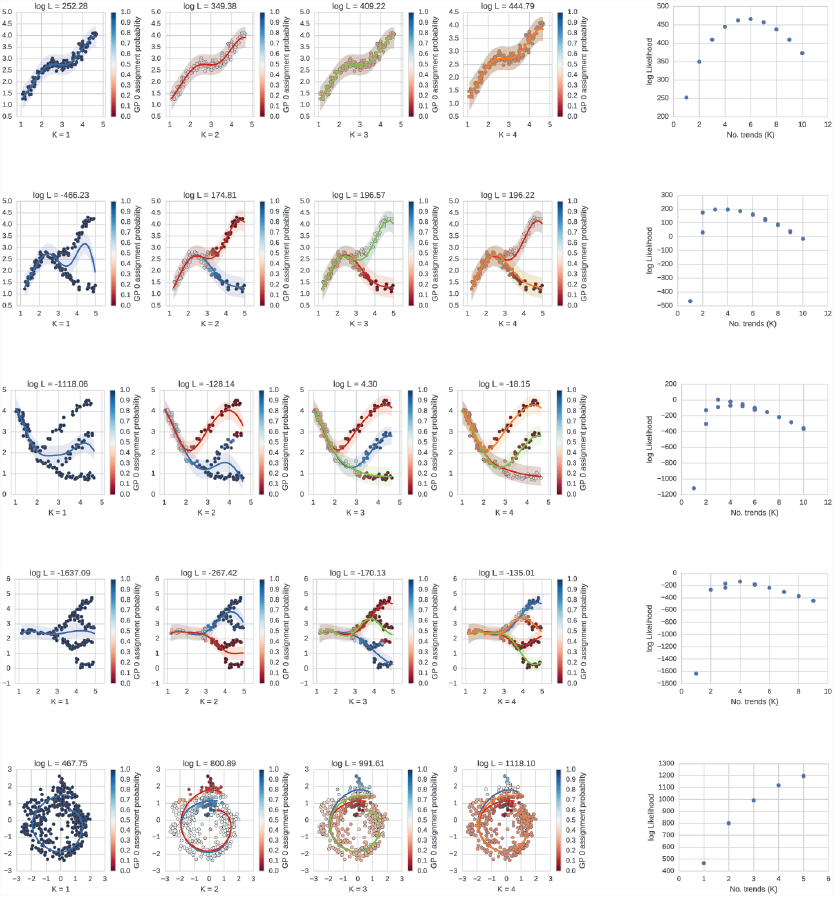
Attempts detecting number of trends with OMGP. Simulated data with expected numbers of trends where fitted with OMGP, where the *C* cutoff was set to a range of values.

For our CD4+ T cell data, we found that the marginal likelihood continuously increased with the number *C*. We elected to keep the model simple and made the assumption that we could sufficiently explain the data with *C* = 2.

It is possible that the optimal likelihood for K is not well defined when we have trends branching off from a common trend. In the original application of the OMGP model, the assumption is that the trends will be completely independent of each other. As we are already to some extent failing to fit two models in the ambiguous case, this might cause the likelihood to reflect a poor fit. For quantitatively determining the number of trends in the data, further work is needed, probably with a model which explicitly considers branching from a common original trend. The marginal likelihood of the model is an indication, but the choice of *C* should also reflect the biological system under consideration.

#### 6.4 Comparison of pseudotime inference with and without priors

For the 1-dimensional Bayesian GPLVM which we use to find the pseudotime of the data, we put priors on the cells based on their known time points. This is not strictly necessary, but helps to enhance interpretability as there is intrinsic invariance to the inferred values. If we do not use priors, qualitatively, the same trajectory is identified. Additionally, comparing the two versions of pseudotime against each other, we see that they correspond to a circular shift relative to each other. The covariance matrices inferred using either strategy have a very similar block structure (Frobenius norm … of difference) indicating that neighbor relations are consistent. Supp. Comp. Fig. 8.

**Sup. Comp. Fig. 8:**
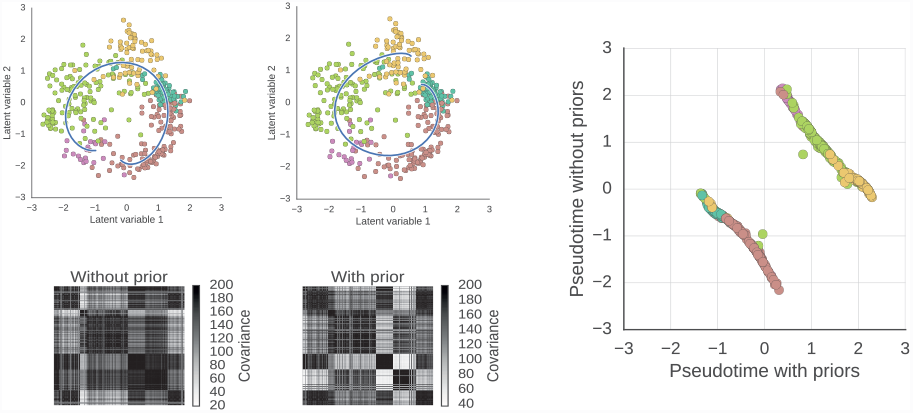
Comparison of pseudotime with and without per-cell priors. The upper left shows the fit of the pseudotime predicted in to the 2D GPLVM with and without priors. Below are the corresponding inferred covariance matrices. The right plot shows the relations between the two versions of pseudotime, clearly indicating that they have an approximate one-to-one mapping.

#### 6.5 Pseudotime uncertainty

As pointed out in Campbell and Yau [2015], we can use the posterior distribution of pseudotime from the Bayesian GPLVM to assess how meaningful the ordering is. By investigating the confidence intervals of the pseudotime for each cell compared to neighboring cells, we see that the ordering is quite meaningful (few cells overlap in confidence interval). (Supp. Comp. Fig. 9)

**Sup. Comp. Fig. 9:**
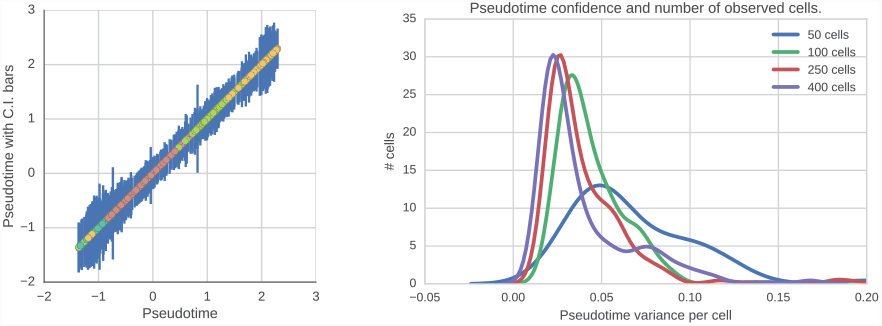
Investigation of uncertainty of inferred pseudotimes. Left panel, since the Bayesian GPLVM fits the variance of the pseudotime for each cell, we can compare the assignments with each other. The bars correspond to 95% confidence intervals. On the right panel we see how the lengths of the confidence intervals globally decrease as the number of cells used increases.

We also investigated how the confidence of the pseudotime depends on the number of cells observed. As the number of observed cells increases, the distribution of variance per cell decreases towards zero. (Supp. Comp. Fig. 9)

#### 6.6 Stability of the circular shape of the GPLVM representation

We wanted to rule out the possibility that the latent variable representations of data which appear circular might be artifacts due to random noise, as suggested by Diaconis et al. [2008]. To make sure this was not the case for our CD4+ T cell data, we removed two ‘slices’ of cells from the circular 2D GPLVM pattern. Following this, we fitted a new GPLVM with this reduced data set. After optimizing the GPLVM, a representation was found which was again missing the same slices, Supp. Comp. Fig. 10A. (The correlation between the two representation for the common cells is very high as well, XX). This control experiment strongly suggests that the GPLVM learns the actual topology of the data.

**Sup. Comp. Fig. 10:**
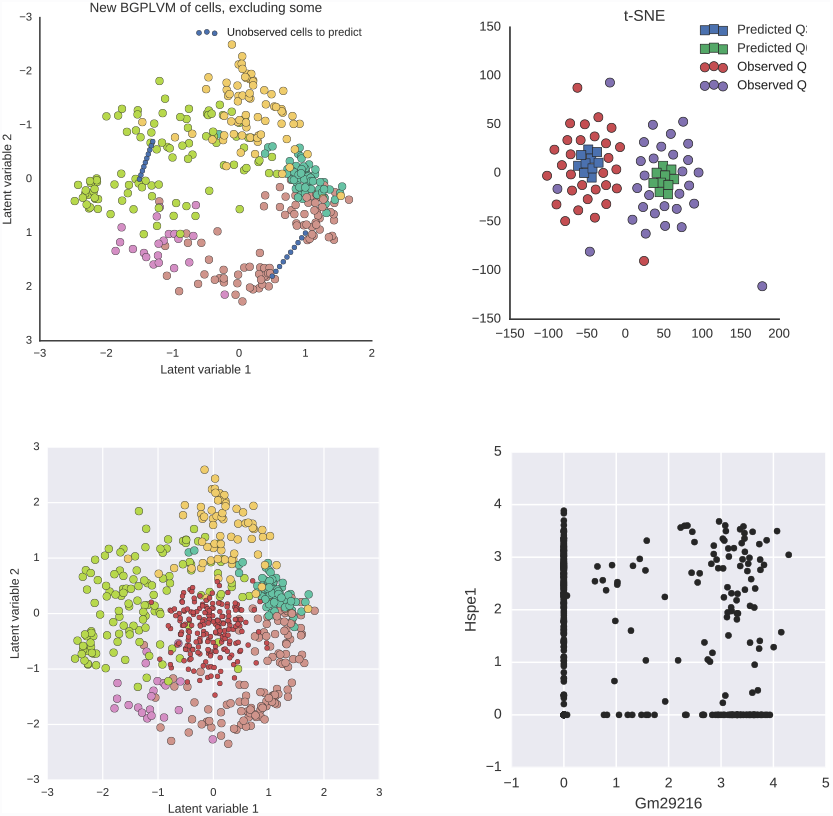
Stability of GPLVM representation, and prediction through GPLVM. Top row: Predicting cells from regions of show higher similarity with left out real cells from corresponding regions than non-corresponding regions. Bottom row: Predicting cells from unobserved regions potentially identifies antagonizing gene combinations.

#### 6.7 Assessing the accuracy of imputing virtual cells

Unlike many other dimensionality reduction techniques, the GPLVM creates a model which maps into the high dimensional observed space. It is, however, not clear how meaningful this representation is. We assessed this by taking the “slice-less” model described above, and in the empty areas corresponding to the removed cells, predicting “virtual cells” (Supp. Comp. Fig 10A). Using an independent clustering technique, t-SNE [Van der Maaten and Hinton, 2008], on both the left out slices of cells and the predicted virtual cells, we find that single cell transcriptomes predicted from a given slice coincide with the real cells from the corresponding slice (Supp. Comp. Fig. 10). This indicates that GPLVM prediction in to high-dimensional spaces is not simply producing overfitted results.

Following from this, we investigated the “hole” in our CD4+ T cell data. We create number of virtual cells from the hole region and compare which genes would be expressed in these cells compared to genes expressed in all cells (Supp. Comp. Fig. 10C). The underlying reasons for data being non-linear is that particular combinations of gene expression patterns do not occur together. If we find genes which are high in the virtual cells but are not observed at the same time in actual cells, this might indicate that they are incompatible with each other. This might be a good complementary tool for generating hypotheses about regulation. For instance, we identified the genes Hspe1 and Gm29216 which would be co-expressed in the hole, but are generally not co-expressed in observed cells (Supp. Comp. Fig. 10D).

### 7 Validating the BGPLVM and OMGP approach by application to other data sets

In order to further corroborate our analysis approach, we considered two recently published single cell data sets produced to investigate progression of single cells in two developmental contexts: mouse fetal lung and human fetal primordial germ cells.

#### 7.1 Analysis of lung development data

We downloaded the data from Treutlein et al. [2014] and quantified the expression using Salmon. To smooth the data over pseudotime, we found genes that vary over the *a priori* known time points by a likelihood ratio test of an ANOVA model of the time points. The expression values for the top varying genes were run on a GPLVM. One of the factors of the optimized GPLVM was used as pseudotime, and the top two factors of the GPLVM were used to represent the entire data set. An OMGP was then optimized on this low-dimensional representation to identify the two trends corresponding to the AT1 and AT2 lung cell lineages without prior annotation. The bifurcation statistic for all expressed genes in these cells reconstituted many of the genes identified in a largely manual manner by Treutlein et al. [2014].

#### 7.2 Analysis of human primordial germ cell data

The data from Guo et al. [2015] was downloaded and quantified with Salmon as with the other data, but with an index based on the human transcriptome: Ensembl 78 annotationa of GRCh38, together with ERCC sequences and human specific repeats from RepBase. To smooth the time course data, we used a likelihood ratio test to find the top genes which were described linearly along the time points in the data. The expression of these genes were then used to fit a GPLVM. This low-dimensional representation of the data was then used to fit an OMGP, taking one of the latent factors as pseudotime.

In this data set, the ground truth about the sex of the cells is known, and thus we could have use a supervised approach such as `GPTwoSample` [Stegle et al., 2010] or `DETime` [Yang et al., 2016]. Interestingly, the OMGP model identifies the split between male and female cells in an unsupervised fashion.

We applied the bifurcation statistic test to identify genes which follow the male and female development differently.

Unlike in the case of the lung development data, the majority of the genes we identify are not discussed in the original study. In the original study, the authors focused on genes specific to given conditions (e.g. Male PGC's from week 11 compared to all other cells). In our analysis, we consider the dynamics of gene expression over development. We find that in the male branch, the GAGE family is highly upregulated over development. Additionally we find the Y-linked gene ENSG00000279950. Also among the top male hits is RHOXF2, a gene linked to male reproduction [Niu et al., 2011]. Further down the list we also interestingly find PIWIL4, a gene with function in development and maintenance of germline stem cells [Sasaki et al., 2003]. On the female side, the top hit is MDK, a gene involved with fetal adrenal gland development (by similariry: UniProtKB P21741). Other top hits include MEIOB, a meiosis related gene, and the satellite repeat GSATII. Surprisingly, we also see upregulation of SPATA22, a gene associated with spermatogenesis.

### 8 Discussion

We have demonstrated our GPfates method, where we use latent variable modelling to infer temporal expression dynamics, and Gaussian process mixture modelling to identify diverging global trends. The method has been investigated in terms of robustness, and applied on several simulated and real data sets showing good results.

Of course there is no silver bullet for these sorts of problems, and it would not be surprising if other methods than the ones we have used work better for some biological systems. We have illustrated that the main component, the Gaussian process mixture modelling, is compatible with other methods in these cases.

A benefit from the methods we use is that diagnostics such as marginal likelihood can be used to aide the user with regards to the models to use. Still, the user will need to keep the biological system in mind, and be critical of results.

**Sup. Comp. Fig. 11:**
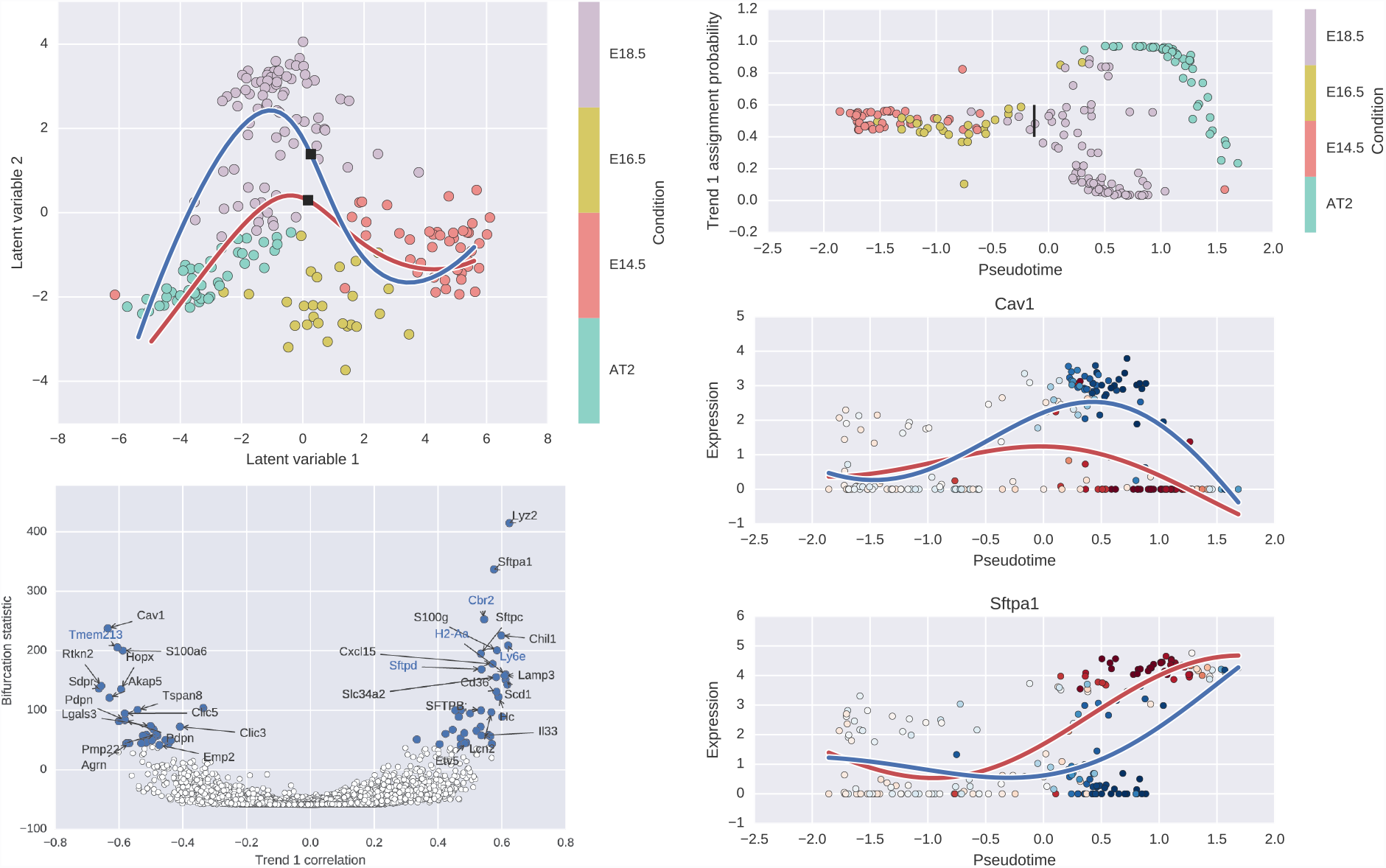
Summary of GPfates result of Treutlein et al developing lung data.

**Sup. Comp. Fig. 12:**
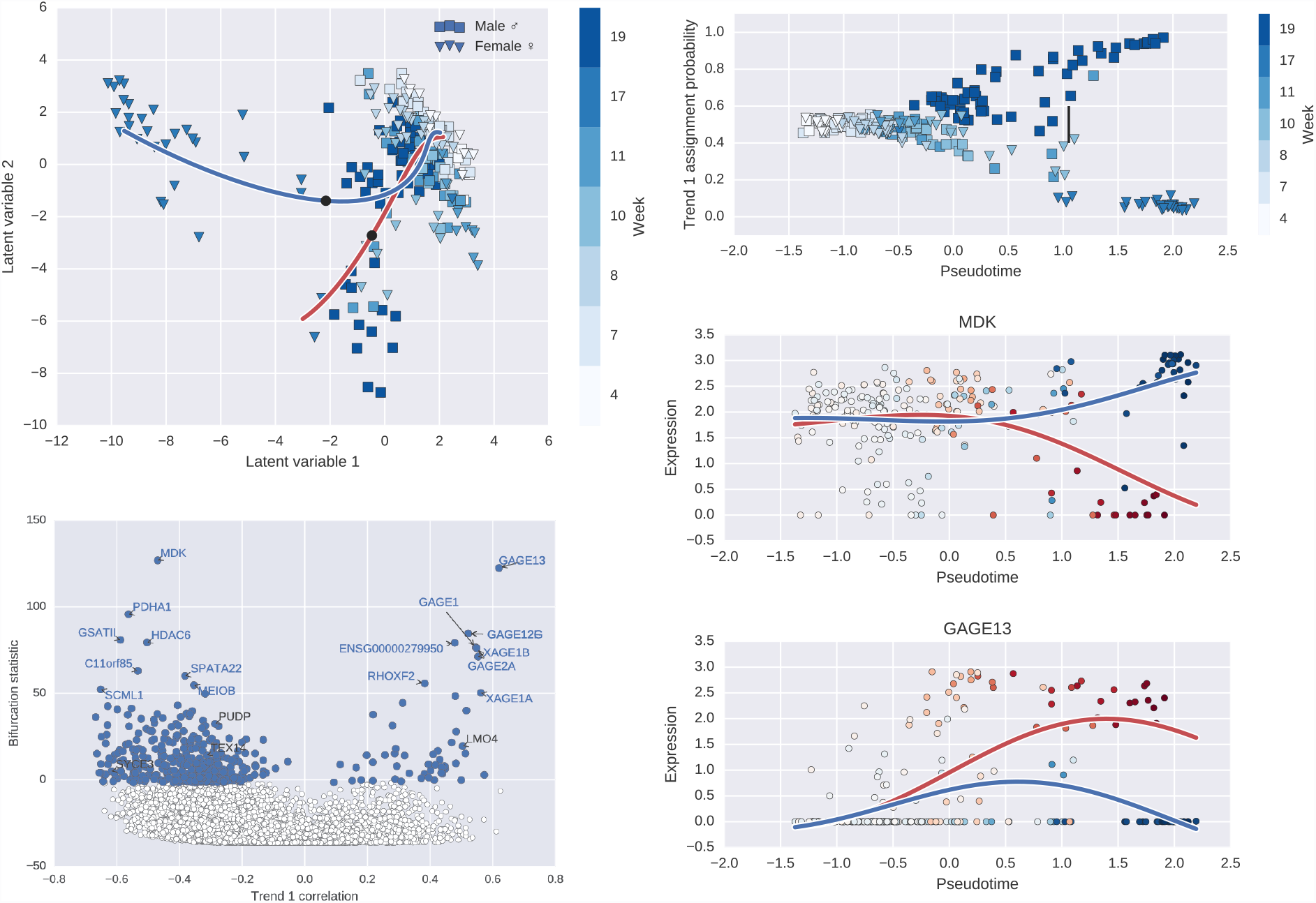
Summary of GPfates result of Guo et al developing primordial germ cell data.

## Materials and Methods

### Ethics and approval

All animal procedures were in accordance with the Animals (Scientific Procedures) Act 1986 and approved by the Animal Welfare and Ethical Review Body of the Wellcome Trust Genome Campus, or in accordance with Australian National Health and Medical Research Council guidelines and approved by the QIMR Berghofer Medical Research Institute Animal Ethics Committee (approval no. A02-633M).

### Mice

C57BL/6 mice were purchased from Australian Resource Center (Canning Vale) or bred inhouse. C57BL/6, PbTIIxCD45.1 and *LysMCre* x *iDTR* mice were maintained under specific pathogen-free conditions within animal facilities at the Wellcome Trust Genome Campus Research Support Facility (Cambridge, UK), registered with the UK Home Office, or at QIMR Berghofer Medical Research Institute (Brisbane, Australia). All mice were female and used at 8-12 weeks of age.

### Adoptive transfer

Spleens from PbTIIxCD45.1 mice were aseptically removed and homogenised through a 100µm strainer before lysis of erythrocytes with RBC lysis buffer (eBioscience). CD4^+^ T cells were enriched (purity >80%) using CD4 microbeads according to the manufacturer's instructions (Miltenyi Biotech) and stained with CellTrace™ Violet (Invitrogen) at 1µM in PBS for 15 minutes at 37ᵒC in the dark. Violet CellTrace-labelled cells were resuspended in PBS and injected (10^6^/200µl RPMI) via a lateral tail vein.

### Infections

*Plasmodium chabaudi chabaudi* AS parasites were used after one *in vivo* passage in WT C57BL/6 mice. Mice were infected with 10^5^ pRBCs i.v. and blood parasitemia was monitored by Giemsa-stained thin blood smears obtained from tail bleeds.

### Flow cytometry and cell isolation

Single-cell suspensions were prepared by homogenising spleens through 100 µm strainers and lysing erythrocytes using RBC lysis buffer (eBioscience). Fc receptors were blocked using anti-CD16/32 antibody (BD Pharmigen or in-house). T cells were stained with the following antibodies (Biolegend): CD4-APC (GK1.5), TCRβ-APC-Cy7 (H57-597), CD45.1-FITC (A20), Vα2-FITC (B20.1), Vβ12-eFluor710 (MR11-1) (eBioscience), CD69-PE (H1.2F3), PD-1-APCCy7 (29F.1A12), CXCR5-Biotinylated (2G8) (BD Pharmigen), Streptavidin-PeCy7, CD183 (CXCR3)-PE (CXCR3-173), IFNγ-BV421 (XMG1.2), Bcl6-PercpCy5.5 (K112-91) (BD Pharmigen), Ki-67-PE (16A8), T-bet-eFluor660 (4-B10) (eBioscience) and TCF-1-PE (CD63D9) (Cell Signaling Technology). Dendritic cells and monocytes were stained with the following antibodies (Biolegend): CD11c-Percp-Cy5.5 (N418), MHCII (1-A/1/E)-APC (M5/114-15.3), B220-AlexaFluor700 (RA3-6B2), TCRβ-APC-Cy7 (H57-597), Ly6C-FITC (HKJ.4), CD11b-BV421 (M1/70), Ly6G-PE (IA8), CD8-PE-Cy7 (53-6.7) and CXCL9-PE (MIG-2F5.5). Intracellular staining for IFNγ, T-bet, Bcl6, Ki-67, TCF-1 or CXCL9 was performed with the eBioscience FoxP3 intracellular kit. Intracellular staining for p-S6 was performing using a monoclonal antibody (D57.2.2E), or respective isotype control (Cell Signaling Technology) with Cell Signaling Buffer Set A (Miltenyi Biotech) according to manufacturer's protocol.

For DNA/RNA staining, Hoechst33342 (10µg/ml; Sigma) was added at 1/500 v/v to cell preparation 15 minutes prior to acquisition using a BD LSRFortessa IV (BD Bioscience).

Cells were sorted using a MoFlo XDP (Beckman Coulter), a FACSAria II (Becton Dickinson) or an Influx (Becton Dickinson) instrument. Activated PbTII T cells were sorted as CD4^+^TCRβ^+^ and CD69^+^ and/or divided at least once as measured using the CellTrace Violet proliferation dye. Dendritic cells were sorted as CD11c^hi^MHCII^hi^TCRβ^−^B220^−^. Naive dendritic cells were further sorted as CD8α^+^CD11b^−^ or CD8α^−^CD11b^+^. Inflammatory monocytes were identified as CD11b^hi^Ly6C^hi^Ly6G^lo^TCRβ^−^B220^−^.

### Single-cell mRNA sequencing

Single cell capture and processing with the Fluidigm C1 system was performed as described in (52). The cell suspension obtained from sorting was loaded onto the Fluidigm C1 platform using small–sized capture chips (5-10µm cells). 1 µl of a 1:4000 dilution of External RNA Control Consortium (ERCC) spike-ins (Ambion, Life Technologies) was included in the lysis buffer. Reverse transcription and pre-amplification of cDNA were performed using the SMARTer Ultra Low RNA kit (Clontech).

For processing with the Smart-seq2 protocol (29), the cells were sorted into 96-well plates containing lysis buffer using either a MoFlo XDP (Beckman Coulter) or an Influx (Becton Dickinson) instrument. The Smart-seq2 amplification was performed as described in (29), with the lysis buffer containing Triton-X, RNase inhibitor, dNTPs, dT30 primer and ERCC spike-ins (Ambion, Life Technologies, final dilution 1:100 million). The cDNA amplification step was performed with 24 cycles.

The sequencing libraries were prepared using Nextera XT DNA Sample Preparation Kit (Illumina), according to the protocol supplied by Fluidigm (PN 100-5950 B1). Libraries from up to 96 single cells were pooled and purified using AMPure XP beads (Beckman Coulter). Pooled samples were sequenced on an Illumina HiSeq 2500 instrument, using paired-end 100 or 125-base pair reads.

### Processing and QC of scRNA-Seq data

Gene expressions were quantified from the paired end reads of the samples using Salmon (41), version 0.4.0. An example command for a one sample would be “salmon quant -i mouse_cdna_38.p3.78_repbase_ercc_index -l IU -p 4 -1 1771-026-195-H4_1.fastq -21771-026-195-H4_2.fastq -o1771-026-195-H4_salmon_out -g mouse_cdna38.78_repbase_ercc_index_gene_map.txt”. The parameter libType=IU, and a transcriptome index built on Ensembl version 78 mouse cDNA sequences. We also had sequences from the ERCC RNA spike-ins in the index, as well as 313 mouse specific repeat sequences from RepBase to potentially capture transcribed repeats.

For quality control of the single-cell data we assessed the number of input read pairs, and the amount of mitochondrial gene content. For all cells, we considered samples with less than 100,000 reads or more than 35% mitochondrial gene content as failed. For T cells, we additionally considered cells where number of genes was less than 100 + 0.008 * (mitochondrial gene content) as failed. For the data generated using a 96-well plate-based Smart-seq2 protocol, which does not permit visual inspection of the captured cells, we additionally excluded low-quality cells from which fewer than 2000 genes were detected, motivated by negative control wells. To verify that that the cells sorted in the wells were PbTII cells, we only selected cells from which both the transgenic TCR alpha and beta chains were detected (Supplementary Tables 2 and 3). Excluded cells were removed from all further analyses, and the remainder of the samples were taken as individual single cells.

For expression values, the Transcripts Per Millions (TPM's) estimated by Salmon included ERCC spike-ins. Thus, for analysis of the cells, we removed ERCC's from the expression table and scaled the TPM's so they again summed to a million. This way we get *endogenous* TPM values, representing the relative abundance of a given gene *within a cell*. We also globally removed genes from analysis where less than three cells expressed the gene at minimum 1 TPM, unless stated otherwise.

### Latent Variable Modelling of data

We modelled the data using an unsupervised Bayesian Gaussian Process Latent Variable model (BGPLVM) (14) on log10 transformed TPM values (with a scaling factor 1 added). The BGPLVM was run with 5 latent variables. As we used an ARD (Automatic Relevance Determination) squared exponential covariance function, we could infer that two latent factors explained the data. All other parameters to the BayesianGPLVM model in GPy (version 1.0.9) were left as default. Upon inspection we noticed a circular pattern. This corresponds to a 1-dimensional topology, which requires two dimensions for a faithful representation. Thus we inferred a new latent variable by 1-dimensional BGPLVM, with priors on the latent variable based on the cell collection times (see the Computational Supplement), where we used the 2D latent variables as input. This way we inferred smoothed “pseudotime” values for the every data point representing the progressive response to the malaria infection.

In the 2-dimensional model of the data, we searched for genes that highly correlated with either of the two explanatory latent factors. Performing functional enrichment analysis using gProfiler (42) on the top genes revealed that factor 1 (which explained most of the variation) was related to cell proliferation. The second factor was largely explained by genes involved with immune response. Upon inspection, it seemed as the second latent factor terminated in two groups of cells. We investigated this in terms of a bifurcating time series.

### Bifurcating time series analysis

To study the cells in terms of a bifurcating time series, we implemented an Overlapping Mixture of Gaussian Processes (OMGP) model (16), see the Computational Supplement for details. The model uses an optimization procedure to associate observed data with a given number of individual independent trends over a time variable. The model was run with pseudotime as input, and the immune response related latent variable as output. For the mixture model, we assumed two trends. The two trends were given squared exponential covariance functions, where we fixed the length scale to 1 based on our prior assumptions on smoothness over pseudotime. We also constrained the model variance to 0.05, which allows trends to share observed data points. Remaining hyperparameters were optimized by gradient descent. (See the Computational Supplement for details)

### Testing genes for bifurcated expression

The output of the OMGP model is a soft assignment to each of two trends for every observed cell. The original model was fitted with the 2nd latent variable from the latent variable analysis. To find genes that significantly drive this bifurcation, we keep all parameters fixed but change the data to be individual genes expression levels and calculate the data likelihood. In order to get a null distribution to assess significance, we performed the same analysis but with randomly permuted pseudotime-values. This is described in detail in the Computational Supplement.

To measure in which *direction* a gene is involved with the bifurcation, we used correlation between expression and trend assignment. For example, a gene's expression being strongly positively correlated with a trends assignment means it is being upregulated on that bifurcated branch.

### Monocle

The Monocle analysis was performed with version 1.2.0 of the Monocle package, following the steps outlined in the original vignette (17). In brief, the analysis was performed using the size-normalized data (TPM) including all genes expressed in ≥10 cells (11439 genes) with default parameters. The genes used for the ordering of cells were defined by carrying out a differential expression analysis of the time points using the differentialGeneTest embedded in the package. Following the original vignette, genes with q-value <0.01 were selected (7738 genes). The num_paths option was set as 2.

### SCUBA

(https://github.com/gcyuan/SCUBA/tree/2ffa4fe5842dfe88db0207c82088bce0e5b97be7) was run using 3003 genes and provided information about time point. RNAseq_preprocess.m and SCUBA scripts were run according to instructions. SCUBA did not find any bifurcation points. Similarly, using 1000 most informative genes (SCUBA default), or scaling of the data (to aid the sensitivity), did not result in any bifurcation either. Changing the number of was done by the variable ngene_select in RNAseq_preprocess.m. All other variables were kept at default.

### Annotation of cell-surface receptors, cytokines and transcription factors

Genes likely to encode transcription factors, cell-surface receptors or cytokines were found by combining information from KEGG (http://www.genome.jp/kegg/), the Gene Ontology Consortium (http://geneontology.org/, PANTHER (http://www.pantherdb.org/) along with the more specific databases detailed below.

Transcription factors were found by searching the Gene Ontology Consortium database using the following ontology term: *GO:0003700 (sequence-specific DNA binding transcription factor activity)*; KEGG for *ko03000 (Transcription Factors)*; PANTHER for *PC00009 (DNA binding) AND PC00218 (Transcription Factors)*. The presence of genes in the following databases was also used as evidence for transcription factor activity: AnimalTFDB (http://www.bioguo.org/AnimalTFDB/index.php), DBD (http://www.transcriptionfactor.org), TFCat (http://www.tfcat.ca), TFClass (http://tfclass.bioinf.med.uni-goettingen.de/tfclass), UniProbe (http://the_brain.bwh.harvard.edu/uniprobe) and TFcheckpoint (http://www.tfcheckpoint.org).

Cell-surface receptors were found by searching the Gene Ontolotgy Consortium database using the following ontology terms *GO:0004888 (transmembrane signaling receptor activity) OR GO:0008305 (integrin complex)) AND NOT (GO:0004984 (olfactory receptor activity) OR GO:0008527 (taste receptor activity)*; KEGG for *ko04030 (G-Protein Coupled Receptors) OR ko04050 (Cytokine Receptors) OR ko01020 (Enzyme-linked Receptors)*; PANTHER for *PC00021 (G-Protein Coupled Receptors) OR PC00084 (Cytokine Receptors) OR PC00194 (Enzyme-linked Receptors)*. Annotation of genes as receptors in the ImmPort (https://immport.niaid.nih.gov/), GPCRDB (http://gpcrdb.org/) or IUPHAR (http://www.guidetopharmacology.org/) databases was also used as evidence for receptor functionality.

Cytokines were found by searching the Gene Ontolotgy Consortium database using the following ontology terms *GO:0005125 (cytokine activity)*; KEGG for *ko04052 (Cytokines)*; PANTHER for *PC00083 (Cytokines)*. Annotation of genes as cytokines in ImmPort was also used in this case. Genes were scored according to the number of databases and search results in which they occurred. Scores were weighted according to the strength of evidence provided by each database such that functional annotations supported by manually reviewed experimental evidence were given a higher score than those that were solely computationally generated (Table)

| <u>Annotation source</u> | <u>Score</u> |
| --- | --- |
| <u>KEGG</u> | <u>3</u> |
| <u>Gene Ontology Consortium evidence codes IDA, IPI, IMP, IGI, IEP, ISS, ISO, ISA, ISM, IGC, IBA, IBD, IKR, IRD, RCA, TAS, IC</u> | <u>5</u> |
| <u>Gene Ontology Consortium evidence codes IEA, NAS, ND</u> | <u>1</u> |
| <u>PANTHER</u> | <u>2</u> |
| <u>AnimalTFDB</u> | <u>4</u> |
| <u>DBD</u> | <u>1</u> |
| <u>TFCat, classed as ‘transcription factor’</u> | <u>7</u> |
| <u>TFCat, classed as ‘candidate’</u> | <u>5</u> |
| <u>TFClass</u> | <u>4</u> |
| <u>UniProbe</u> | <u>7</u> |
| <u>TFcheckpoint (if manually reviewed)</u> | <u>6</u> |
| <u>ImmPort</u> | <u>4</u> |
| <u>GPRCDB</u> | <u>2</u> |
| <u>IUPHAR</u> | <u>7</u> |

Genes were assigned as likely cell-surface receptors or cytokines if they had a cumulative score greater than or equal to 5 in that category. Genes were assigned as likely transcription factors if they had a cumulative score greater than or equal to 6 in that category.

### In vivo cell depletion

Cellular depletion in *LysMCre* x *iDTR* mice was performed by intraperitoneal injection of 10ng/g DT (Sigma-Aldrich) in 200µl 0.9% saline (Baxter) at day 3 post-infection. Control mice were given 0.9% saline only.

For B cell depletion, anti-CD20 (Genentech) or isotype control antibody was administered in a single 0.25mg dose via i.p. injection in 200ul 0.9% NaCl (Baxter), 7 days prior to infection.

### Confocal microscopy

Confocal microscopy was performed on 10–20 µm frozen spleen sections. Briefly, splenic tissues were snap frozen in embedding optimal cutting temperature (OCT) medium (Sakura) and stored at −80°C until use. Sections were fixed in ice-cold acetone for 10 minutes prior to labelling with antibodies against B220-PE (clone-RA3-6B2) as well as CD68-Alexa Fluor 594 (clone-FA-11) or SIGN-R1-Alexa Fluor 647 (clone ER-TR9). Antibodies against CD68 were obtained from Biolegend (San Diego, CA), and against SIGN-R1 from BIO-RAD (USA). DAPI was used to aid visualization of white pulp areas. Samples were imaged on a Zeiss 780-NLO laser-scanning confocal microscope (Carl Zeiss Microimaging) and data analysed using Imaris image analysis software, version 8.1.2 (Bitplane). Cells were identified using the spots function in Imaris, with thresholds <10mM and intensities <150. All objects were manually inspected for accuracy before data were plotted and analyzed in GraphPad prism (version 6).

**Fig. S1.**
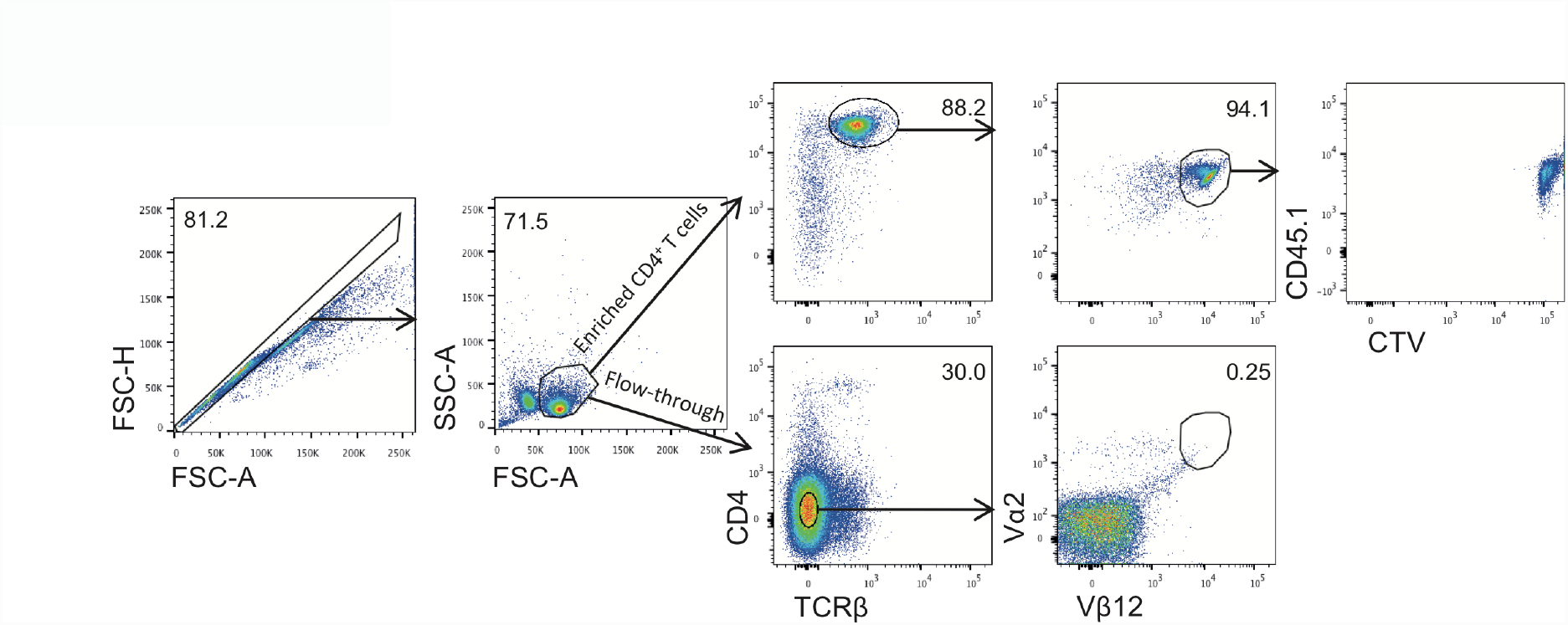
Enrichment of PbTII cells for adoptive transfer. **(A)** CD4^+^ T cells were enriched using positive selection (MACS microbeads) from the spleen of a naive, PbTII x CD45.1 mouse. FACS plots show purity, expression of Vα2 and Vβ12 transgenes, and CellTrace^™^ Violet (CTV) staining of enriched PbTII compared to corresponding flow-through from the enrichment process.

**Fig. S2.**
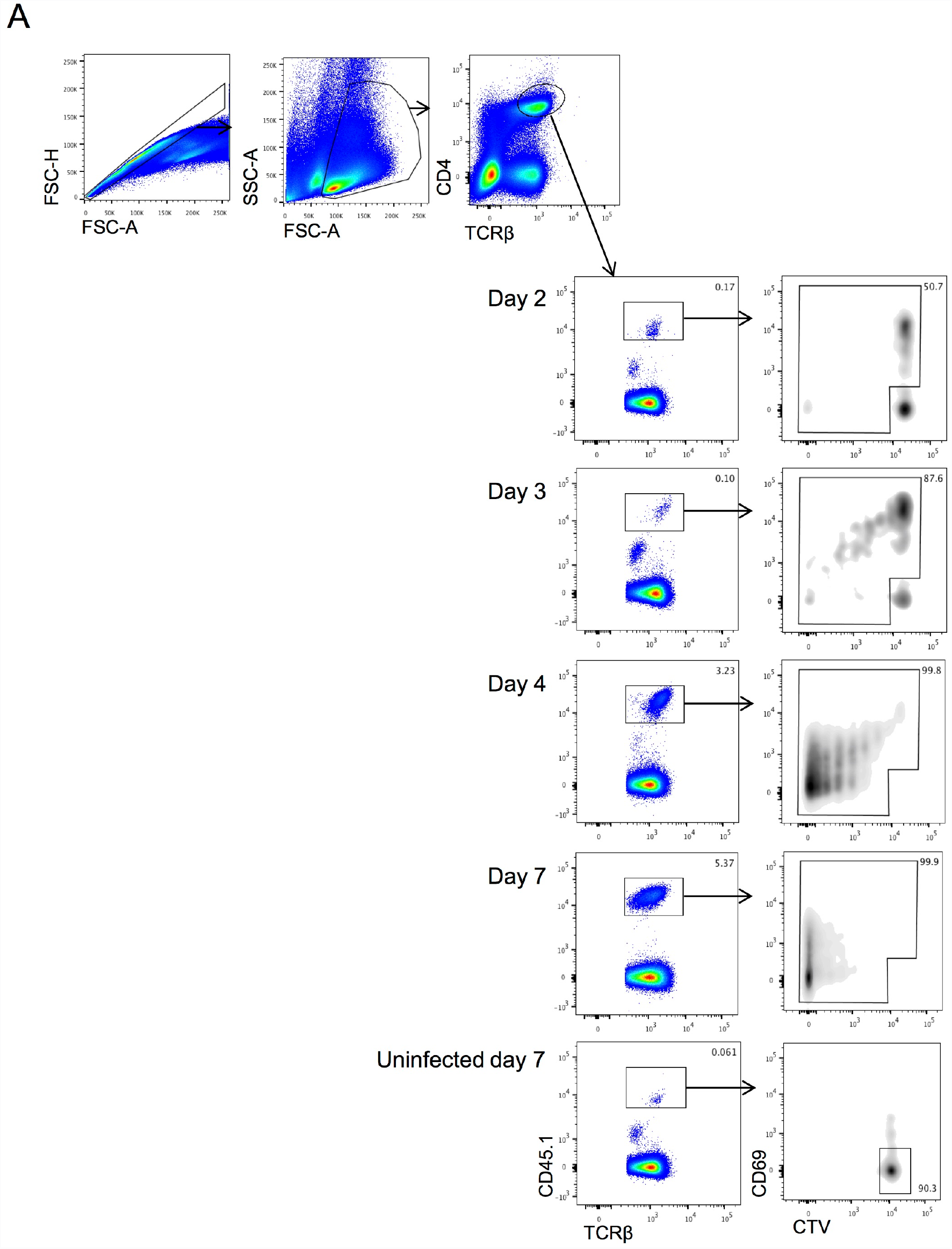
Sorting strategy for PbTII cells. **(A)** PbTII cells (CD4^+^ TCRβ^+^ CD45.1^+^) were adoptively transferred into WT congenic (CD45.2^+^) recipient mice At indicated days, early activated (CD69^+^) and/or proliferated (CTV^lo^) PbTII cells were cell-sorted from spleens of *Plasmodium chabaudi chabaudi* AS infected mice, and naïve PbTII cells (CD69^lo^CTV^hi^) were cell-sorted from the spleens of naïve mice at day 7 post-transfer.

**Fig. S3.**
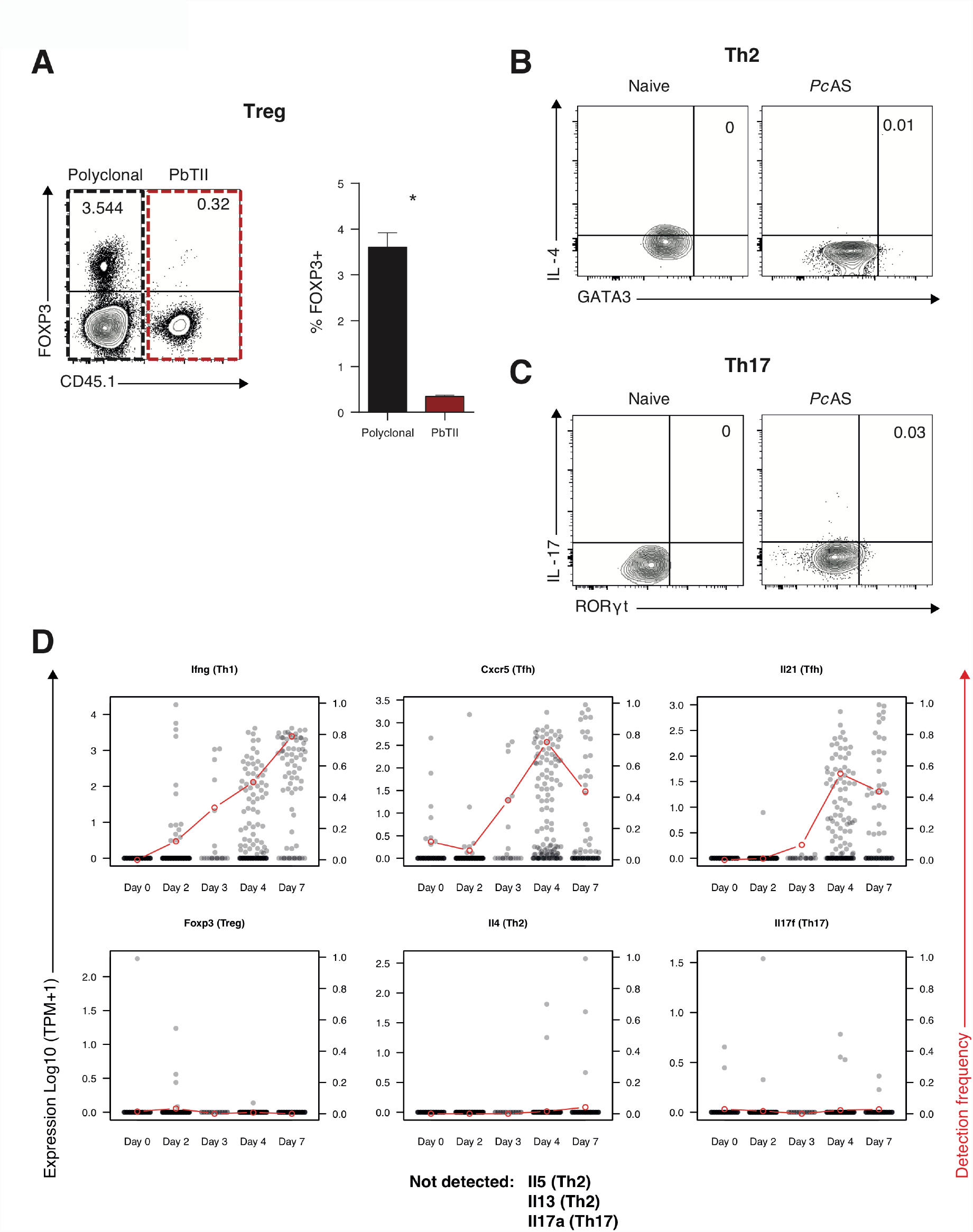
Expression of subset-specific marker genes in PbTII cells. **(A)** Representative FACS plot (gated on CD4^+^ TCRβ^+^ live singlets) and proportion of FOXP3^+^ (Treg) splenic PbTII (10^4^ transferred) (CD45.1^+^; red dashed box) or polyclonal CD4^+^ T (CD45.1^−^; black dashed box) cells from mice (n=6) at day 7 post-infection. **(B-C)** Representative FACS plots (gated on CD45.1^+^ CD4^+^ TCRβ^+^ live singlets) of (B) IL-4^+^GATA3^+^ (Th2) and (C) IL-17^+^RORγt^+^ (Th17) splenic PbTII cells in naive (receiving 10^6^ cells, n=3) or *Pc*AS-infected mice (receiving 10^4^ cells, n=6) at day 7 post-infection. (A-C) Data are representative of two independent experiments. Statistics: Mann-Whitney U test; *p<0.05. **(D)** The mRNA expression of selected subset-specific cytokines and the Treg hallmark transcription factor *Foxp3* in PbTII cells. The red dots and line indicate the fraction of cells in each time point where the particular mRNA was detected.

**Fig. S4.**
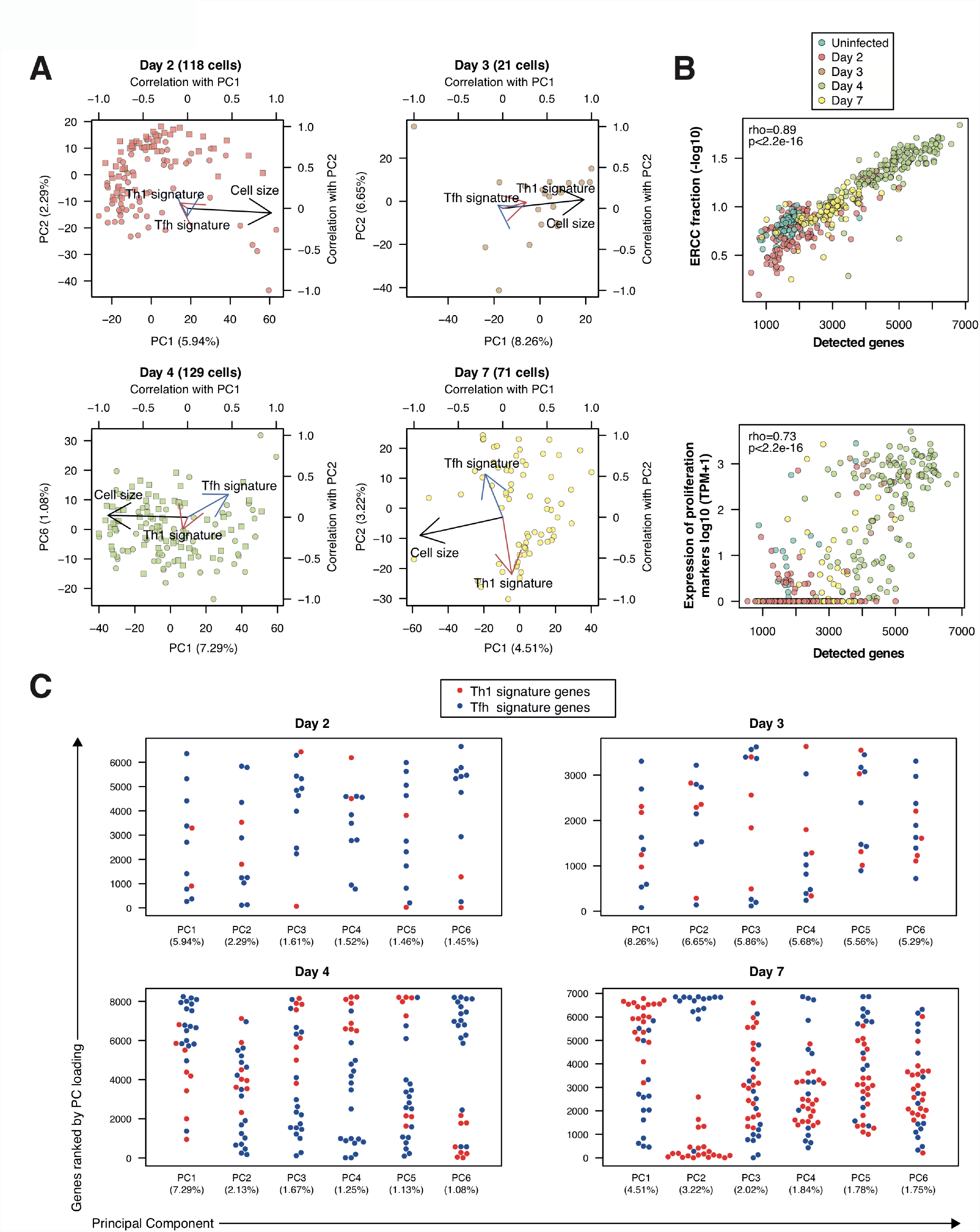
Heterogeneity of activated PbTII cells. **(A)** PCA of single PbTII cells at 2, 3, 4 and 7 days post-infection with *Pc*AS. The PCA was based on all genes expressed at ≥100 TPM in at least 2 cells. The arrows represent the Pearson correlation with PC1 and PC2. Cell size refers to the number of detected genes. “Th1 signature” and “Tfh signature” refer to cumulative expression of top 30 signature genes associated with Th1 and Tfh phenotypes (15). The numbers in parenthesis show proportional contribution of respective PC. **(B)** The relationship of detected cell number with the fraction or reads mapping to ERCC spike-in RNA (top) and with cumulative expression of proliferation markers *Mki67, Mybl2, Bub1, Plk1, Ccne1, Ccnd1* and *Ccnb1* (21) (Figure 4B and S9). **(C)** Ranked loading scores for PC1-PC6 of the Th1 and Tfh signature genes in the PCA shown in (A). The numbers in parenthesis show proportional contribution of respective PC.

**Fig. S5.**
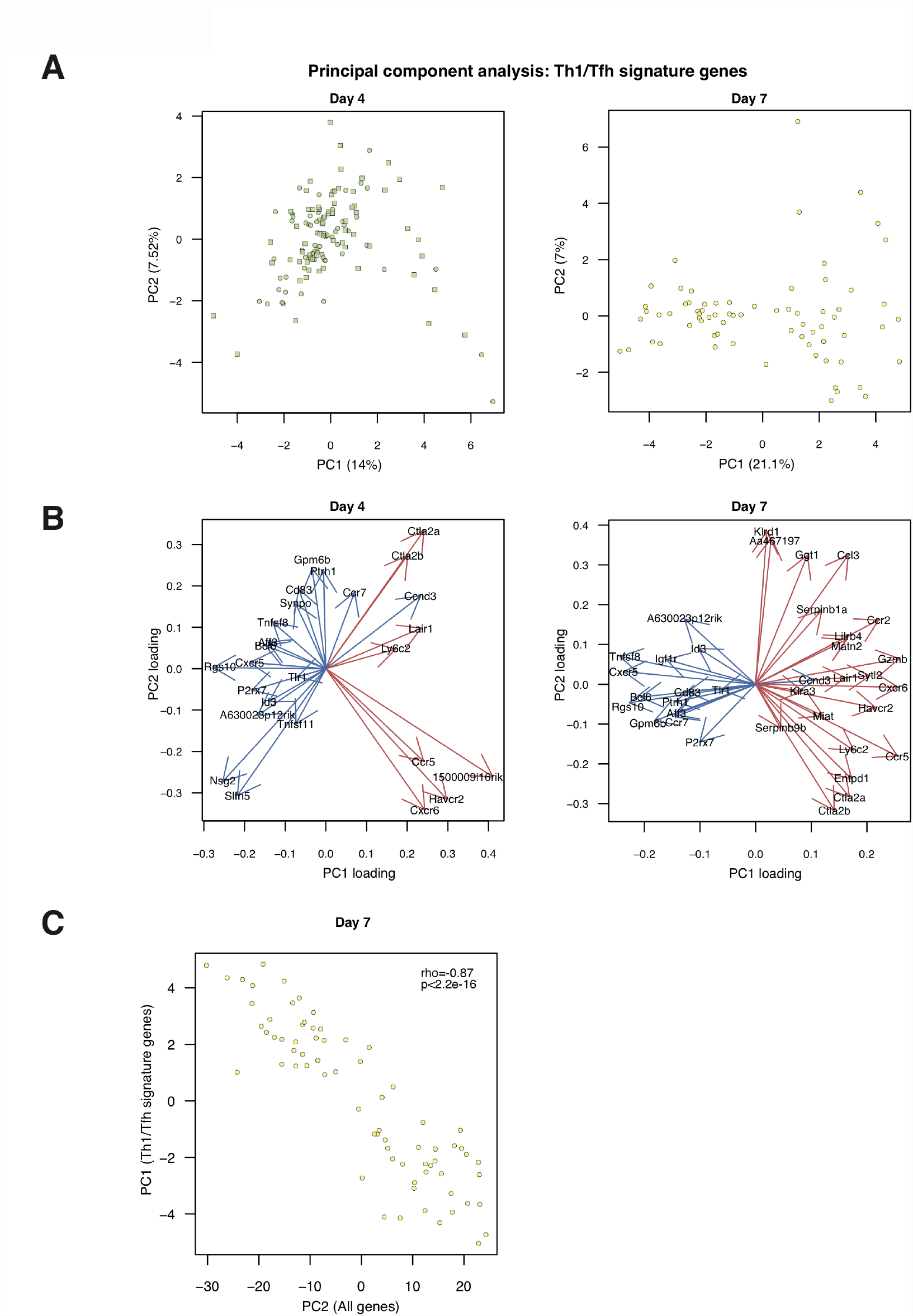
Heterogeneity of Th1/Tfh signature gene expression in activated PbTII cells. (**A**) Principal component analysis of day 4 (left) and day 7 (right) PbTII cells using Th1/Tfh signature genes (15) detected at the level ≥ 100 TPM in at least 2 cells. The numbers in parenthesis show proportional contribution of respective PC. (**B**) The PC1 and PC2 loadings of individual Th1 (red) and Tfh (blue) signature genes in PCA of day 4 and day 7 PbTII cells (A). PC, Principal Component (**C**) The correlation of PC1 from the analysis with the signature genes alone and PC2 of the genome-wide analysis.

**Fig. S6.**
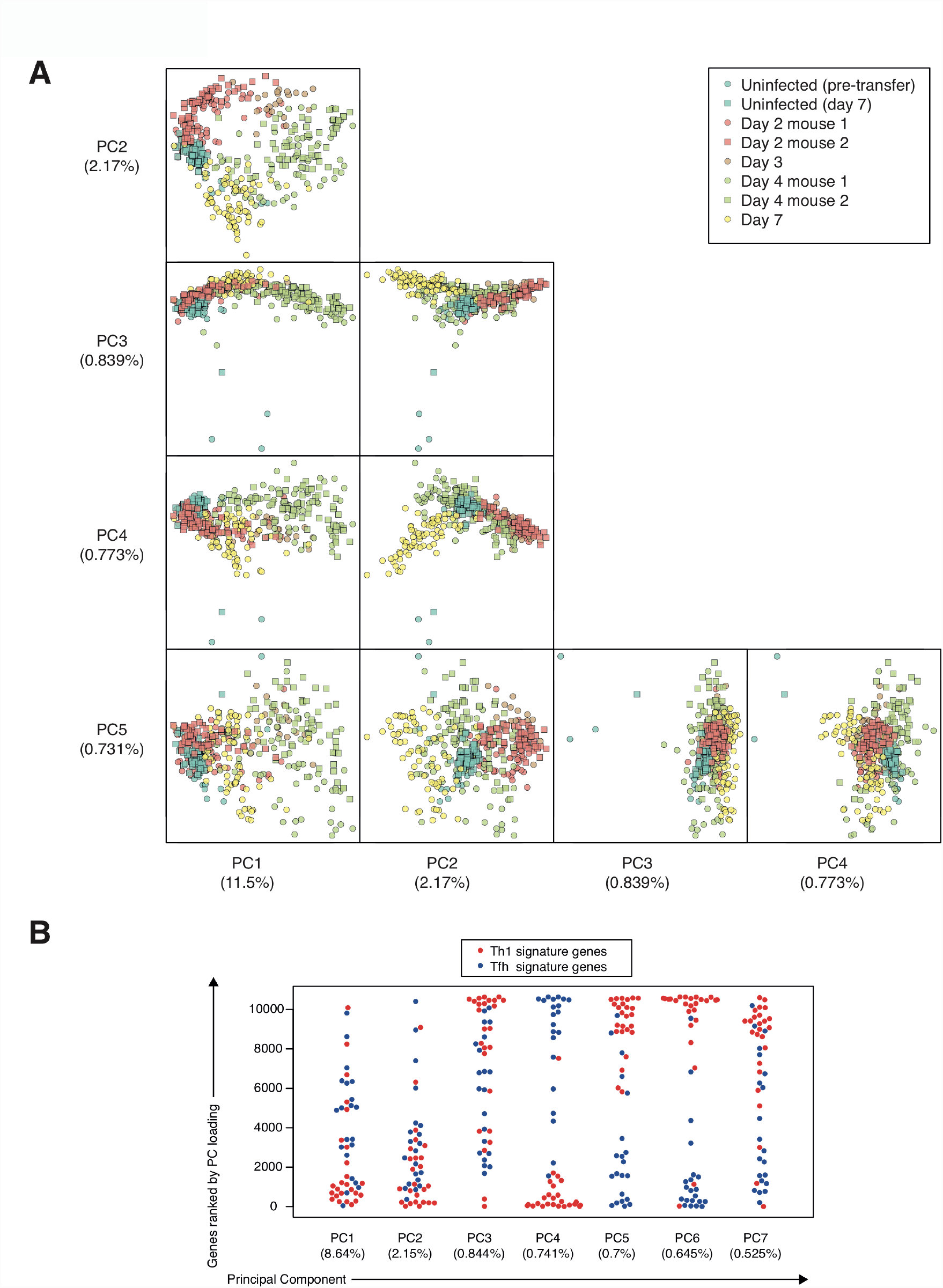
The contribution of Th1 and Tfh signature genes to the overall heterogeneity of the PbTII time series. **(A)** The first five components of the Principal Component Analysis of the entire time series. The numbers in parenthesis show proportional contribution of respective PC. **(B)** The rankings of the Th1 and Tfh signature genes among the loadings of Principal Components 1-7.

**Fig. S7.**
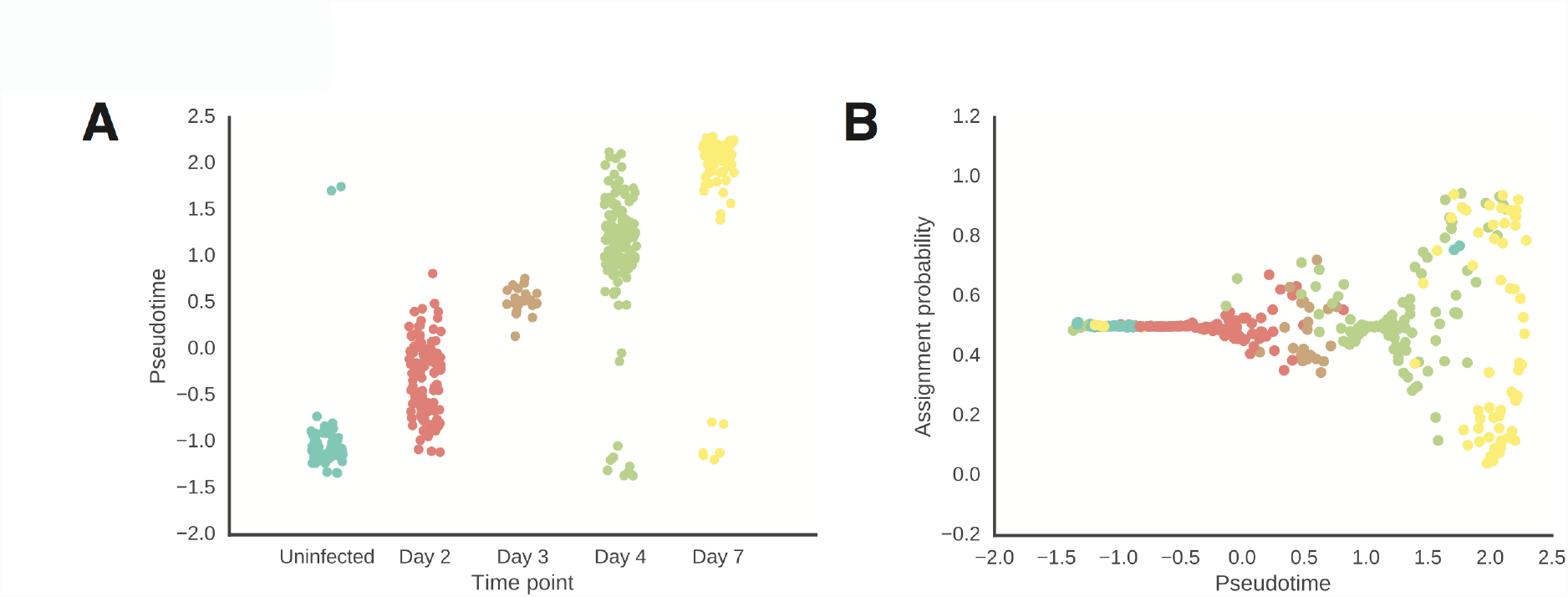
The relationship of pseudotime with time points **(A)** and with the Th1 assignment probability **(B)**.

**Fig. S8.**
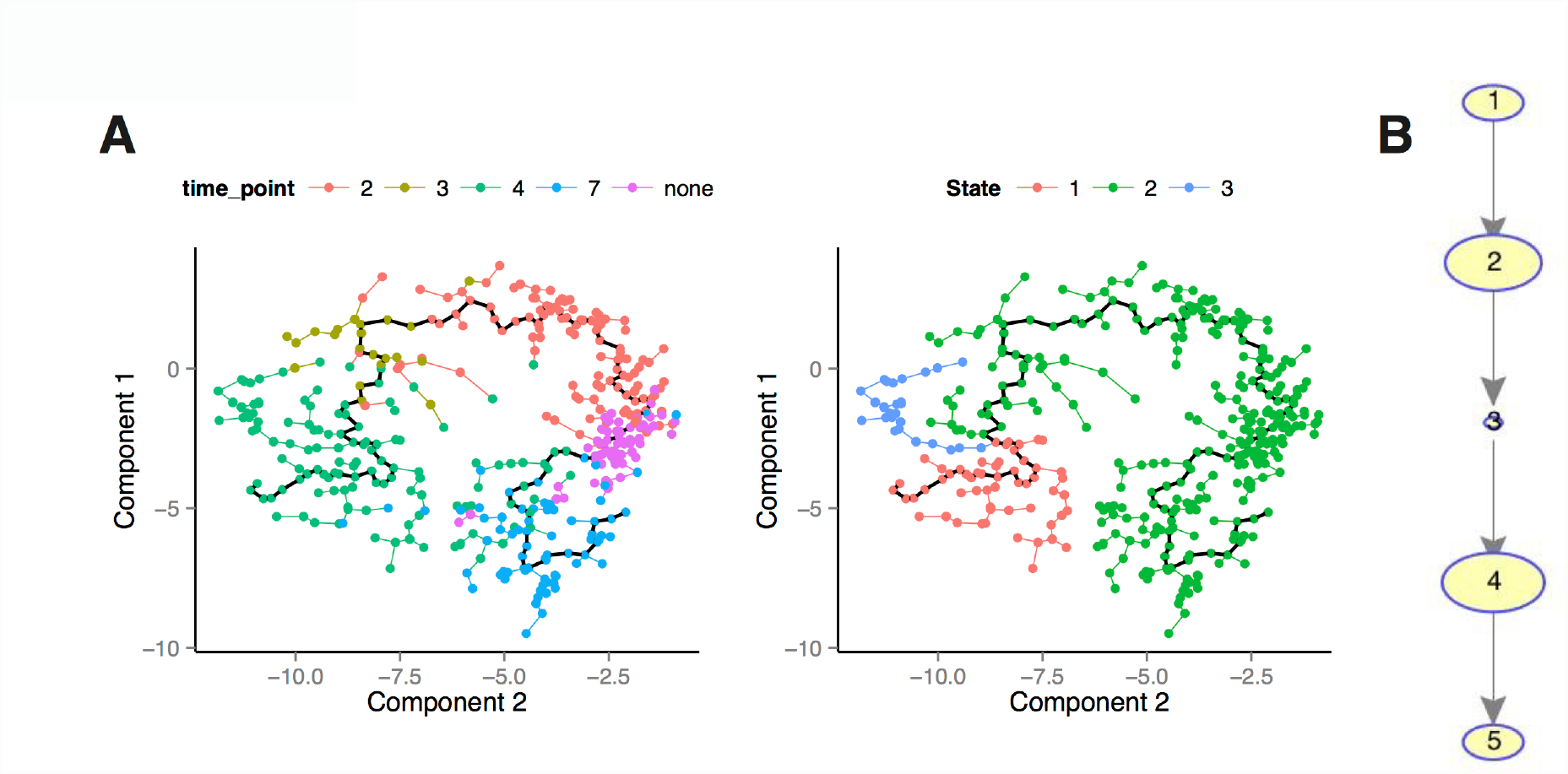
Modelling the data using Monocle and SCUBA. **(A)** Monocle model of the data, coloured by time points (left) and cell states identified by Monocle (right). **(B)** SCUBA bifurcation analysis failed to yield any bifurcating points. Sizes of bubbles are according to number of cells.

**Fig. S9.**
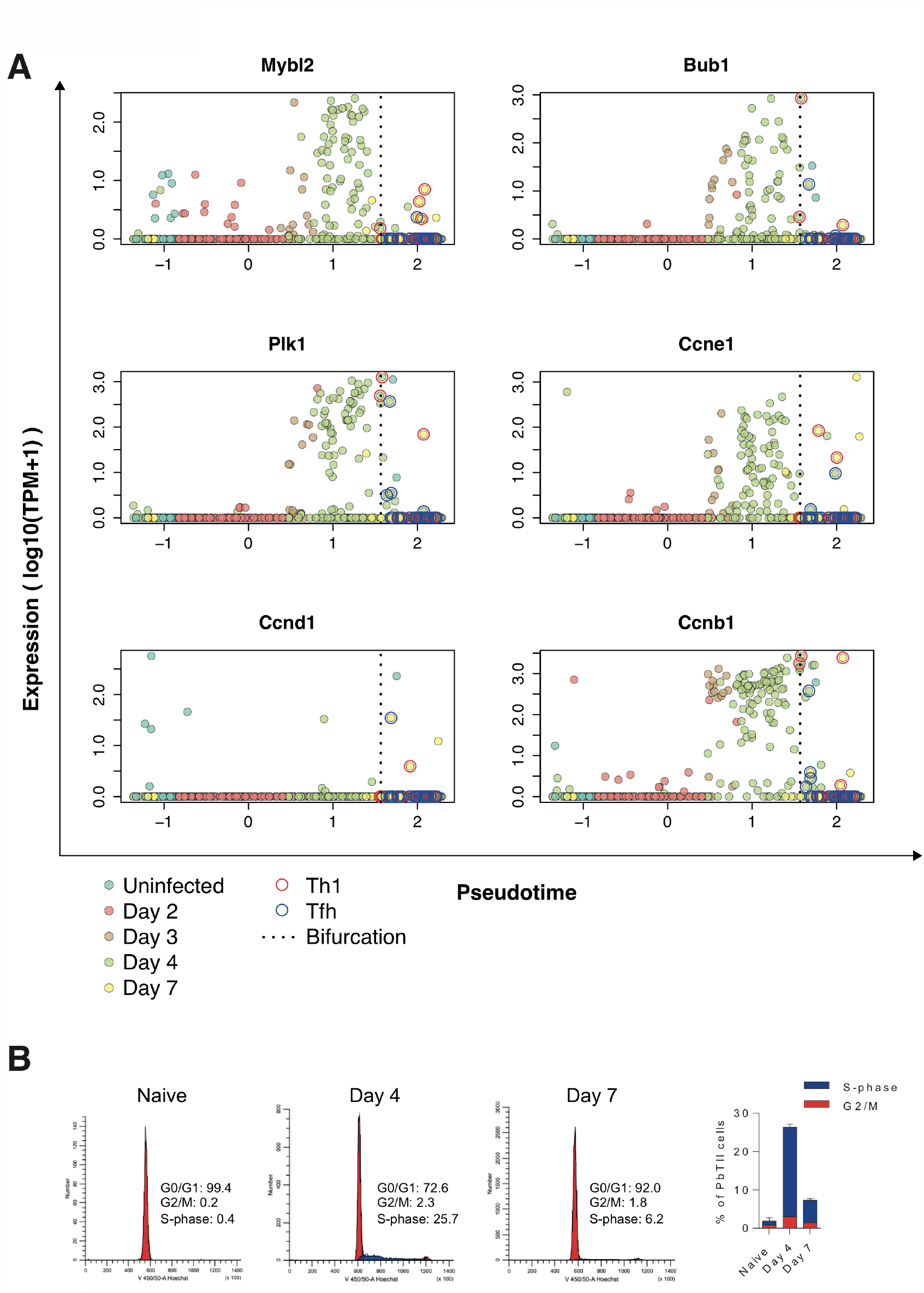
Proliferative burst of activated PbTII cells. **(A)** The expression of established proliferation genes (21) along pseudotime. **(B)** ModFit plots and proportions of PbTII cells in G0/G1, G2/M and S-phase of cell cycle as determined by Hoechst staining.

**Fig. S10.**
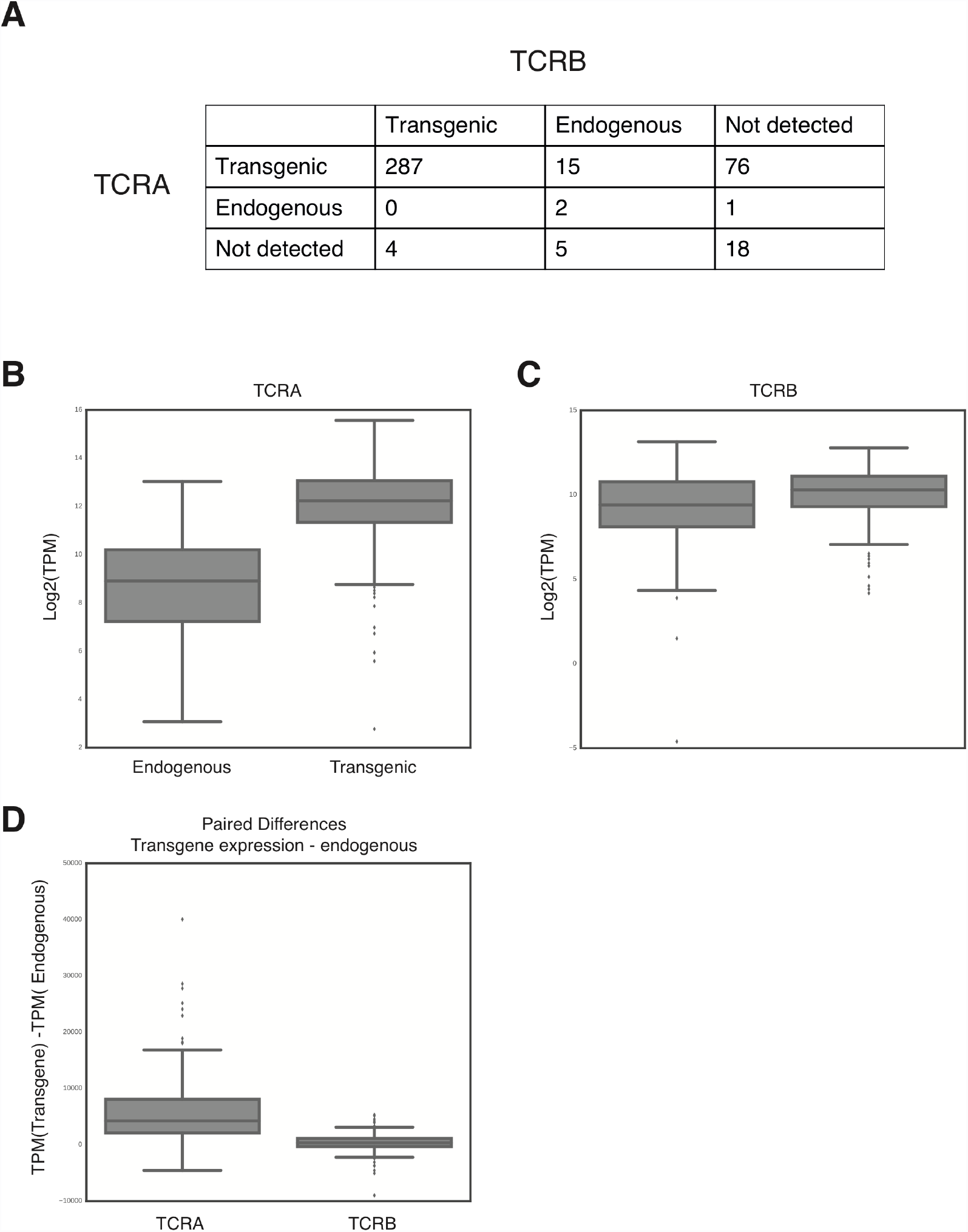
Expression of transgenic and endogenic TCRs. (**A**) Statistics of TCR sequence detection. Numbers correspond to single cells in which the corresponding transcript was detected. (**B**) Expression levels (log2(TPM)) of for the endogenous or transgenic TCRA chains across the entire dataset.

**Fig. S11.**
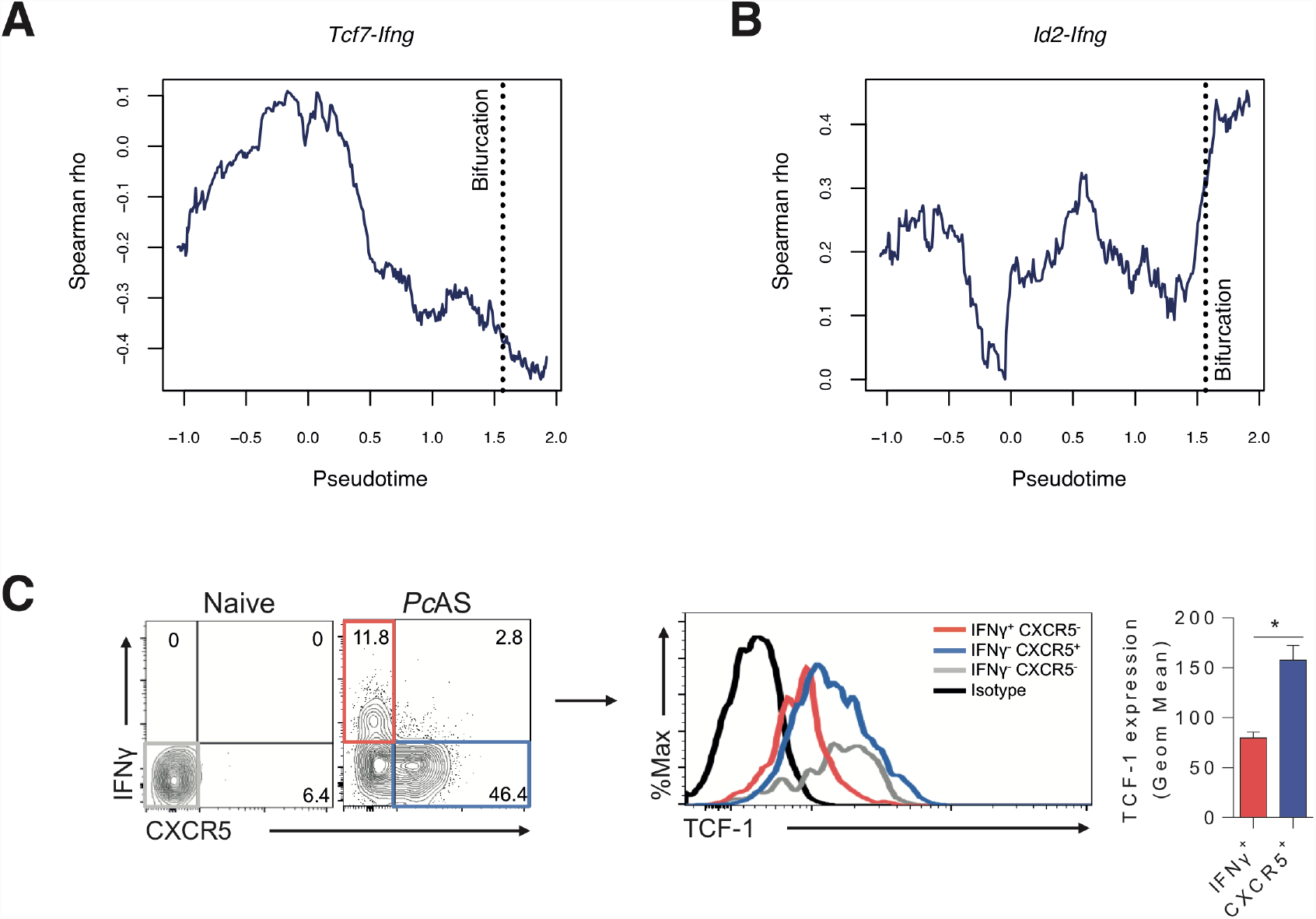
Correlation of expression of *Ifng* with *Tcf7 and Id2* across pseudotime. **(A-B)** The correlation of the expression *Ifng* with *Tcf7* (A) and with *Id2* (B) at single-cell level. Using a rolling window method, Spearman rho was calculated in windows of 100 cells. The pseudotime values are mean values within each window. **(C)** Representative FACS plots showing TCF-1 (gene product of *Tcf7*) expression in CXCR5+ (blue gate) and IFNγ+ (red gate) PbTIIs, compared to naïve PbTIIs (gray) (isotype control shown in black in FACS histogram) at 7 days post-infection. Summary graph shows mean & standard deviations for geometric mean fluorescence intensity of TCF-1 expression in gated PbTII populations (n=4 mice) Statistics: Mann-Whitney U test *p<0.05.

**Fig. S12.**
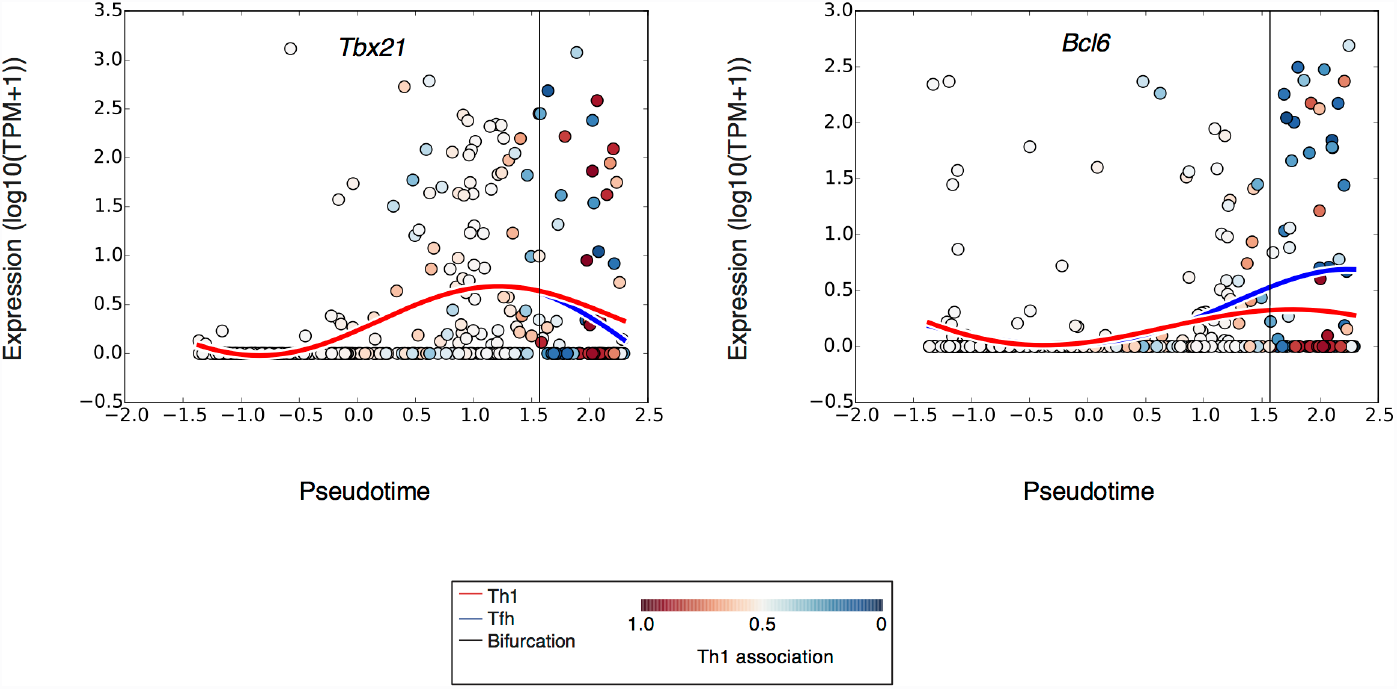
The expression of *Tbx21* (left) and *Bcl6* (right) across pseudotime. The curves represent the Th1 (red) and Tfh (blue) trends when weighing the information from data points according to trend assignment. The color of the data points represents the strength of the relationship with the Th1 trend.

**Fig. S13.**
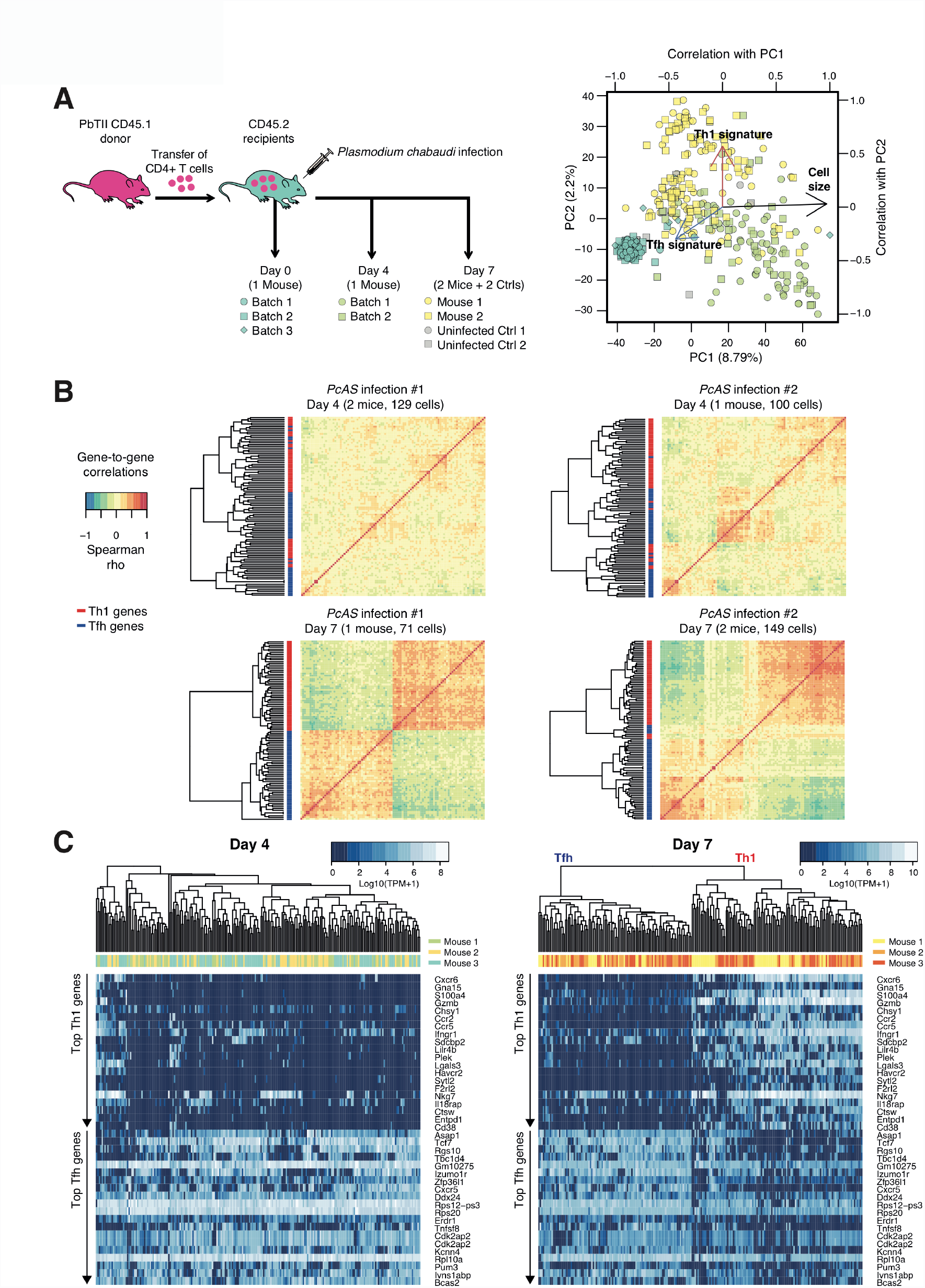
Robustness of top bifurcating genes across experiments. **(A)** Principal Component Analysis of the single cells from the replicate *Pc*AS infection. The single cells were sorted on 96-well plates and cDNA was amplified using the Smart-seq2 protocol (29). The arrows represent the Pearson correlation with PC1 and PC2. Cell size refers to the number of detected genes. “Th1 signature” and “Tfh signature” refer to cumulative expression of genes associated with Th1 or Tfh phenotypes (15). PC, Principal Component. **(B)** The emergence of subset-specific gene patterns at day 7 of infection. For the top bifurcating genes (Fig S5C) pairwise gene-to-gene Spearman correlations were calculated. The rowside colours represent the association of the gene with either Th1 fate (red) or Tfh fate (blue). **(C)** The expression of top 20 Th1 and Tfh associated genes identified using GPfates in single PbTII cells at days 4 and 7. *Cdk2ap2* appears twice because two alternative genomic annotations exist.

**Fig. S14.**
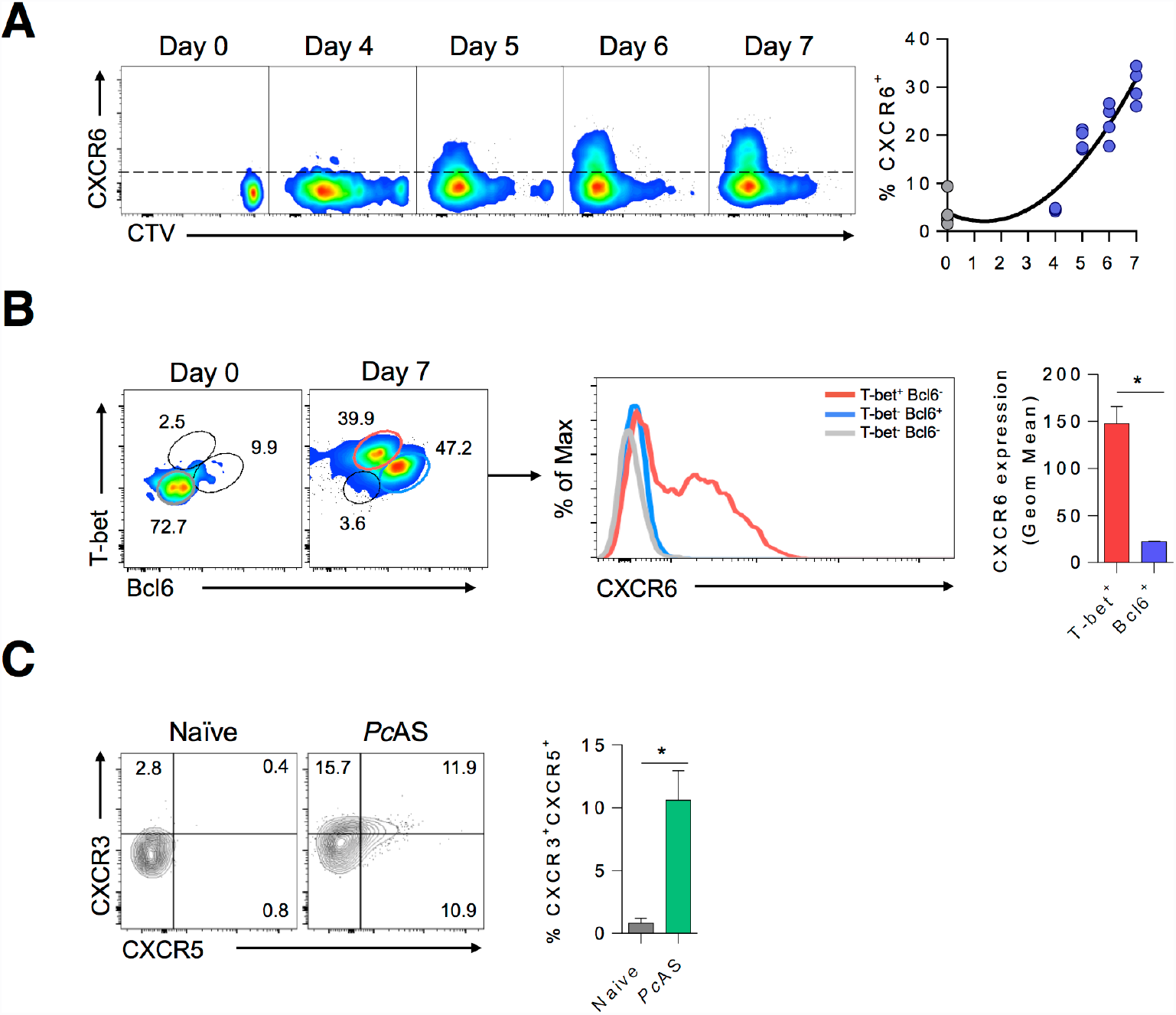
Flow cytometric validation of select marker genes in PbTII cells prior to and after bifurcation. **(A)** Representative FACS plots showing kinetics of CellTraceTM Violet (CTV) dilution and CXCR6 expression, with summary graphs showing % of PbTII cells expressing this (after 106 PbTII cells transferred) in un-infected (Day 0) and *PcAS*-infected mice at indicated days post-infection (n=4 mice/timepoint, with individual mouse data shown in summary graphs; solid line in summary graphs indicates results from third order polynominal regression analysis.) Data are representative of two independent experiments. **(B)** Representative FACS plots showing CXCR6 expression in Tbethi (red gate) and Bcl6hi (blue gate) PbTII cells, compared to naïve PbTIIs (grey) at 7 days post-infection. Summary graph shows mean & standard deviations for geometric mean fluorescence intensity of CXCR6 expression in gated PbTII populations (n=4 mice) Statistics: Mann-Whitney U test *p<0.05. **(C)** Representative FACS plots and proportions of splenic PbTII cells co-expressing CXCR5 and CXCR3 in naive (gray; n=3) or infected mice (green; n=6) at 4 days post-infection with *P.chabaudi chabaudi* AS (*Pc*AS). Results are representative of two independent experiments. Statistics: Mann-Whitney U test *p<0.05.

**Fig. S15.**
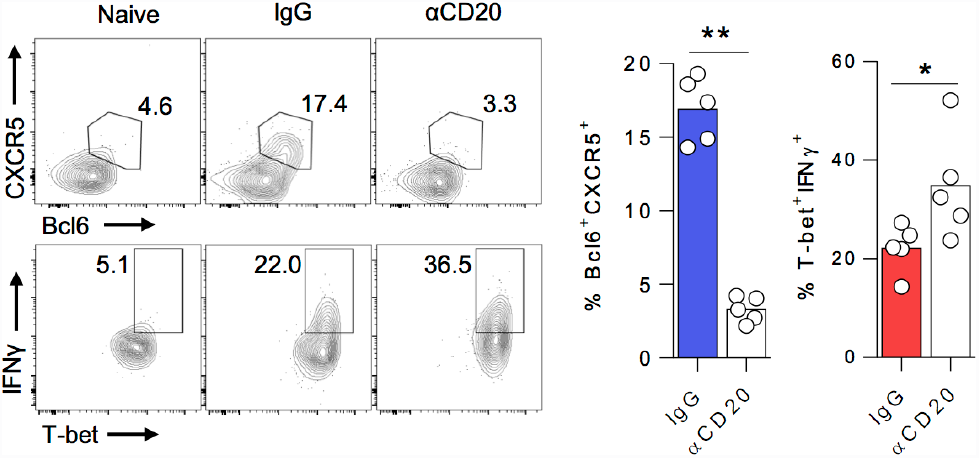
B cells are essential for Tfh responses in PbTII cells during *Pc*AS infection. Representative FACS plots (gated on CD4+ TCRβ+ CD45.1+ live singlets) of splenic PbTII cells, showing proportions exhibiting Tfh (Bcl6+ CXCR5+) and Th1 (Tbet+ IFNγ+) phenotypes in WT mice (receiving 104 PbTII cells), treated with anti-CD20 monoclonal antibodies (0.25mg) to deplete B-cells, or control IgG, and infected for 7 days with *Pc*AS. Individual mice data (n=5) shown in summary graph. Mann-Whitney U test *p<0.05; **p<0.01. Results are representative of two independent experiments.

**Fig. S16.**
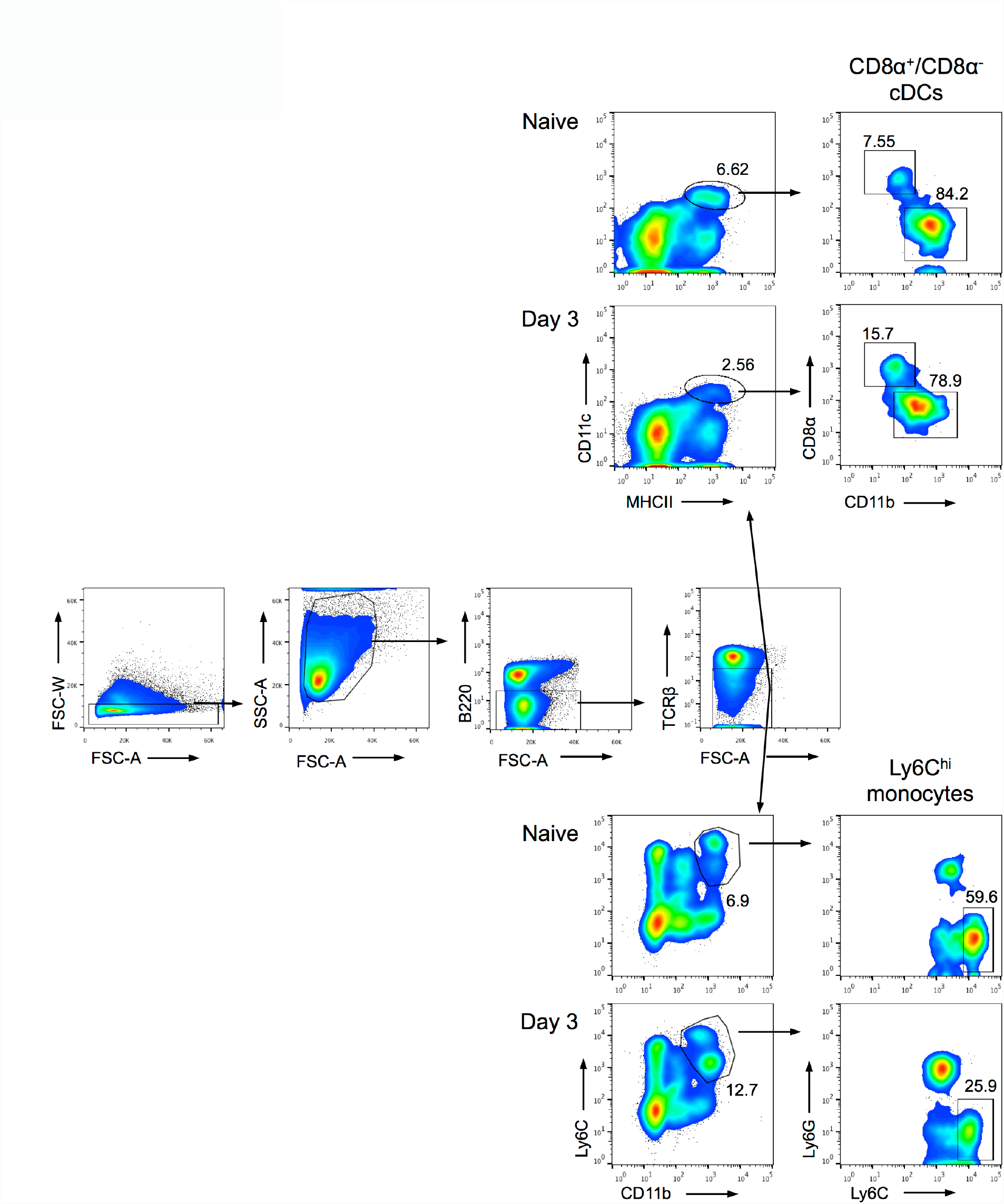
Sorting strategy for myeloid cells. Representative FACS plots showing sorting strategy for CD8α+ and CD11b+ cDC, and Ly6Chi inflammatory monocytes from the spleens of naive and 3-day *Pc*AS-infected mice.

**Fig. S17.**
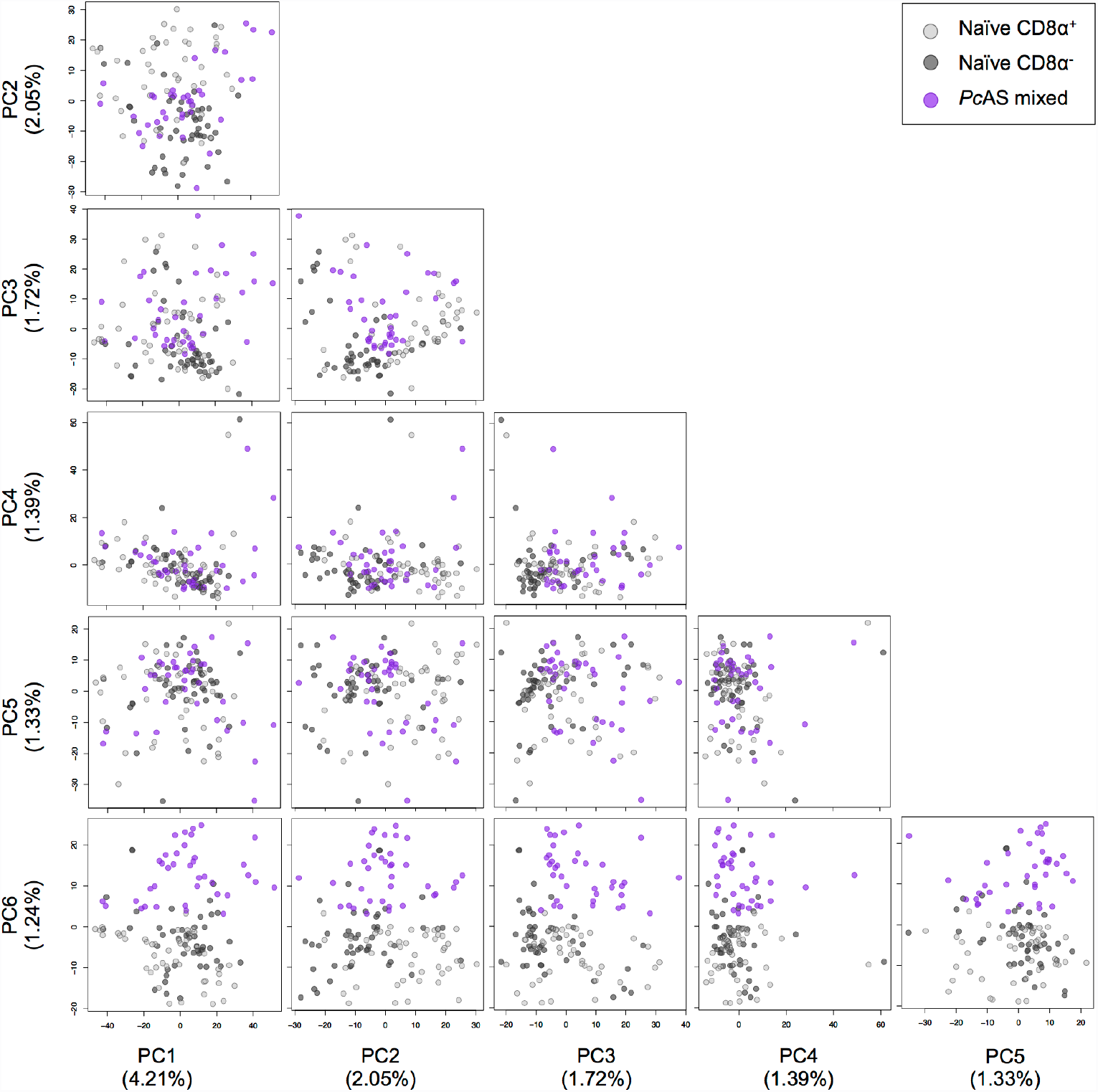
Principal Component Analysis of cDCs from naïve and infected mice. Results of Principal Component (PC) Analysis on scRNAseq mRNA reads (filtered by minimum expression of 100 TPM in at least 2 cells) from 131 single splenic naïve CD8α+ and CD8α− and mixed day 3 *Pc*AS-infected cDC. PC1-PC6 shown. Axis labels show proportional contribution of respective PC.

**Fig. S18.**
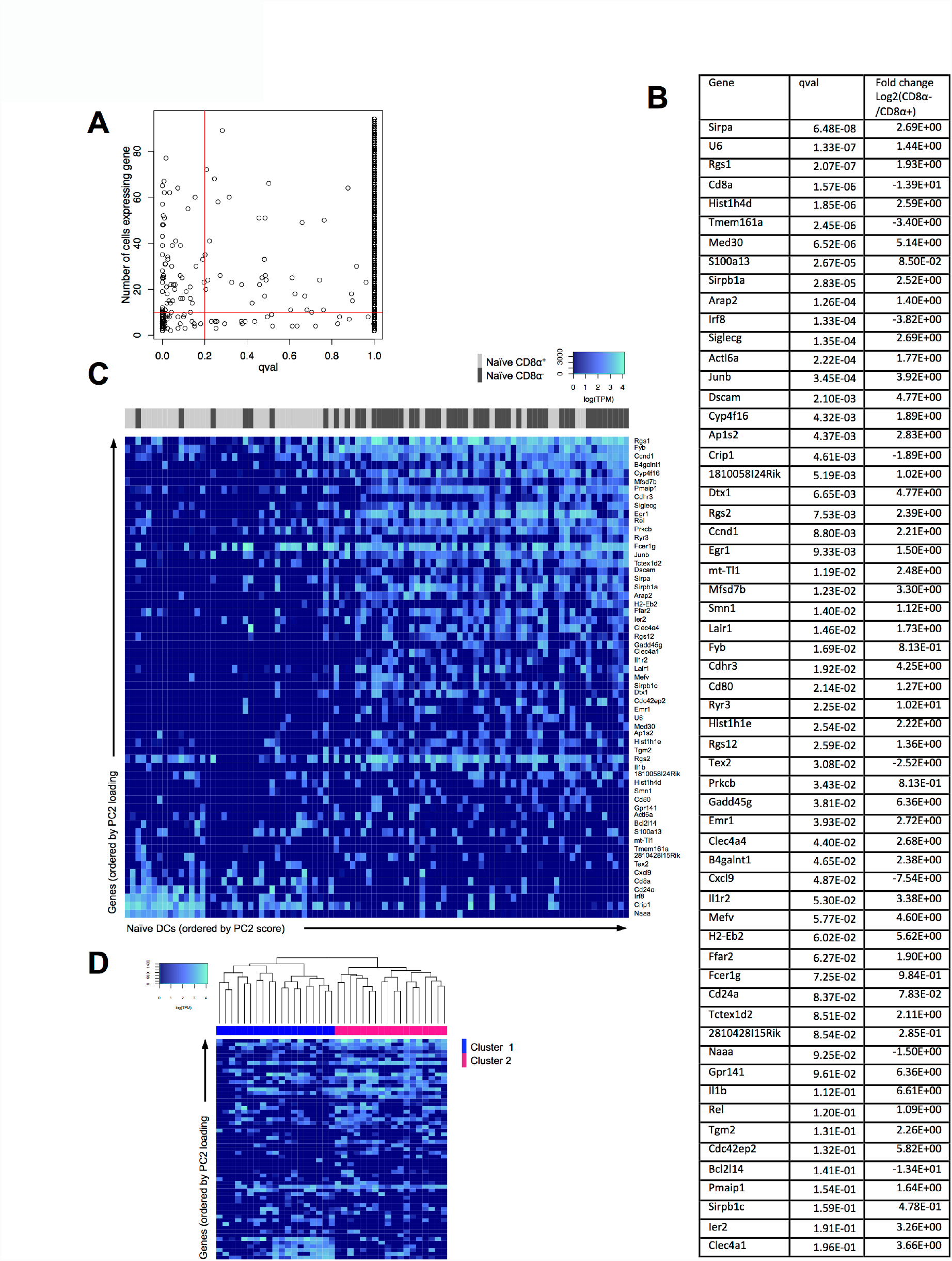
Differential gene expression between single splenic CD8α+ and CD8α− cDCs. **(A)** Results of differential gene expression analysis between naïve splenic CD8α+ and CD8α− cDCs, for all genes expressed in greater than 2 cells. **(B)** Complete list of differentially-expressed genes between naïve CD8α+ and CD8α− cDCs, which were expressed in >10 cells of either subset with a qval <0.2 as determined in (A). **(C)** Heatmap of naïve cDCs ordered by PC2 (Fig. 6A) and expression of genes from (B) ordered by PC2 loading in (Fig 6A). **(D)** Heatmap examining hierarchical clustering of mixed CD8α+ and CD8α− CD11b+ day 3-infected cDCs (cell-sorted and mixed at a ratio of 50:50 prior to scRNAseq) using differentially expressed genes from (B) ordered by PC2 loading shown in (Fig 6A).

**Fig. S19.**
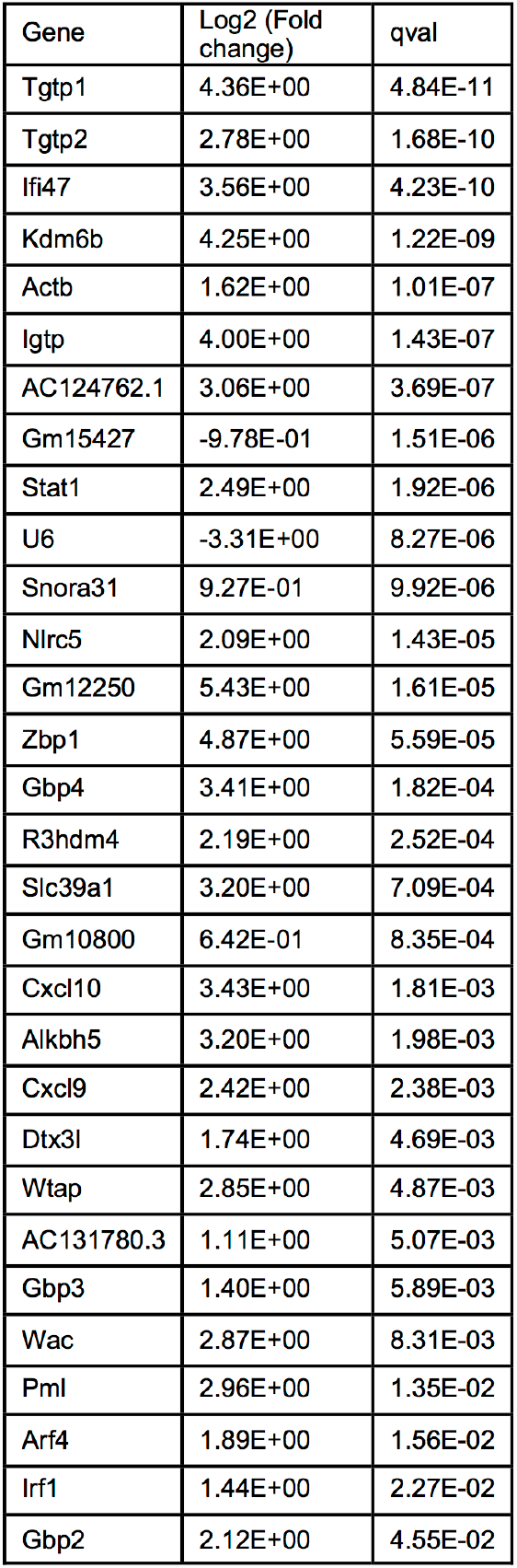
Differentially expressed genes between single naïve and day 3 *Pc*AS-infected cDCs. List of differentially expressed genes, expressed in >10 cells (qval <0.05) between naïve and day 3-infected cDCs. Mean TPM fold-change in gene expression relative to naïve levels.

**Fig. S20.**
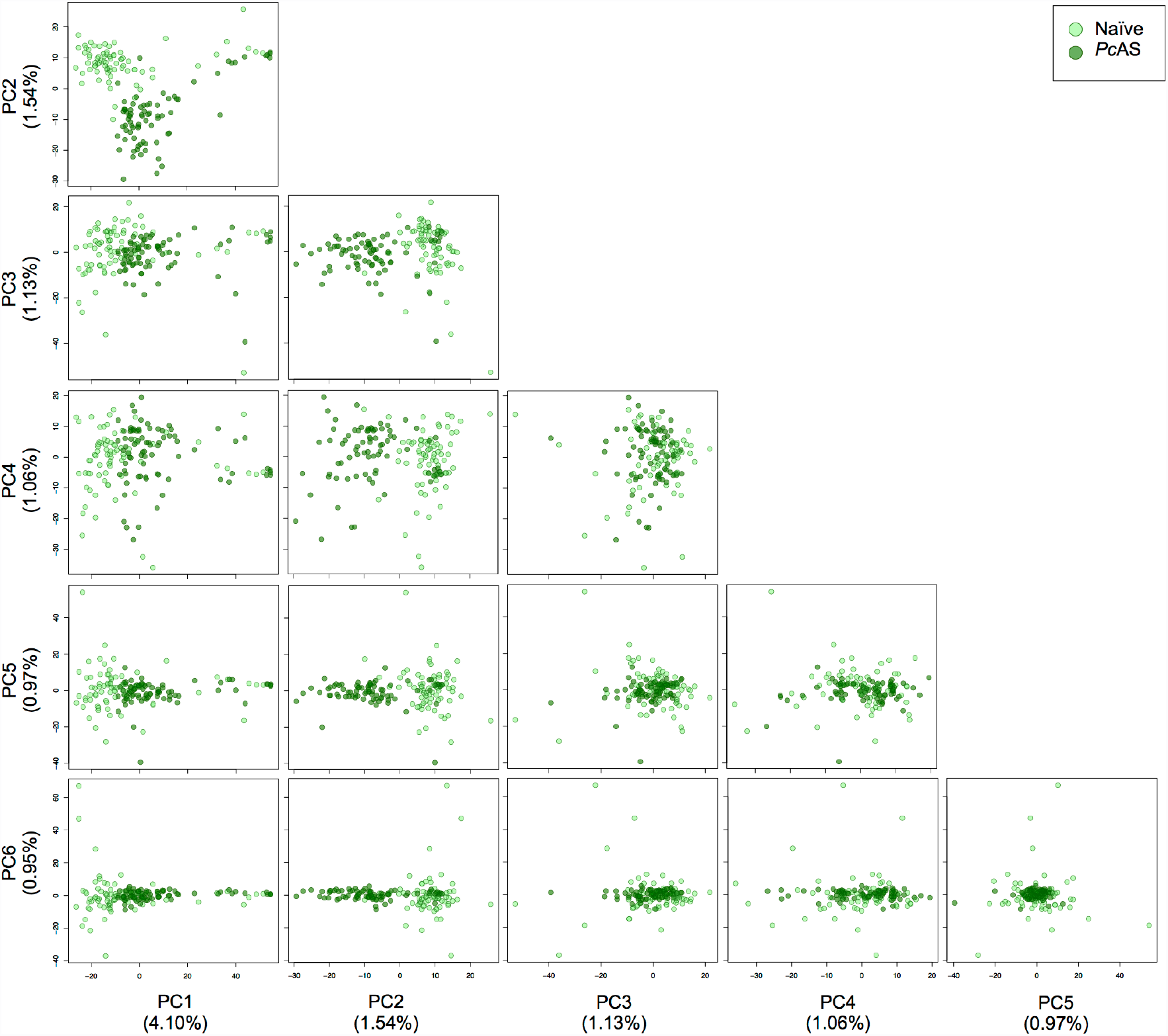
Principal Component Analysis of Ly6Chi monocytes from naïve and infected mice. Results of Principal Component (PC) Analysis using scRNAseq mRNA reads (filtered by minimum expression of 100 TPM in at least 2 cells) of 154 single splenic Ly6Chi monocytes from naïve and infected mice. PC1-PC6 shown. Axis labels show proportional contribution of respective PC.

**Fig. S21.**
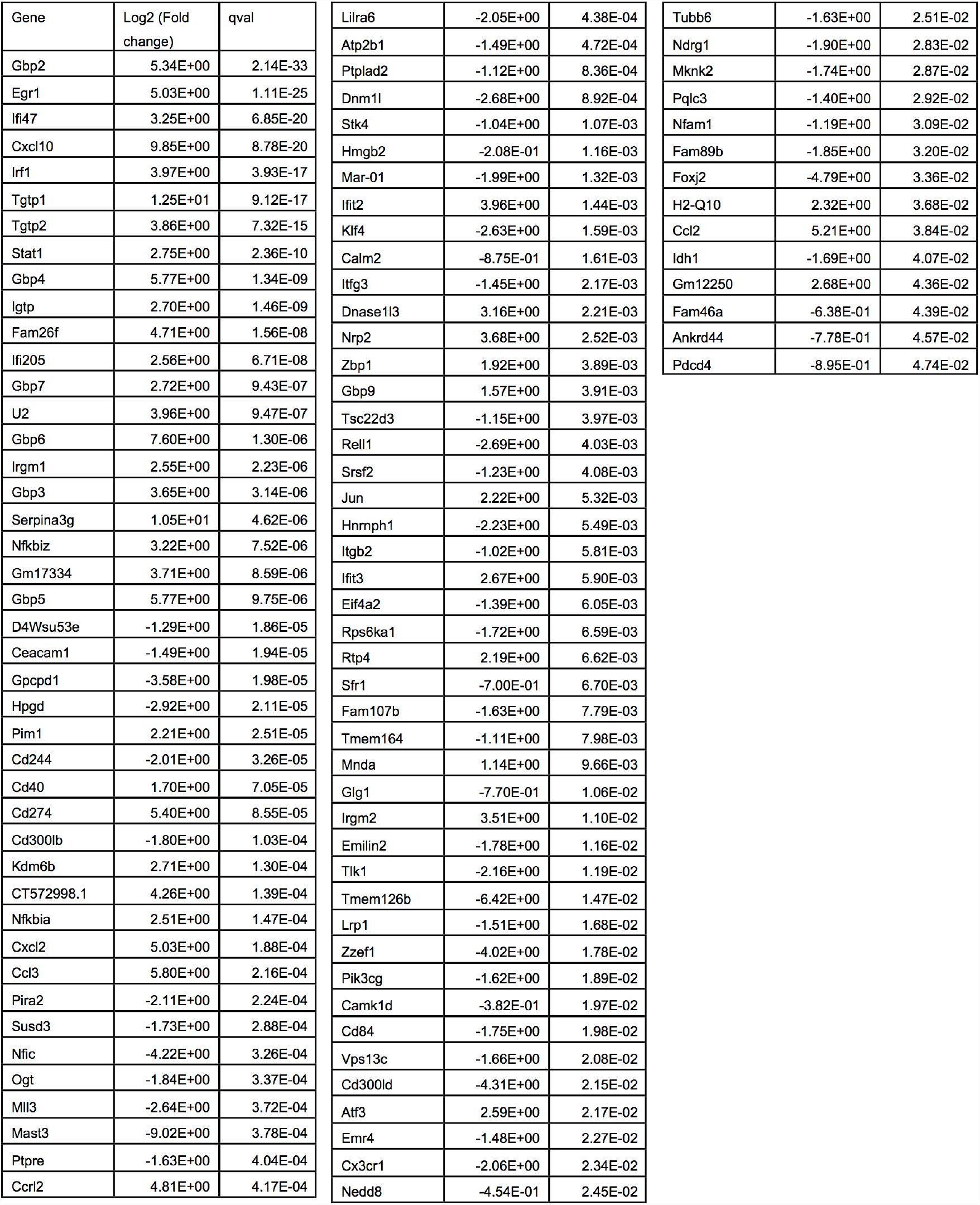
Differentially expressed genes between single Ly6Chi monocytes from naïve and day 3 *Pc*AS-infected mice. List of differentially expressed genes, expressed in >10 cells (qval <0.05) between Ly6Chi monocytes from naïve and day 3-infected mice. Mean TPM fold-change in gene expression relative to naïve levels.

**Fig. S22.**
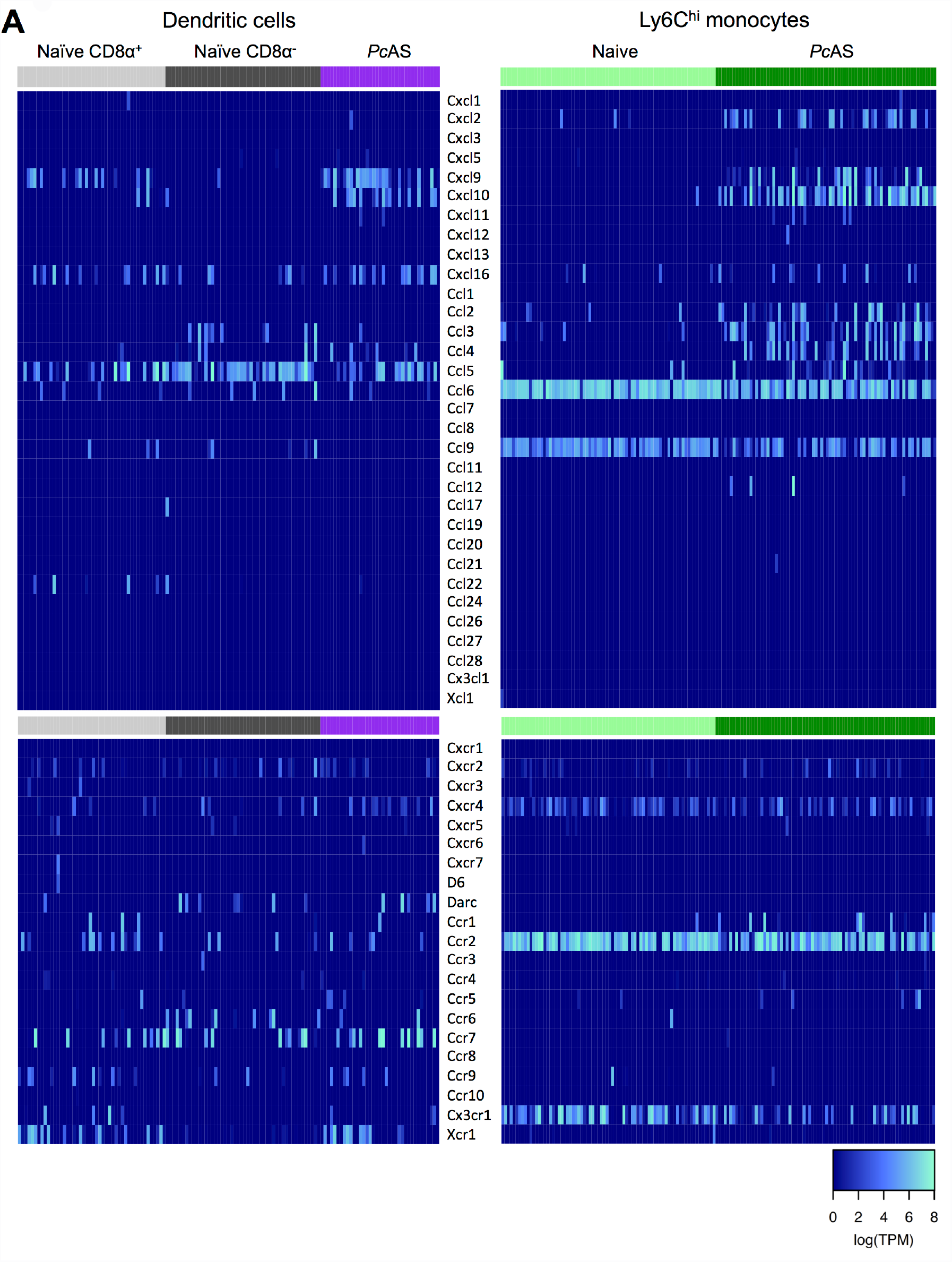
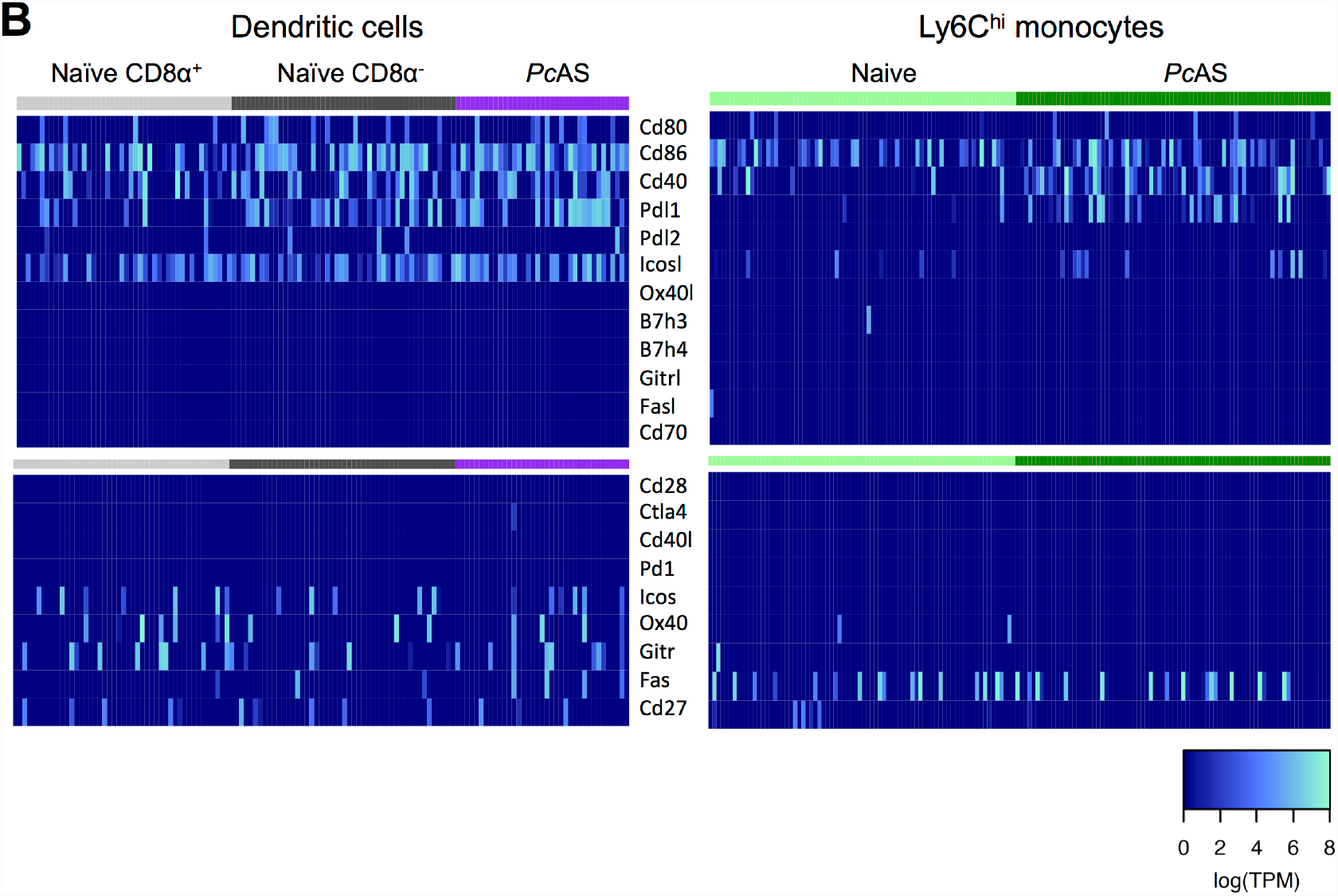
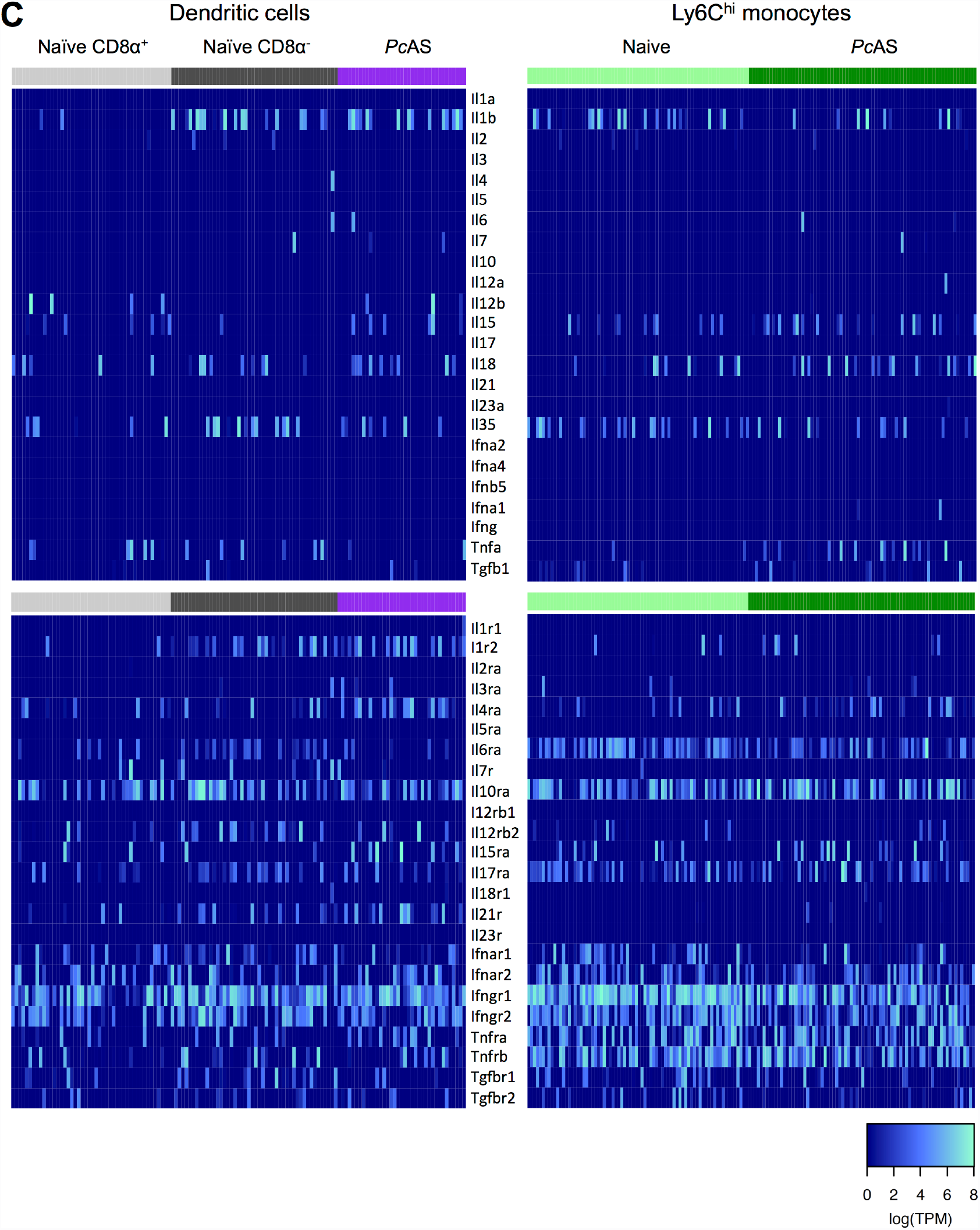
Expression of immune signalling genes by cDCs and monocytes. **(A-C)** Heatmaps showing normalised mRNA expression of select **(A)** chemokines, **(B)** costimulatory molecules and **(C)** cytokines and respective receptors (rows) by single splenic cDCs and Ly6Chi monocytes (columns) from naïve or 3-day *Plasmodium chabaudi chabaudi* AS-infected mice.

**Fig. S23.**
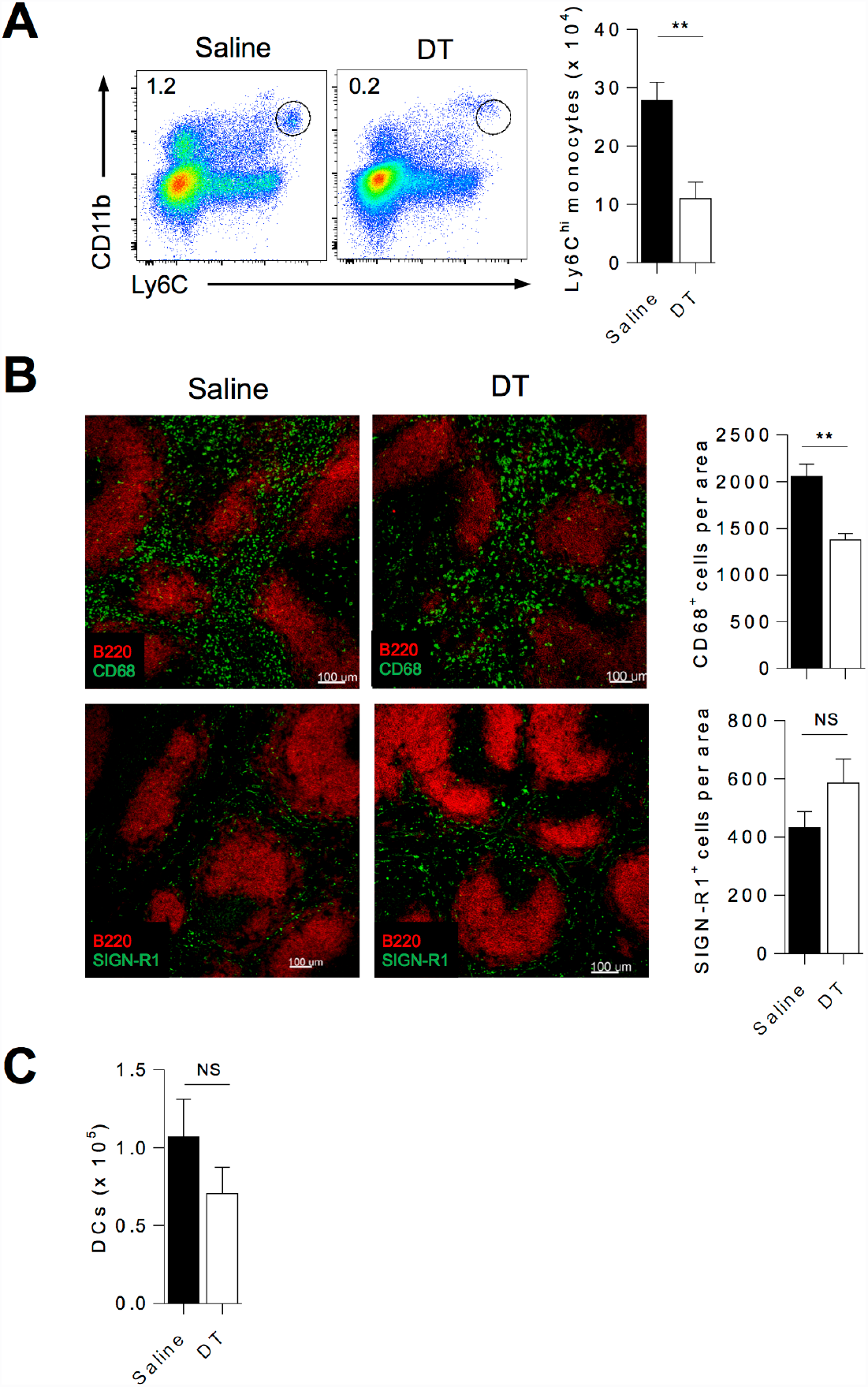
Myeloid cell depletion in LysMCre x iDTR mice. LysMCre x iDTR mice were infected with *Pc*AS, and treated 3 days later with DT (10ng/g intraperitoneal injection) or control saline (n=6 per group). 24 hours later spleens were harvested for cellular compositional analysis: **(A)** Representative FACS plots enumerating splenic inflammatory monocytes (Ly6Chi CD11bhi Ly6G-B220-TCRβ-). (**B)** Representative fluorescence micrographs showing spleen tissue sections co-stained for B cells (B220 in red) and macrophages (CD68 (top panel) or SIGN-R1 (bottom panel) in green) and summary graphs of average cell number in three fields of view covering the total cross section of a spleen. **(C)** Flow cytometric enumeration of splenic cDC (CD11chi MHCIIhi B220-TCRβ-).

### Tables S1-S4

**Table S1** The expression data for all genes on day 7, the PCA loadings for PC1-PC10, and functional annotations for the genes (external file). Th1 annotations are based on studies by Hale *et al.* (SMARTA transgenic, day 6 of LCMV infection, CXCR5^−^Ly6c^hi^), Marshall *et al.* (SMARTA transgenic, day 8 of LCMV infection, PSGL1^hi^Ly6c^hi^), Stubbington *et al.* (*In vitro*, day 4) (15, 46, 47). Tfh annotations are based by studies by Hale *et al.* (CXCR5+Ly6c^lo^, Marshall *et al.* (PSGL1^lo^Ly6c^lo^) and Liu *et al.* (Bcl6-RFP reporter, KLH immunization, CXCR5^+^Bcl6^hi^) (38). Th2 and Th17 annotations are based on Stubbington *et al*.. Annotations for genes associated with exhausted CD4^+^ T cell phenotype are based on Crawford *et al*. (Day 30 of LCMV infection, genes upregulated in exhausted cells but not in memory cells) (51).

**Table S2** TraCeR detection statistics for T cell receptor sequences in single-cell RNA-seq data from the first set of experiments, performed using the C1 platform (external file).

**Table S3** TraCeR detection statistics for T cell receptor sequences in single-cell RNA-seq data from the second set of experiments, performed using the Smart-seq2 platform (external file).

**Table S4** Annotation of receptors, cytokines and transcription factors.

## Footnotes

1 https://github.com/SheffieldML/GPy

2 https://github.com/SheffieldML/GPclust

